# Unique genetic bases of repeated life-history divergence associated with high altitude adaptation in *Mimulus* perennials

**DOI:** 10.1101/2025.08.12.669946

**Authors:** Hongfei Chen, Patricia Vallejo Joseph, Jenn M. Coughlan

## Abstract

Understanding evolutionary repeatability is a central question in biology, as it informs how predictably organisms respond to similar selection pressures. However, the extent to which phenotypic repeatability is recapitulated at the genetic level remains unclear, particularly for quantitative traits. The recurrent evolution of similar phenotypes in high altitude plant populations relative to their low altitude counterparts offers an ideal model for testing genetic repeatability, as these habitats are associated with shifts in complex suites of phenotypes. Here, we investigate the modularity and genetic architecture of life-history trait divergence across four independent transitions to high altitude habitats among closely related perennial taxa in the *Mimulus guttatus* species complex. High altitude taxa exhibit largely repeated phenotypic evolution in 40 univariate traits and suites of traits form correlated modules that are highly similar across taxa. Nonetheless, the genetic architecture underlying each trait was largely non-repeatable, a pattern consistent for both quantitative and genetically simple traits. Despite a general lack of overall repeatability, individual QTLs with larger effects and those that were associated with multiple traits were more likely to be repeatable than smaller-effect or single-trait associated loci. These findings suggest that evolution may follow distinct genetic pathways while repeatedly converging on functionally integrated trait modules. Additionally, although there may be several genetic routes to the same phenotypic outcomes, aspects of genetic architecture can influence the most likely genetic routes taken. Overall, our results provide insights into adaptation to high altitude environments and also advance our understanding of evolutionary repeatability of complex traits.

## Introduction

The remarkable diversity of phenotypes across the tree of life has long captivated evolutionary biologists, yet perhaps even more remarkable are examples in which organisms independently evolve the same (or similar) phenotypes in response to similar selective pressures (herein termed ‘repeated evolution’; Cerca 2023). Repeated evolution not only provides strong evidence of adaptation (Endler 1986; Harvey and Pagel 1991), it can fundamentally inform our understanding of the evolutionary process. For example, repeated evolution allows researchers to gain insights into the predictability and constraint of evolution, the relative roles of drift versus natural selection in population trajectories, as well as how much of the genome can be leveraged to shape particular phenotypes (i.e. a phenotype’s mutational target size) (Bolnick et al. 2018). While much work has focused on *if* evolution is repeatable at a phenotypic or genetic level (i.e. (Losos 1992; Thompson et al. 1997; Colosimo et al. 2005; Steiner et al. 2009; Rosenblum et al. 2010; Elmer et al. 2014; Herman et al. 2018; Jones et al. 2020; Huang et al. 2025), recent interest in this field has sought to understand the circumstances under which evolution is repeatable.

One likely determinant of evolutionary repeatability at the genetic level is the genetic architecture of the phenotype in question. Genetic architecture describes the number and effect size(s) of loci underlying trait expression, their mode of gene action and interaction (i.e. dominance and epistasis), and the extent to which the same loci control multiple traits (i.e. pleiotropy). A rich body of theory and some recent empirical work has revealed that each of these aspects of genetic architecture may influence genetic repeatability. Quantitative traits may be less repeatable than simple traits particularly when selection at each individual locus is weak, as multiple genetic pathways can lead to similar phenotypic outcomes (Orr 2005; MacPherson and Nuismer 2017; Konečná et al. 2022; Poore et al. 2023). Adaptations governed by recessive mutations should be less likely to be involved in repeated evolution, as they are predicted to be commonly lost by drift except under strong assortative mating (i.e. Haldane’s Sieve; Haldane 1927; Orr 2010; Ronfort and Glemin 2013; Wessinger and Kelly 2018). The effect of epistasis on repeated evolution is much less well studied; on one hand, the context-dependency of an adaptive mutation may hamper its ability to evolve (Natarajan et al. 2016), on the other hand, epistasis may constrain the mutational path that evolution can take, leading to increased repeatability (Fisher et al. 2019; Stern et al. 2022) . The role of pleiotropy in facilitating or constraining repeated evolution is also complex. While some early theory and empirical work hypothesized that the extent of pleiotropy should constrain repeated evolution, as pleiotropy increases the likelihood of disrupting other phenotypes (Chevin et al. 2010; Yang et al. 2019), recent empirical work shows that more pleiotropic loci may be more likely to be involved in repeated evolution (Rennison and Peichel 2022; Whiting et al. 2024). This discrepancy may be due, in part, to the relationship between pleiotropy and natural selection. For example, classic quantitative genetic theory predicts that trait correlations can facilitate evolution when the direction of correlation is the same as the direction of selection (i.e. synergistic pleiotropy of ‘trade-ups’, Schluter 1996; Lovell et al. 2013; Burmeister and Turner 2020), but constrain evolution when trait correlations oppose the direction of selection (i.e. antagonistic pleiotropy or trade-offs (Troth et al. 2018). We therefore might expect that under repeated adaptation, traits governed by synergistic pleiotropy may be more repeatable at the genetic level, while traits governed by antagonistic pleiotropy may be less repeatable at a genetic level. Yet, despite the observation that much of adaptation in the wild is conferred by quantitative traits with complex genetic bases (including varying dominance, epistasis, and genetic correlations potentially caused by pleiotropy; Mackay 2010; Johnston et al. 2022; Mackay and Anholt 2024), empirical studies testing these hypotheses are only beginning to emerge.

Moreover, although much work to date has examined repeated evolution in the context of univariate traits, much of adaptation involves complex suites of traits that are either co-selected (i.e. phenotypic integration), co-vary due to persistent genetic correlations (i.e. phenotypic modularity), or both. A classic example is floral divergence that accompanies evolution of a novel pollinator, wherein multiple aspects of floral morphology and biochemistry may evolve in concert, but their evolution at a genetic level involves sets of highly correlated phenotypes that form distinct modules (Diggle 2014; Wessinger and Kelly 2018; Liao et al. 2022; Chen et al. 2025). Creating complex phenotypes through semi-independent modules has been predicted to promote rapid evolution, allowing organisms to fine-tune specific modules to selection (Schlosser 2004; Wagner et al. 2007; Klingenberg 2008; Diggle 2014; Melo et al. 2016), suggesting that highly modular adaptive syndromes may be highly repeatable. Yet, it’s also unclear how such modules can/do change across divergence times (though see McGlothlin et al. 2022), and how modularity influences repeatability at the genetic level.

High altitude adaptation is a classic example of repeated evolution in both plants and animals (Tranquillini 1964; Byars et al. 2007; Scott and Milsom 2007; Alonso-Amelot 2008; Storz et al. 2010; Cheviron and Brumfield 2012; Kooyers et al. 2015; Natarajan et al. 2016; Halbritter et al. 2018; Bohutínská et al. 2021; Coughlan et al. 2021; Wang et al. 2021; Durán-Castillo et al. 2022; Barnbrook et al. 2023; Zhang et al. 2023; Gamba et al. 2024). In plants, high altitude adaptation often involves phenotypic transitions in complex suites of phenological, morphological, biochemical, physiological, and life history phenotypes, such as delayed flowering time, shorter stature, increased production of protective pigmentation, changes to photosynthetic machinery, differences in trichome composition and anti-herbivore defenses, and/or investment in vegetative biomass, including the production of novel, belowground vegetative structures such as belowground storage organs (Gonzalo-Turpin and Hazard 2009; Kammer et al. 2015; Kooyers et al. 2015; Wingler et al. 2015; De Villemereuil et al. 2018; Buckley et al. 2019; Kooyers et al. 2019; Anderson and Wadgymar 2020; Knotek et al. 2020; Bohutínská et al. 2021; Coughlan et al. 2021; Wang et al. 2021; Gamba et al. 2024). These phenotypic shifts are predicted to be underlined by diverse genetic architectures, including highly polygenic (i.e. flowering time, size; Leinonen et al. 2013; Romero Navarro et al. 2017; Yan et al. 2021; Fournier-Level et al. 2022), and relatively simple traits (i.e. pigmentation and trichome production; Holeski et al. 2010; Hendrick et al. 2016; Arteaga et al. 2022), traits that exhibit significant genetic correlations indicative of a potentially pleiotropic basis (i.e. aspects of life history; Mitchell-Olds 1996; Hall et al. 2006; Friedman et al. 2015; Coughlan et al. 2021; Wright et al. 2022), and traits governed by loci with epistatic effects (i.e. rhizomes and trichomes; Lauter et al. 2004; Friedman 2014; Coughlan et al. 2021). High altitude adaptation therefore presents an opportunity for us to understand how genetic architecture, trait modularity, and phenotypic integration influence evolutionary repeatability. Moreover, given that high altitude transitions are common both within species (i.e. via local adaptation) and between species (i.e. via endemism), they present ideal environments to assess repeatability of phenotypic modules and loci across divergence times.

The *Mimulus guttatus* species complex consists of several closely related taxa that inhabit a wide range of altitudes across the western half of North America and exhibit significant variation in life histories, making it an ideal model for studying evolutionary repeatability (synon: *Erythranthe guttata* species complex; Barker et al. 2012; Nesom 2014, but see Wu et al. 2008; Twyford et al. 2015; Lowry et al. 2019; Coughlan et al. 2021). Within the perennials of this group, three are mid to high altitude endemics (*M. tilingii, M. decorus, M. corallinus*; Fig. 1), perennial *M. guttatus* includes both low elevation coastal perennials (Lowry et al. 2008; Hall et al. 2010; Lowry and Willis 2010) and inland perennials that span a broad altitudinal range (Lowry et al. 2008; Friedman and Willis 2013; Coughlan et al. 2021). In this complex, perennials are distinguished by the production of stolons, which are horizontally growing vegetative stems that develop from axillary meristems, root at each node, and remain above ground throughout development. The three high altitude endemic perennial taxa produce both stolons and rhizomes, which are horizontally growing vegetative stems that initially emerge above ground but quickly transition underground during early development. Once underground, these stems undergo radical morphological changes, including chlorosis, reduction of leaves, and increased branching (Fig. 1; Coughlan et al. 2021). By contrast, the low altitude coastal perennial *M. guttatus* lacks rhizomes entirely. High altitude perennial *M. guttatus* occasionally makes rhizomes (Coughlan et al. 2021). Along with rhizome formation, the high altitude taxa display traits characteristic of high altitude environments, such as increased investment in above ground vegetative biomass, earlier flowering time, shorter stolons, and reduced internode length compared to the coastal *M. guttatus* (Fig. 1; Table S1).

**Fig. 1:**
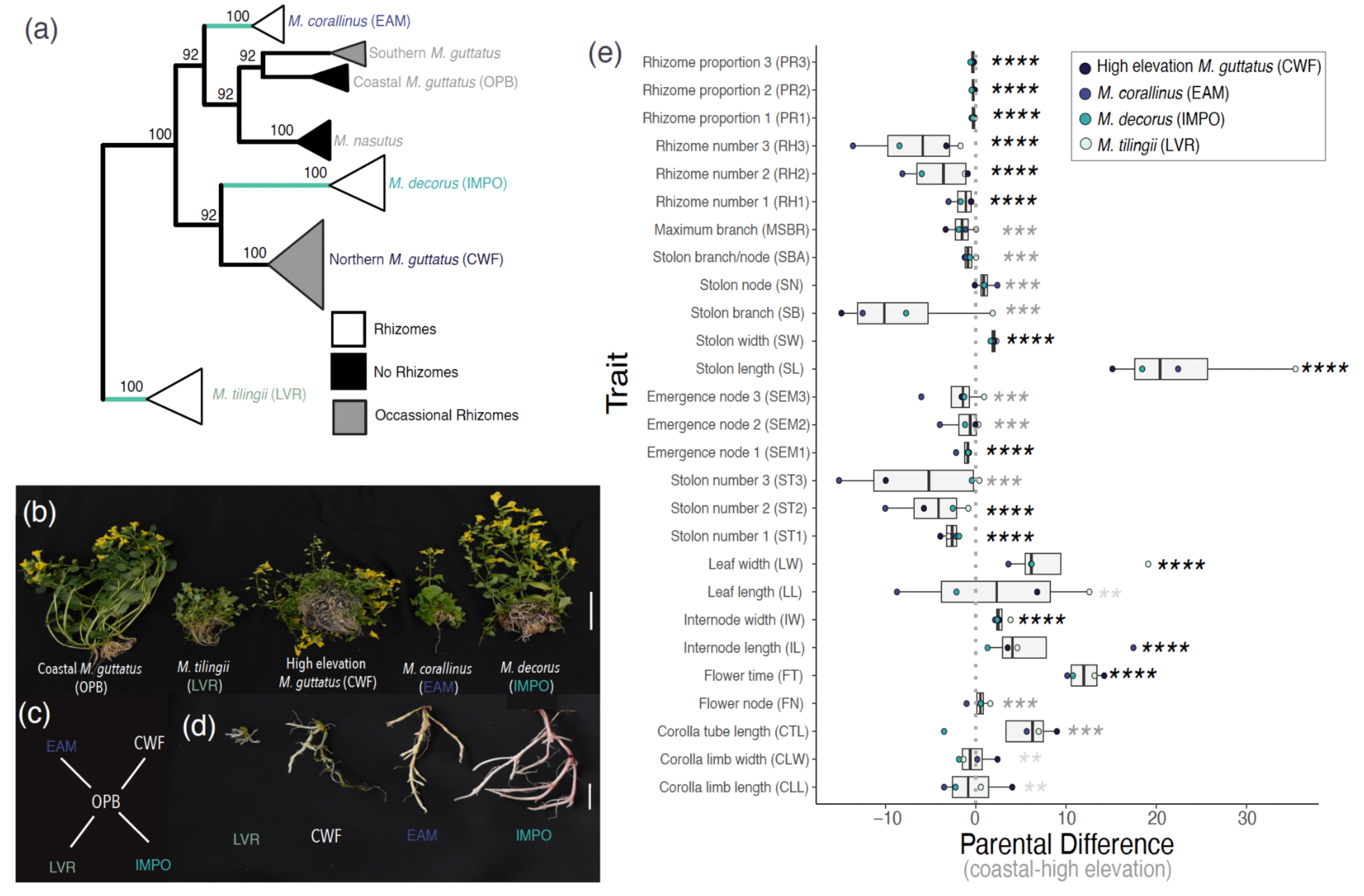
Repeated high elevation adaptation in the *M. guttatus* species complex. (a) Phylogeny indicating the distribution of high elevation endemics (green branches) and rhizomes (sharing of branch tips). Adapted from Ivey et al. 2023 and Coughlan et al. 2021. (b) Photos of the five lines used in all F2 mapping: low-elevation coastal perennial (OPB), *M. tilingii* (LVR), high-elevation inland perennial *M. guttatus* (CWF), *M. corallinus* (EAM), *M. decorus* (IMPO). (c) Crossing design for replicated F2 populations where each high-elevation line was crossed to a common low-elevation line (OPB) and a single F1 individual was self-fertilized to produce an F2 mapping population. (d) Representative photos of rhizomes across the four high-elevation taxa. (e) Parental difference between the low-elevation *M. guttatus* and each high-elevation taxa. Asterisks denote the number of crosses that exhibit a significant parental difference in the same direction.

Here, we investigate the genetic architecture underlying parallel life-history divergence among four independent F2 populations, each formed by crossing a distinct high altitude perennial taxa with a common low altitude, coastal perennial *M. guttatus*. Specifically, we define distinct modules related to morphological and phenological divergence, and analyze the repeatability within and between these modules at the phenotypic and genetic level using quantitative trait loci (QTLs) as well as their dynamics throughout development. Our study aims to address several key questions regarding the genetic architecture of life-history divergence: (1) How repeatable is high altitude adaptation at the level of univariate traits? (2) Do life-history traits form distinct phenotypic modules and how repeatable are these modules? (3) How repeatable is the genetic architecture of trait divergence and how do aspects of the genetic architecture of high altitude associated traits, such as number and effect size of QTLs, influence repeatability? (4) Does the extent of divergence time between high altitude taxa influence the likelihood of repeatability? Our findings provide significant insights into the genetic architecture of life-history divergence among closely related perennial taxa and enhance our understanding of evolutionary repeatability.

## Results

### High-altitude taxa exhibit repeated phenotypic divergence

All four high altitude parental lines differed significantly from a common low altitude, coastal *M. guttatus* line across almost all measured traits, and the direction of differentiation was consistent across high altitude taxa for the majority of these traits (15/29 traits were consistent across all four high altitude lines, and 10 additional traits showed consistent directions of differentiation in ¾ high altitude lines; Fig. 1; Fig. S1; Table S1). High altitude taxa invested more in vegetative biomass and asexual reproduction, as exemplified by producing more, longer, and narrower stolons that continued to emerge at higher nodes, particularly at earlier stages of development. They also exhibited shorter internodes on primary stems, resulting in an overall shorter rosette. Lastly, they flowered earlier and, on average, at earlier nodes than the coastal perennial *M. guttatus* line (although see Coughlan et al. 2021). These results suggest that repeated high altitude adaptation involves several quantitative traits that respond similarly across divergence times.

Most strikingly, all high altitude taxa consistently produced rhizomes while coastal perennial *M. guttatus* lacked rhizomes entirely (Fig. 1a-1b; Table S1). Within the high altitude taxa, rhizomes varied in length, color, and node characteristics (Fig. 1d; Table S1). For example, the rhizomes of *M. decorus* (IMPO) exhibited increased anthocyanin pigmentation and tended to be longer with extended internodes and smoother nodes, while the rhizomes of *M. corallinus* (EAM) had reduced anthocyanin pigmentation, shorter internodes, and more prominent swelling at the node. The rhizomes of *M. tilingii* (LVR) and high altitude altitude *M. guttatus* (CWF) were notably shorter and completely lack pigmentation. These results suggest that although rhizomes are strongly associated with high altitude habitats in this species complex, there is much flexibility in the morphology of these below ground structures. Additionally, this phenotypic diversity may either suggest independent modifier loci involved in pigmentation or length, and/or an entirely separate genetic basis of rhizome production; a hypothesis we explore below.

This consistent pattern of phenotypic differentiation is indicative of repeated phenotypic evolution in response to similar selective pressures in high altitude environments. To test this hypothesis, we constructed four F2 mapping populations between a common, low altitude coastal *M. guttatus* line (OPB) and a representative line from each high altitude taxa (Fig. 1c). For each mapping population we grew between 430-582 F2 individuals, resulting in a total of 2,309 plants. For each trait/cross combination, we calculated Fraser’s *v*-test, which compares the variance among parental lines to the variance within an F2 population to detect divergent selection (Fraser 2020). Fraser’s *v*-test revealed a handful of traits with evidence of divergent selection, particularly for rhizome and stolon traits (Table S2). By contrast, floral, phenological, and size-related traits were less likely to exhibit consistent evidence of directional selection. This suggests that both above- and below ground vegetative traits may be frequent targets of selection in high altitude environments.

### Phenotypes form repeatable modules across divergence time

Phenotypic correlations in advanced generation hybrids can reveal genetically correlated traits (i.e. traits that share a genetic basis via pleiotropic loci or are controlled by loci that are tightly linked). Such correlated traits are sometimes conceptualized as modules; trait groups that evolve together and can facilitate or constrain natural selection, depending on the direction of the correlation and the direction of natural selection (Diggle 2014; Melo et al. 2016). We next calculated pairwise correlation coefficients for each trait within each population. We find patterns of trait correlations were highly similar among crosses (Fig 2a; Mantel tests between correlation matrices for each pair of cross: *r^2^* ranged from 0.62-0.78, all *p-*values <0.0001), suggesting that such correlations are, on average, conserved across the *M. guttatus* species complex. Nonetheless, similarity among these trait correlation matrices tended to decline with divergence, although this trend was not statistically significant (Fig 2b; *r^2^*=0.61, *p*=0.067). These results suggest that despite significant conservation, such correlations may evolve.

**Fig. 2:**
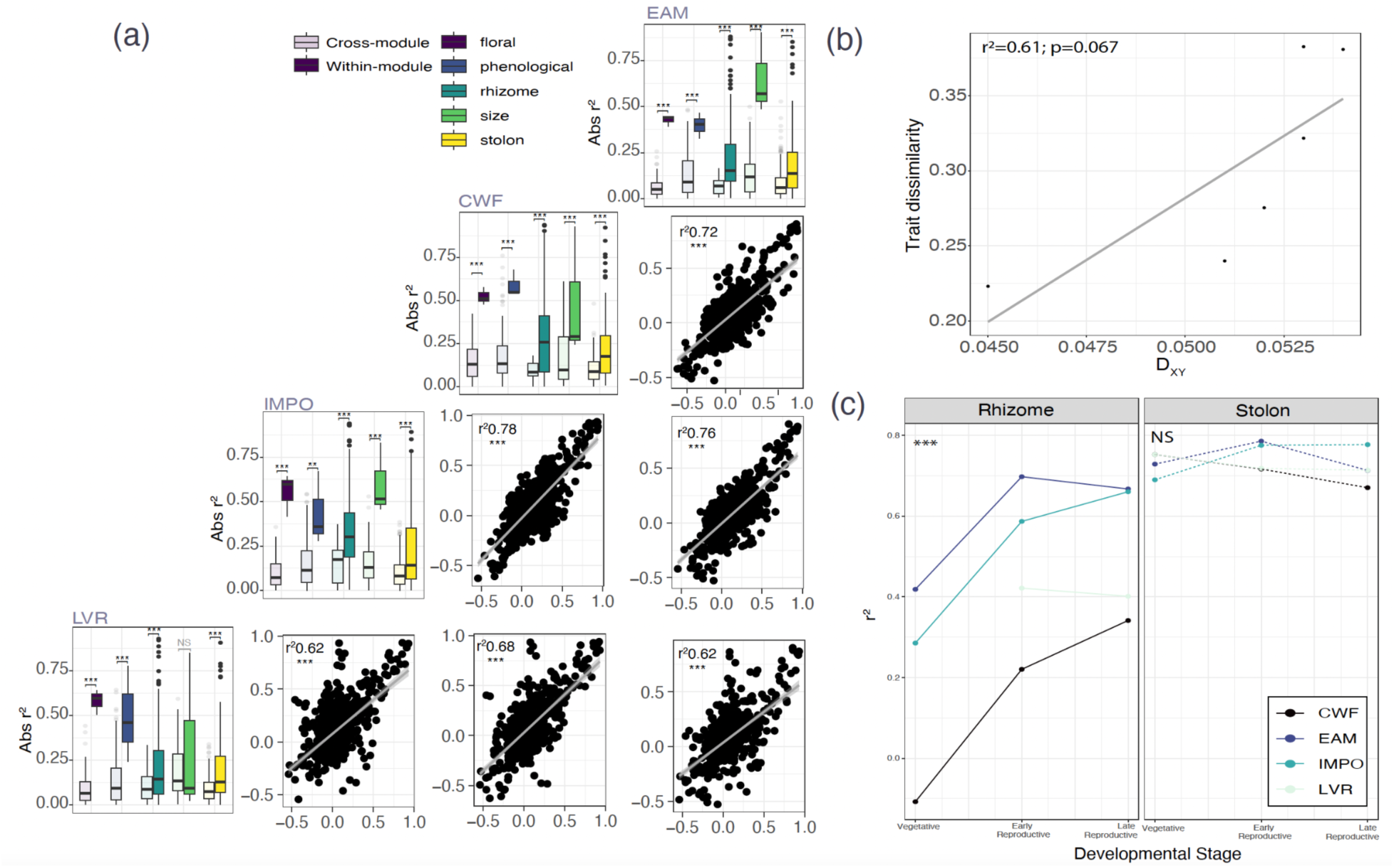
Traits form repeated modules across replicate crosses. (a) Absolute correlation within and between trait modules for each cross (diagonal) and correlations among mapping populations (lower triangle). (b) Dissimilarity among trait-correlation matrices increases with divergence time. (c) Trait correlations of the same trait across development for rhizome-related and stolon-related traits.

In order to assess whether these correlated traits formed evolutionary modules, we then categorised traits as: floral, phenological, size, stolon, or rhizome-related. We asked whether such groupings had higher average correlations than between-category correlation coefficients, and found that these pre-defined trait categories were highly correlated in all cases except for size-related traits in the *M. guttatus* x *M. tilingii* cross (Fig. 2a;; TableS3). Moreover, the average correlation coefficient was higher for these pre-defined trait categories for all crosses than when traits were randomly permuted across categories (again, except for size-related traits in the *M. guttatus* x *M. tilingii* cross; Table S3). Nonetheless, despite this modularity, there were some strong correlations among traits between modules (Fig. S2). In particular, phenological traits were strongly correlated with both floral size traits as well as the number of stolons in all crosses (Fig. S2), recapitulating known life history tradeoffs that have been previously identified within and between annuals and perennials in *Mimulus* (Friedman et al. 2015; Coughlan et al. 2021; Coughlan et al. 2021) as well as other species (Sun and Frelich 2011; Lundgren and Des Marais 2020).

To assess the stability of trait modules across development, we asked if trait correlations changed for traits that were measured across all three sampling periods. These include the number of rhizomes and stolons produced and the highest node of emergence for both rhizomes and stolons. We found that stolon- and rhizome- related traits exhibited different trends across development (module*development interaction: *F*=4.56, *p*=0.028). For all crosses, stolon-related traits maintained consistent and high correlations across all developmental stages (Fig. 2c; Table S4; all developmental contrasts: *p*>0.05), while rhizome-related traits tended to become more integrated across development (Fig. 2c; Table S5; vegetative:late reproductive stage contrast: *p*=0.0052). These findings suggest that modularity is a function of development for some traits, but also that the role of development in defining modules is consistent across independent crosses.

### The genetic basis of high altitude adaptation is shared across integrated trait modules, but largely unique to each evolutionary incident

To assess whether repeated phenotypic evolution is caused by shared genetic bases, we performed QTL mapping in all four mapping populations. In one mapping population (*M. corallinus* x coastal *M. guttatus*) we found extreme, genome-wide segregation distortion likely due to strong crossing barriers between these species (Fig.S3; Frayer unpublished data) which limited our power to detect QTLs. We therefore exclude this cross from most of the following formal analyses of overlap (although we include them in the individual QTL analyses; see below). In total, we identified 262 QTLs across the remaining three mapping populations (115 in the *M. tilingii* x coastal *M. guttatus* cross; 77 in the high elevation *M. guttatus* x coastal *M. guttatus* cross; and 70 in the *M. decorus* x coastal *M. guttatus* cross). These QTLs were distributed across all fourteen chromosomes, with the highest density observed on chromosomes 5 and 6 (Fig. 3a and 3b; Table S6). Across all traits and all crosses, the additive effect sizes of these QTLs were small to moderate (mean effect over all traits and all crosses: 8.1% *r^2^*, range: 0.08-41.2%), with only 6 QTLs having an effect size of >20% of the F2 trait variance, in concordance with a largely polygenic basis for most traits (Fig. S4). In line with this, we find a strong relationship between the number of QTLs identified and the average effect size for each trait (Fig. S5; CWF: *p*=0.027, IMPO: *p*=0.009, LVR: *p*=0.005). Despite a largely polygenic basis for most traits, we find that modules significantly differ in genetic architecture, with rhizome-related traits consistently having a more simple genetic basis than traits belonging to any other module (*F*=9.6, *df*=4, *p*<0.0001; Fig. S4). Additionally, although rare, we do identify 30 epistatic QTL interactions, one of which corresponds to a previously identified epistatic interaction (i.e. LG9/11 epistatic effect for the proportion of stems belowground in the coastal *M. guttatus* x *M. decorus* cross; Coughlan et al. 2021; Table S7). In total, these results suggest that many of these life history traits exhibit different genetic architectures, with some being much more polygenic than others.

**Fig. 3:**
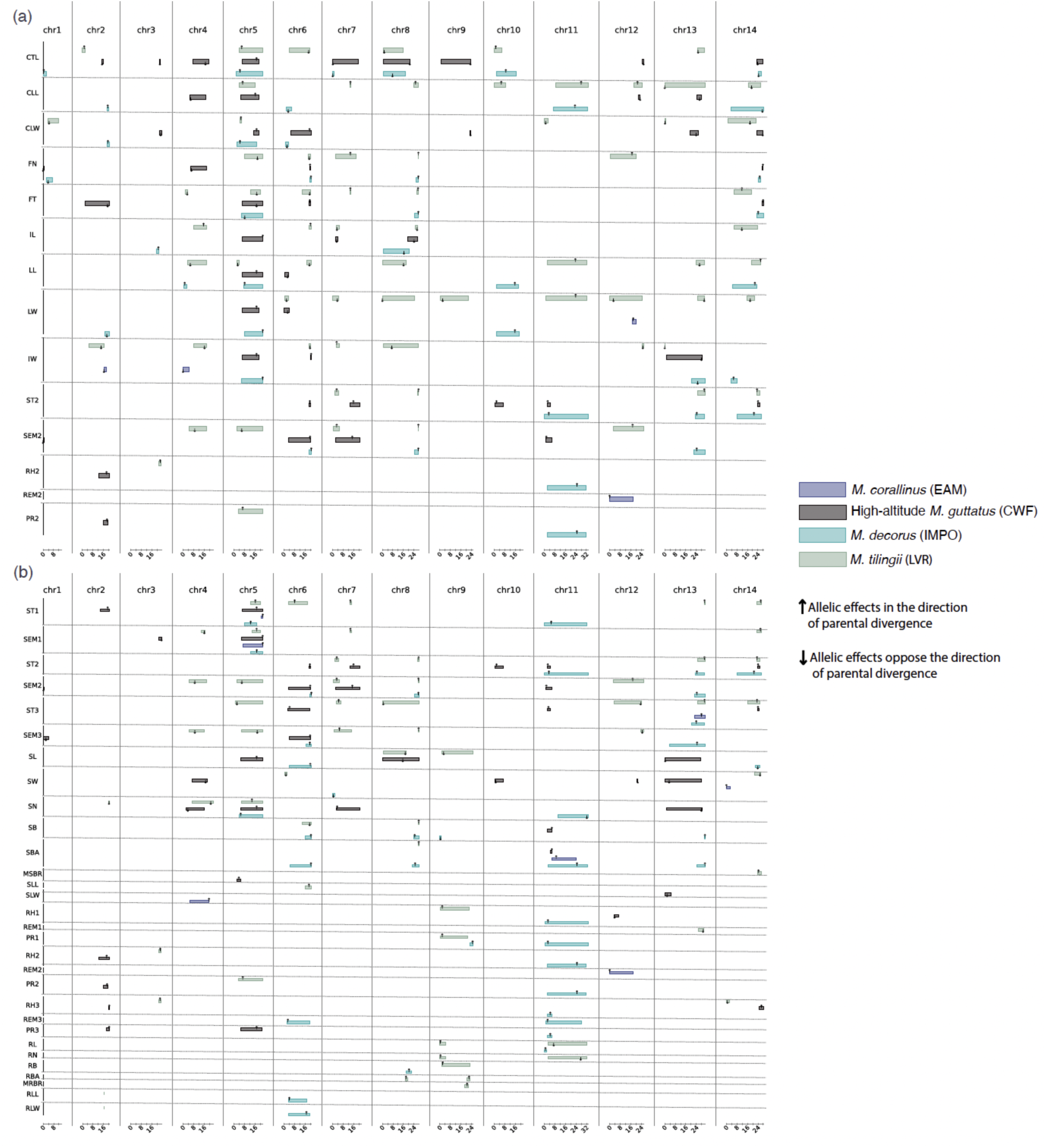
QTLs for life-history trait divergence across the four mapping populations. (a) QTLs associated with various life-history traits measured during the ’early reproductive’ developmental stages. F, floral traits; P, phenological traits; S, size traits; T, stolon traits; R, rhizome traits. (b) QTLs associated with stolon traits and rhizome traits at three developmental stages. T1: stolon traits at the ’vegetative’ stage; T2: stolon traits at the ’early reproductive’ stage; T3: stolon traits at the ’late reproductive’ stage; R1: rhizome traits at the ’vegetative’ stage; R2: rhizome traits at the ’early reproductive’ stage; R3: rhizome traits at the ’late reproductive’ stage. represent the 1.5 LOD-drop confidence intervals arrows denote the QTL peak positions upward arrows indicate a QTL effect in the direction expected from parental divergence, while downward arrows indicate the opposite. The x-axis indicates the genomic positions on each of the fourteen chromosomes (in 8 Mb units).

We next assessed the relationship between trait modularity and QTL overlap to determine whether a shared genetic basis as inferred by F2 trait correlations could be attributable to shared QTLs. We find that the phenotypic correlation among F2s was strongly correlated with the extent of QTL overlap between traits (Fig. 4b; *r^2^*=0.284, *p*<0.0001). This trend was also recapitulated at the level of modules: traits that fell within the same module were, on average, more likely to have overlapping QTLs, although this trend differed by both module and cross (Fig. 4c; cross*module type*within vs between module status: *F*=3.14, *df*=8, *p*=0.0015; Table S8). In all three crosses, rhizome- and size-related traits were more likely to share QTLs with other within-module traits, and in ⅔ of crosses, stolon-related traits were also more likely to share QTLs with each other than with traits in other modules. By contrast, floral-related traits were not more likely to have shared QTLs in any cross (Fig. 4c). Lastly, we find that QTL overlap increased across developmental stages for both rhizome- and stolon-related traits (Fig. 4a; Table S9 and S10), suggesting that investment in either above- or below-ground biomass post-reproduction shares a genetic basis regardless of plant age, while investment in these traits pre-reproduction may be more genetically independent. In total, these results support the notion that phenotypically integrated trait modules are more likely to partially share a genetic basis, and again that this modularity can vary across development.

**Fig. 4:**
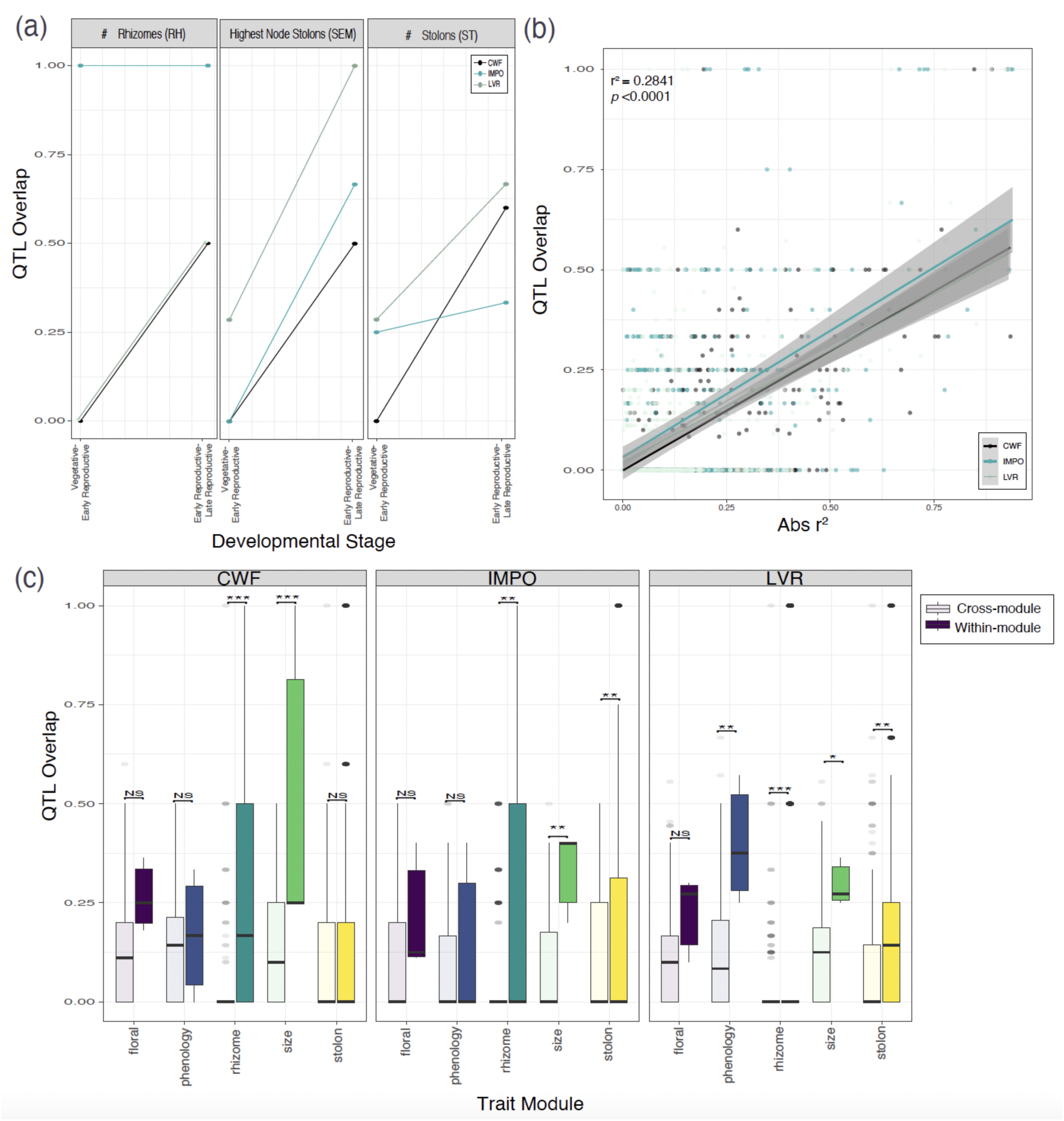
Correlated traits are more likely to share QTL. (a) The extent of QTL sharing between developmental stages for rhizome- and stolon-related traits. (b) The relationship between absolute trait correlation and the extent of QTL overlap for each pair of traits and each cross. (c) The extent to which QTL are shared for traits that are within versus between trait modules for each cross.

### Repeatability between populations is low, but influenced aspects of the genetic architecture and divergence time

We next compared the degree to which the genetic basis of life-history divergence was shared among the four replicated cross populations. For all traits, we find that no pair of crosses shares more QTLs than expected by chance (Fig. 3; Table S11), suggesting that overall the genetic basis of life-history divergence was largely not repeatable. Most striking in this trend was the genetic basis of rhizome-related traits, in which no QTLs were shared across any pair of populations.

Although there is no more QTL sharing than expected by chance, there are many individual QTLs that are shared among crosses (except for rhizome-related traits, as noted above). We therefore next sought to understand if certain factors increased the likelihood that a given QTL would be shared among crosses. Specifically, we asked whether the probability that an individual QTL was identified in more than one cross was related to aspects of the genetic architecture (*r^2^*, *a, d,* epistasis, pleiotropy) or the history of directional selection on that trait (Fraser’s *v-*statistic). We find that the probability that a QTL was found in more than one cross was strongly related to its effect size, wherein larger effect QTLs were more likely to be shared (Fig. 5a; *r^2^*: *χ^2^*= 5.9, *df*=1, *p*=0.015). Additionally, QTLs that were implicated in more than one trait were more likely to be shared across crosses, in line with a positive relationship between pleiotropy and repeatability (Fig. 5b; Fisher’s exact test: odds ratio=3.5; *p*=0.049). We do not find a significant association between the probability that a QTL was shared across crosses and the average additive effect (Fig.5e; here we standardized the additive effect, *a*, by the parental difference to compare across traits and crosses: *a*/parental difference: *χ^2^*=1.9, *p*=0.15), dominance deviation (Fig.5f; *d/a: χ^2^*=0.04, *p*=0.83), or whether or not a QTL had a significant epistatic effect (Fig.5c; Fisher’s exact test: odds ratio=1.6; *p*=0.11). Additionally, we do not find a significant association between the probability a QTL was identified in more than one cross with our measure of divergent selection (Fig. 5d; *v-* value: *p*=0.9). In total, this suggests that although the genetic basis of life history divergence is largely unshared among replicate incidences of high altitude adaptation in the *M. guttatus* species complex, certain aspects of genetic architecture may influence whether individual QTLs are shared.

**Fig. 5:**
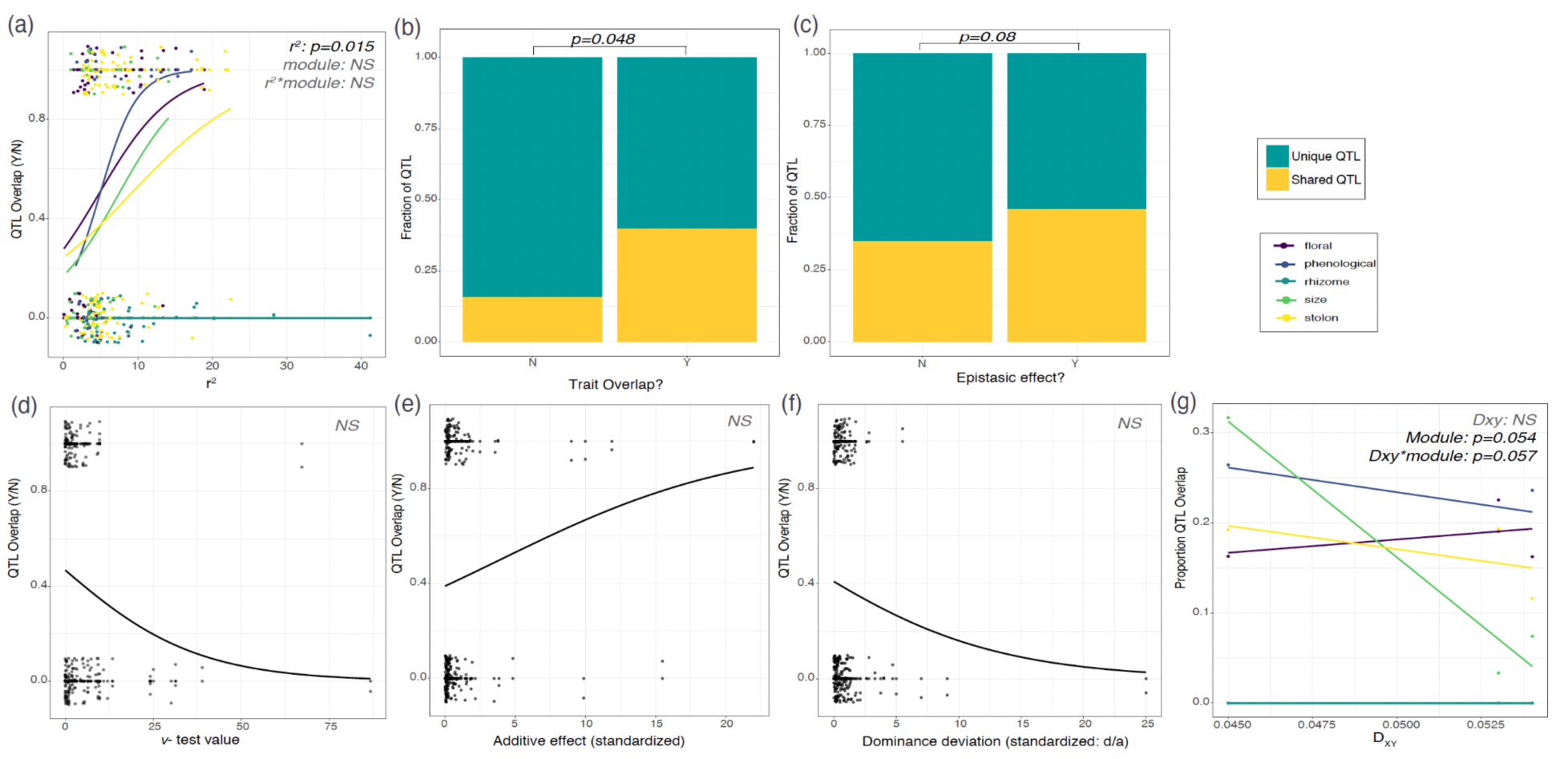
Factors that influence the extent of QTL sharing. (a) the QTL effect size (*r^2^*). (b) Whether a QTL was identified in more than one trait (a proxy for pleiotropy). (c) Whether a QTL had an epistatic effect (d) Fraser’s *V*-statistic (a proxy for a history of directional selection). (e) The additive effect of a QTL (standardized by the difference between parental lines. (f) The dominance deviation of a QTL (calculated as d/a). (g) Divergence time (D_XY_).

Lastly, we also asked whether the proportion of QTLs that overlap between a pair of crosses was related to divergence between the high altitude adapted lines. We find a nearly significant trend that the extent of QTL overlap among crosses declined with DXY, but that this trend differed among trait modules (Fig. 5g; DXY*module: *F*=4.8, *df*=4, *p*=0.057). Although this finding is in broad agreement with theoretical predictions (Conte et al. 2012; Bohutínská et al. 2021; Bohutínská and Peichel 2024), it is important to note that we are much less likely to detect overlap for small effect QTLs, given our more limited power to detect individual small effect QTLs multiple times and the observation that there are many more small effect QTLs than large effect ones.

### Strong candidate genes for FT, FN, SB, SBA, and RH3, across the three mapping populations

To identify plausible candidate genes underlying life-history trait divergence, we extracted all annotated genes from QTL intervals ≤1 Mb in size (32 QTLs in total; Table S12). Candidate genes were further prioritized within each QTL interval based on their known or predicted functions from genome annotations. This approach yielded strong candidate genes associated with several key traits, including flowering time, node of first flower, the number of stolon branches, average branches per stolon node, and the number of rhizomes at the latest stage of development across the three mapping populations.

In the LVR population, FT7 harbors *Migut.07G085800*, which encodes a spliceosome-associated protein potentially involved in temperature-sensitive alternative splicing and circadian regulation, suggesting a post-transcriptional mechanism influencing flowering time (Schlaen et al. 2015). In the CWF population, FT6 includes *Migut.06G177100*, a homolog of *FLOWERING LOCUS T* (*OsFTL2*), a central integrator of photoperiodic flowering signals (Kojima et al. 2002; Hayama et al. 2003; Abe et al. 2005; Komiya et al. 2008). In FT14 of the same population, *Migut.14G297800* encodes histone H3, a core chromatin component implicated in the epigenetic regulation of flowering through histone methylation and transcriptional control (Tamada et al. 2009; Zhang et al. 2021; Li et al. 2025). Also, within FT14, *Migut.14G300800* encodes a COP1-interacting RING finger protein (CIP8 homolog), is known to interact with COP1 (Hardtke et al. 2002); COP1 is a key repressor of photomorphogenesis and flowering under non-inductive photoperiods (Sarid-Krebs et al. 2015; Xu et al. 2016; Lee et al. 2017). For FN14 in the CWF population, *Migut.14G299900* encodes a GASA/GAST/Snakin family peptide, which is responsive to gibberellin and known to modulate flowering time and stem elongation (Roxrud et al. 2007; Bouteraa et al. 2023; Chen et al. 2024). Moreover, *Migut.01G000800* within FN1 encodes the CHD3-type chromatin remodeling factor PICKLE, which influences *FT* expression and meristem identity (Fu et al. 2016; Jing et al. 2019; Yoon et al. 2021). In the LVR population, SBA8 harbors *Migut.08G238300*, a leucine-rich repeat (LRR) receptor-like kinase involved in meristem maintenance and tiller formation (DeYoung et al. 2006; Kang et al. 2017). Within SB9 of the IMPO population, *Migut.09G000800* encodes a TCP transcription factor, a well-established integrator of branching signals that suppresses axillary bud outgrowth (Takeda et al. 2003; Aguilar-Martínez et al. 2007; Gastaldi et al. 2024). For rhizome number, we highlight *Migut.02G173500* within RH3_2 of the CWF population, which encodes a DEAD/DEAH box RNA helicase previously implicated in regulating rhizome number in *Chrysanthemum morifolium* under drought stress (Zhang et al. 2022) .

### Chromosomal inversions, modularity, and repeatability

Chromosomal inversions are one mechanism that can underlie genetic correlations among traits and are commonly associated with ecological divergence, particularly in taxa that experience significant gene flow, such as those in the *M. guttatus* species complex (Lowry and Willis 2010; Wellenreuther and Bernatchez 2018; Huang et al. 2020; Funk et al. 2021; Koch et al. 2021;

Akopyan et al. 2022). Three known chromosomal inversions segregate within these crosses: one on chromosome 5 (coastal *M.guttatus* x *M. tilingii* cross and the coastal *M.guttatus* x *M. decorus*; Garner et al. 2016), one on chromosome 8 (coastal *M.guttatus* x *M. tilingii* cross; Lowry and Willis 2010; Garner et al. 2016; Coughlan and Willis 2019), and one on chromosome 13 (coastal *M.guttatus* x *M. tilingii* cross and the coastal *M.guttatus* x *M. decorus*; (Garner et al. 2016; Frayer unpublished data). Although we find a high density of QTLs encoding multiple traits in these inversion-regions (particularly for chromosome 5), we do not find an excess of QTLs in crosses where the inversion segregates versus those in which both parental lines were collinear (chromosome 5: Fisher’s Exact test: odds ratio=2.02, *p*=1; chromosome 8: Fisher’s Exact test: odds ratio=1.05, *p*=1, chromosome 13: odds ratio=0.81, *p*=1). This suggests that although some of these inversions have been shown to play a central role in local fitness in other contexts (i.e. annual and perennial ecotypes of *M. guttatus*; Lowry and Willis 2010), the inversions themselves are likely not involved in high altitude adaptation (at least for the phenotypes we include herein). Furthermore, the abundance of segregating QTLs in inversion-regions (whether or not they actually contain an inversion) may support the idea that inversions capture pre-existing genetic diversity, in line with inversions playing an adaptive role in ecological divergence via recombination suppression. However, we note that a knowledge of the ancestral states of these loci would be required to further test this idea.

### High altitude adaptation and trait novelty: rhizomes have independently evolved at least four times in the *M. guttatus* complex

Among all of the 40 traits measured across all four mapping populations, rhizome-related traits were unique in a number of qualities. They showed the strongest signals of divergent selection and exhibited the least polygenic genetic architecture, typically involving one or two QTLs, although in some cases with epistatic effects. Previously, we described an intriguing pattern of development wherein F1 and F2 hybrids tended to invest more in rhizome production across development, while parental lines did not (Coughlan et al. 2021). We determined that this pattern was caused by two epistatically interacting QTLs at the onset of reproduction, only one of which was significantly associated with rhizome production later in development (Coughlan et al. 2021). We hypothesized that one such QTL controlled the ability to produce rhizomes and the second modulated to the timing of rhizome production (Coughlan et al. 2021). Intriguingly, all but the *M. corallinus* x coastal *M. guttatus* cross showed this pattern of rhizome investment, wherein F1 and F2 hybrids invest more in rhizomes as development proceeds, while parental lines remain relatively consistent in rhizome investment (Fig. 6). In the current study, we again recover two QTLs in the coastal perennial *M. guttatus* x *M. decorus* cross that act epistatically earlier in development, but only one of which remains associated with rhizome investment later in development (although we note that the QTLs exhibited a significant epistatic effect at an earlier stage of development). We also uncover direct evidence of epistasis in the coastal perennial *M. guttatus* x high altitude *M. guttatus* cross (Fig. 6; *F*=6.47, *p*=0), and non-significant patterns of non-additivity in the coastal perennial *M. guttatus* x *M. tilingii* cross at later developmental stages (Fig. 6). Despite this similar developmental trajectory in ¾ of our mapping populations, we find that the QTLs implicated in rhizome production are entirely different in all crosses. Even in the coastal perennial *M. guttatus x M. corallinus* cross, in which severe segregation distortion prevents us from confidently comparing QTLs across mapping populations, the distribution of traits in the F2 strongly suggests an independent genetic basis. In total this suggests that there are many genetic mechanisms by which to produce rhizomes despite some potentially shared aspects of genetic architecture. This may also explain the phenotypic diversity that we see within rhizome production in this group (i.e. Fig. 1).

**Fig. 6:**
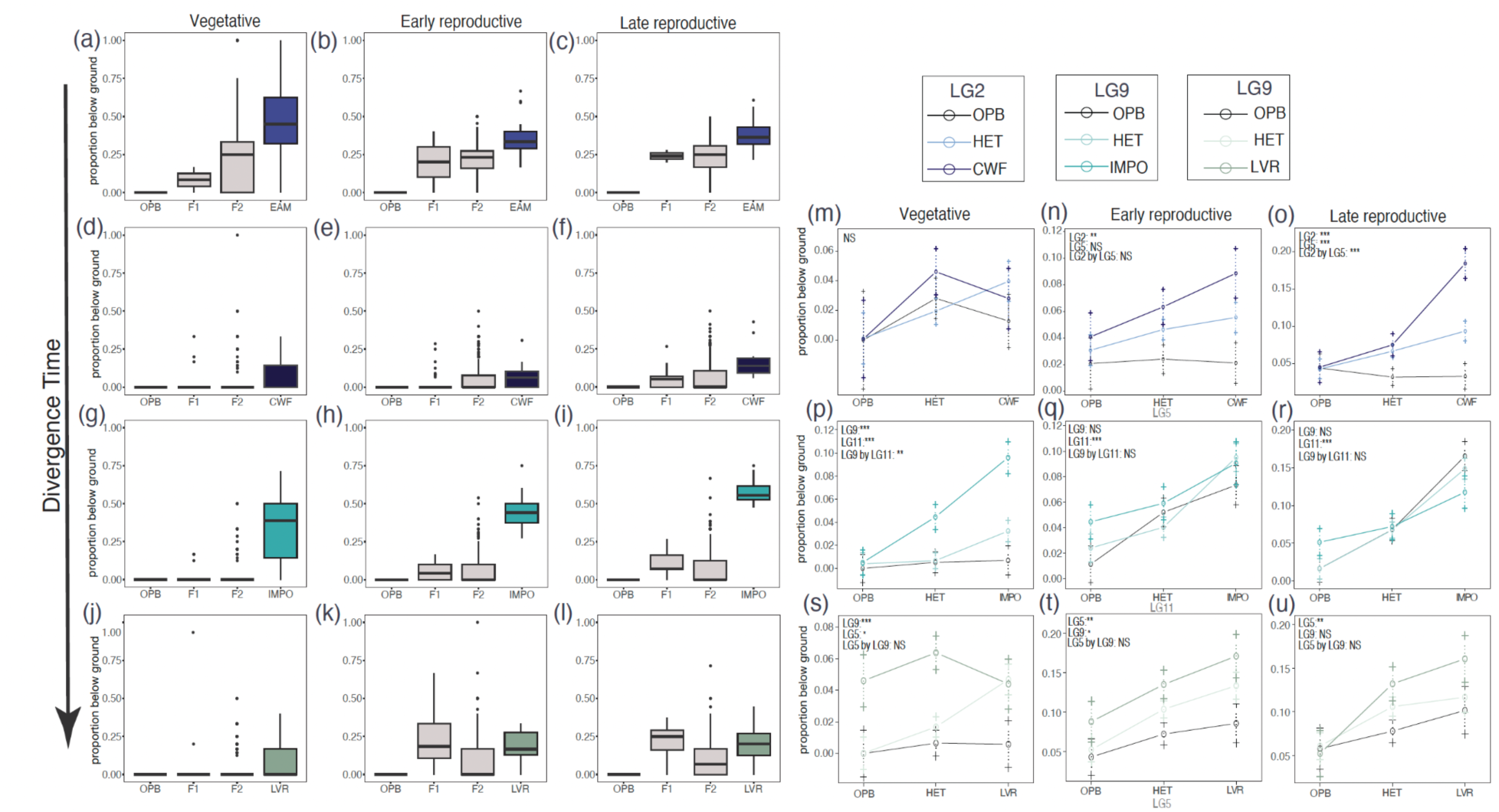
The developmental basis of rhizome production is similar across divergence time, but utilizes different genes. (a-l) Developmental trajectory of investment in rhizomes for all 4 mapping populations at 3 developmental times (EAM: a-c; CWF: d-f; IMPO g-i; LVR: j-l). In all crosses, the parental lines exhibit relatively stable phenotypes through time, while in ¾ of crosses hybrids tend to increase investment across development. Despite this shared developmental trajectory, each incident of rhizome production exhibits a different genetic basis, although potentially with some shared aspects of architecture. (m-u): Effect plots for ¾ of crosses across 3 developmental times. (m-o): coastal perennial *M. guttatus* x high altitude *M. guttatus*. (p-r): coastal perennial *M. guttatus* x *M. decorus*. (s-u): coastal perennial *M. guttatus* x *M. tilingii*.

## Discussion

Here we explored the extent of evolutionary repeatability of adaptation to high altitude habitats in closely related perennial taxa of the *M. guttatus* species complex. We quantified repeatability across scales, including univariate traits, correlated trait modules, and QTLs. Across four incidences of high altitude adaptation, we find that high altitude plants show repeated patterns of trait evolution, resulting in greater investment in vegetative biomass (such as producing more stolons and from higher nodes), shorter internodes, and earlier flowering. These traits exhibited strong genetic correlations, and formed predictable modules that were highly repeatable across divergence times. We find that life-history traits exhibited a diversity of genetic architectures, including many polygenic traits and at least one trait with a relatively simple genetic basis (i.e. rhizome production). Although we find significant QTL overlap within crosses for highly correlated traits, QTLs were not shared among crosses more than expected by chance. Nonetheless, the extent of repeatability at individual QTLs was related to aspects of the underlying genetic architecture, such as effect size and pleiotropy. Our results provide insights into the genetic architecture of life-history divergence and high altitude adaptation, as well as inform our understanding of evolutionary repeatability of complex, quantitative traits.

### High altitude adaptation involves repeated phenotypic evolution

High altitude habitats typically impose myriad challenges to plant life, including a shorter growing season, harsher winters, lower partial pressure of CO2, increased UV reflectance, and differences in biotic communities (including both herbivores and pollinators; Billings 1974). It’s therefore unsurprising that many traits have repeatedly evolved across four replicated incidences of high altitude adaptation in the *M. guttatus* species complex. We find that higher altitude taxa invested significantly more in vegetative biomass and asexual reproduction via stolons, as well as rhizome production, in line with previous work in many plant taxa (Billings 1974; Hautier et al. 2009; Kim and Donohue 2011; Šťastná et al. 2012; Coughlan et al. 2021; Doležal et al. 2024). Greater investment in asexual reproduction may aid in high altitude or high latitude habitats by providing reproductive reassurance in areas of patchy pollinators, low plant density, or harsh weather events that damage floral tissue (such as early growing season frosts; Inouye 2008; Urquhart-Cronish et al. 2024). They may also allow individuals to “explore” patchy habitats while mitigating the cost of such exploration (i.e. clonal integration; Chen et al. 2006; Liu et al. 2016). For example, greater investment in clonal reproduction has been associated with increased resilience to herbivore damage (Liu et al. 2007), and increased drought resistance (Zhou et al. 2014), and therefore may also serve as a form of bet hedging in heterogeneous habitats.

We also found that all high altitude taxa flowered earlier (and ¾ of taxa flowered at earlier nodes), which is common (although not ubiquitous) in other high altitude adapted plants (Anderson and Gezon 2015; Wang et al. 2021; Barnes et al. 2022, though see Halbritter et al. 2018; Kooyers et al. 2019; Gamba et al. 2024), although differs from previous findings in *Mimulus* (Kooyers et al. 2015; Coughlan et al. 2021). Flowering time is strongly correlated with fitness in many wildflower systems (Hall and Willis 2006; Anderson, LEE, et al. 2011; Fournier-Level et al. 2022), and has been implicated as a phenotype under selection specifically at high altitude in many plant taxa (Méndez-Vigo et al. 2011; Anderson and Gezon 2015; Geng et al. 2021; Wang et al. 2021; Barnes et al. 2022; Gamba et al. 2024). Earlier flowering may allow plants to maximize sexual reproductive effort during short growing seasons. On the other hand, flowering too early in a season may expose individuals to spring freezing events causing floral damage or create mismatches between sexual reproductive effort and peak pollinator densities (Pardee et al. 2019). The most striking phenotypic difference between high and low altitude taxa was the presence of a novel trait: rhizomes. In the *M. guttatus* species complex, high altitude perennial species uniformly produce rhizomes, while in the broad-ranging *M. guttatus,* this trait is more variable, but is associated with plants from higher altitudes and latitudes (Coughlan unpub. data). Rhizomes exhibit tremendous phenotypic diversity across the four high altitude taxa sampled here, which aligns with our finding that the genetic basis of rhizome production was entirely independent in each cross. Rhizomes are storage organs, and as such are hypothesized to confer adaptation to harsh environments by allowing plants to survive in more stable and warmer belowground environments when aboveground air temperatures are extremely cold, or under freeze-induced drought, but also by allowing plants to regrow more rapidly in short growing seasons (Mooney and Billings 1960; Hiltbrunner et al. 2021). We find modest evidence of divergent selection on rhizome traits in high altitude taxa using Fraser’s *v*-test, particularly in the *M. decorus*, *M. corallinus*, and high altitude *M. guttatus* crosses. In the case of *M. tilingii*, a lack of evidence of divergent selection may stem from the fact that plants produced fewer and smaller rhizomes in our relatively ‘low altitude’ common garden environment than they typically do in their native habitat (Coughlan pers. obs). Such minimal parental differences (and variance among replicates of our *M. tilingii* parental line) may have limited our power to detect signatures of divergent natural selection. Future manipulative experiments and fieldwork will be required to directly test the adaptive value of each of these traits, including rhizomes, as well as assess potential agents of natural selection.

### Correlated traits, trait modularity, and repeated adaptation to high altitude

High-altitude elevation adaptation often involves the coordinated changes of multiple traits (Halbritter et al. 2018; Duruflé et al. 2019). Patterns of correlation were highly conserved across our populations (Fig. 2a), indicating a repeatable framework of phenotypic integration. This conserved structure likely arises from shared developmental programs or functional coupling among traits (Pigliucci 2003; Klingenberg 2008; Armbruster et al. 2014), providing a foundation for evolutionary repeatability among taxa. Notably, the slight decline in matrix similarity with increasing divergence time (Fig. 2b) suggests that the overall structure of trait correlations is itself evolvable, potentially providing a mechanistic basis for taxa-specific differences in phenotypic evolutionary trajectories.

At the modular level, trait modules exhibited high repeatability across taxa (Fig.2a), supporting the hypothesis that trait modules facilitate adaptive evolution (Wagner and Altenberg 1996; Melo et al. 2016). However, these modules were not entirely independent (Fig. S2); Some degree of coupling among certain modules suggests that such structural interdependence may limit their potential to evolve as fully independent units. Instead, it may promote coordinated phenotypic responses under complex environmental conditions. We further found that module status shifts across developmental stages (Fig. 2c), indicating that modularity is not a static feature, but rather a dynamic attribute embedded within the organism’s ontogeny. Although these phenotypic modules partly reflect a shared genetic basis (Fig. 4c), modularity is fundamentally a product of the long-term interplay among genetic effects, developmental regulation, and functional integration (Klingenberg 2008; Assis et al. 2016), rather than a fixed genetic unit. This balance of structural constraint and developmental flexibility may underlie the ability of modules to respond in parallel to selection across taxa, serving as a key mediator of phenotypic evolutionary repeatability.

At the level of individual traits, correlations among traits are also subject to dynamic regulation by both genetic architecture and developmental processes. Across all four crosses, flowering time and the number of stolons produced early in the vegetative stage consistently exhibited strong negative correlations (Fig. S2; Fig. S6), recapitulating the early-flowering, increased stolon number observed in parental lines (Table S1). In contrast, during all reproductive stages flowering time was strongly positively correlated with the number of stolons produced (Fig. S6), which recapitulates well established correlations and apparent life-history tradeoffs (Friedman et al. 2015; Coughlan et al. 2021). This divergence in correlation likely reflects differences in the genetic and developmental bases of these traits: stolon production in the vegetative stage is controlled by a small number of large-effect QTLs (Table S6) and is strongly associated with overall plant size (i.e. internode width; Fig. S2), suggesting that it may represent an early developmental switch trait linked to growth thresholds. In comparison, stolon production during the reproductive phase (i.e. ST2 and ST3) more likely represent dynamic outcomes of resource allocation during development, shaped by multiple loci and regulatory pathways, as has been previously hypothesized (Friedman et al. 2015). As a result, their expression in hybrid progeny is more susceptible to recombinational disruption, giving rise to phenotypic combinations (e.g., late flowering and increased stolon number) that better align with classic life-history trade-off expectations (Friedman et al. 2015; Rubin et al. 2019).

### The complex genetic architecture underlying life-history divergence

Life-history divergence in perennials of the *M. guttatus* species complex was largely polygenic. Specifically, for floral, phenological, stolon, and size traits, QTLs effect size ranged from 0.08-22.5% of the variance in F2s (which corresponds to 7-100% of the heritable variation). The finding that such quantitative traits are largely controlled by many minor-effect QTLs is consistent with previous work, both in *Mimulus* and other flowering plants (Salisbury et al. 1987; Fishman et al. 2002; Galliot et al. 2006; Hall et al. 2006; Friedman et al. 2015; Ahsan et al. 2019; Feng et al. 2019; Fournier-Level et al. 2022; Minadakis et al. 2024; Chen et al. 2025). More broadly, work leveraging population genomic data comparing low and high altitude populations also suggests that adaptation to high altitudes involves many genes (Kubota et al. 2015; Holliday et al. 2016; Bohutínská et al. 2021; Geng et al. 2021; Wang et al. 2021), although in many cases the specific traits involved (and the polygenicity of each trait) are unknown.

Rhizome-related traits were significantly more simple, typically involving one or two QTLs, which aligns with previous work in *Mimulus* (Coughlan et al. 2021; Nelson et al. 2021). Our findings are in contrast to work in several crop species (i.e. rice (Li et al. 2022), sorghum (Kong et al. 2015), wildrye (Larson et al. 2014), and lotus (Huang et al. 2021)), where rhizome traits involving their expression and abundance (e.g., branching and length) are generally highly polygenic. This inconsistency may arise from many factors, including human-mediated selection on quantity versus presence/absence traits under natural selection in the wild. The difference in genetic architecture may also reflect taxonomic divergence, as most of these crops are grasses (i.e. family Poaceae, with the exception of lotus, which is in the family Nelumbonaceae), while *Mimulus guttatus* species complex taxa are small herbs in the family Phrymaceae. Despite their preponderance in high elevation habitats, very few studies have investigated the genetic basis of rhizomes in non-crop species, and much work is needed to assess the commonality of simple versus polygenic architecture of this ecologically important trait.

### QTLs underlying life-history divergence are not shared among populations

Much recent work in repeated evolution has begun to explore the dynamics of repeated evolution of complex traits (Bohutínská and Peichel 2024; Nosil et al. 2024; Gompert et al. 2025; Hoitinga and Birkeland 2025; Huang et al. 2025). In contrast to earlier work on fairly simple traits, work on complex traits reveals mixed evidence for genetic repeatability (Kaeuffer et al. 2012; Konečná et al. 2021; Magalhaes et al. 2021; Konečná et al. 2022; James et al. 2023; Moran et al. 2023; Gutiérrez-Guerrero et al. 2024; St. John et al. 2024; Huang et al. 2025). Only a few studies have begun to disentangle the factors that promote or hamper genetic repeatability of complex traits (Rennison and Peichel 2022; Yeaman 2022; Battlay et al. 2024; Bohutínská and Peichel 2024; Whiting et al. 2024). Here we contribute to this growing body of work by leveraging repeated evolution of several complex traits thought to be involved in high altitude adaptation in the *M. guttatus* species complex. For all traits, we find no excess of QTL sharing among crosses, suggesting that repeatability in this system is, overall, very low. The ability to produce rhizomes in particular exhibited no shared QTLs, suggesting that rhizomes can evolve readily through multiple genetic avenues. Despite no shared QTLs, aspects of rhizome development and genetic architecture are shared in ¾ of our crosses. In these cases, F2 hybrids between coastal perennial *M. guttatus* and each of *M. decorus, M. tilingii,* and high altitude *M. guttatus* all increased their investment in rhizomes across development, despite a fairly consistent level of investment in the parental lines. In each cross we identify one or two QTLs, depending on the developmental stage in question, and in ⅔ of crosses those QTLs have epistatic effects in one developmental stage, but not others. Previously, we hypothesized that these two interacting loci in *M. decorus* may independently control the ability to produce rhizomes and the timing of rhizome production. This model may also be accurate for high altitude *M. guttatus*. Although in the coastal *M. guttatus* x high altitude *M. guttatus* cross, the epistatic effects did not manifest until later in development, this may reflect broad differences in the number of rhizomes produced between these high altitude taxa (*M. decorus* makes many more rhizomes and at earlier stages than does high altitude *M. guttatus*). Although not significant, QTLs in the coastal *M. guttatus* x *M. tilingii* cross also appear non-additive at the latest developmental stage, and similar to high altitude *M. guttatus*, *M. tilingii* makes many fewer rhizomes than does *M. decorus*. Thus, although further fine mapping will be required to fully dissect the developmental genetic basis of rhizome production, a model in which rhizome development is predictably controlled by independent loci involved in rhizome production and timing of production may be a shared aspect of rhizome development.

Although we do not find excess QTL sharing among crosses, we do find individual QTLs that are shared in multiple crosses. QTLs that are implicated in more than one trait (i.e. potentially pleiotropic) as well as large effect QTLs are more likely to be shared. These findings align with recent work in both plants and animals that have found that pleiotropic loci are more likely to be shared in incidences of repeated adaptation (Rennison and Peichel 2022; Battlay et al. 2024; Whiting et al. 2024). Contrary to earlier theoretical work, which suggested a ‘cost of complexity’ (Orr 2000; Welch and Waxman 2003), such large effect, pleiotropic loci underlying adaptation can readily evolve repeatedly, particularly under models of gene flow. Although we strongly note that our reduced power to detect small effect QTL across multiple crosses likely biases our results, the observation that pleiotropic loci were more likely to be shared across crosses aligns with our finding that patterns of trait correlation were highly repeatable across high altitude taxa. Shared QTLs underlying such correlated traits may arise from shared genetic variation (either through introgression or ancient standing genetic variation) or independent mutations for traits that have inherent genetic constraints (i.e. those underlying tradeoffs). Although explicit tests of these non-mutually exclusive hypotheses will require a knowledge of the specific genes underlying life-history divergence, we suspect that genetic variants that are shared by introgression or standing genetic variation may be a major contributor of these potentially pleiotropic QTLs (as well as other QTLs, more broadly). This is because the *M. guttatus* species group is quite young (∼674kya; Sandstedt et al. 2021), exhibits tremendous levels of shared genetic variants, including those underlying life history tradeoffs (Kelly 2022), and many species within the complex have an extensive history of introgression (Brandvain et al. 2014; Ivey et al. 2023; Mantel and Sweigart 2024; Farnitano et al. 2025, though see Sandstedt et al. 2021).

### Functionally diverse candidate genes underlie the genetic architecture of life-history trait divergence

The identification of strong candidate genes across multiple QTLs associated with flowering time and node, branching architecture of stolons, and rhizome production reveals a complex and multilayered genetic basis underlying divergence in life-history traits (Anderson, Willis, et al. 2011; Mojica et al. 2012; Savolainen et al. 2013; Dixit et al. 2017; Troth et al. 2018). These candidate genes span multiple layers of regulatory mechanisms, including chromatin-level regulation (e.g., the CHD3-type chromatin remodeler PICKLE and core histone H3, both involved in aspects of flowering time), post-transcriptional control (e.g., a spliceosome-associated factor implicated in flowering time), putative proteasome-mediated regulation (e.g., a COP1-interacting CIP8 homolog implicated in flowering time), and hormone signaling (e.g., a gibberellin-responsive GASA family peptide implicated in flowering time) (Table S12). The combinatorial effects of these epigenetic, transcriptional, post-transcriptional, and post-translational regulatory pathways likely contribute to the polygenic basis of life-history trait divergence (Fig. 3; Table S6).

Furthermore, comparative analysis across the three mapping populations reveals convergence at the trait level but divergence at the regulatory level. For example, although QTLs associated with flowering time were identified in both the *M. tilingii* and high altitude *M. guttatus* populations, the strongest candidate genes differ in both identity and regulatory mechanism (Table S12), highlighting a population-specific regulatory basis for the same trait. In addition, distinct QTLs within the same population may contain genes that participate in similar regulatory pathways—such as chromatin remodeling—suggesting that different molecular mechanisms can converge to produce similar phenotypic outcomes. For instance, *PICKLE* in and histone H3 (both implicated in aspects of flowering time) represent mechanistically distinct components of chromatin-level regulation, yet both likely influence developmental timing by modulating transcriptional accessibility (Table S12). Such convergence at the pathway level may underscore the presence of trait modules conserved across populations (Fig.2a; Table S3).

Although many of the questions that we set out to answer will require a more thorough investigation of the genes involved in high altitude adaptation, our findings contribute to a growing literature regarding repeated adaptation involving complex traits. We show that natural selection may repeatedly result in shifted quantitative phenotypes, the origin of novel traits (i.e. rhizomes), and the emergence of repeatable trait modules. Nonetheless, there are many genetic paths to similar high altitude adapted phenotypes, but aspects of genetic architecture likely influence what routes are more likely to be taken.

## Materials and Methods

### Plant materials and crossing design

We leveraged repeated incidences of high altitude adaptation to determine the extent of phenotypic and genetic repeatability (Fig. 1). We crossed a common, low altitude, rhizome-lacking, coastal perennial *M. guttatus* (OPB: 42.64, -124.42, 25ft) with each of four high altitude, rhizome-producing taxa include *M. tilingii* (LVR: 37.95, -119.23, 9,027ft), *M. corallinus* (EAM: 38.32, - 119.92, 6712ft), *M. decorus* (IMPO: 44.41, -122.14, 4900ft), and high altitude *M. guttatus* (CWF: 43.25, -122.23, 4173ft). Due to asymmetric seed barriers between *M. decorus* (IMPO) and *M. guttatus* (Coughlan et al. 2020) and between *M. corallinus* (EAM) and *M. guttatus* (Frayer unpublished data), we obtained F1 hybrids in these crosses only when the coastal perennial *M. guttatus* was the maternal donor. For the *M. tilingii* (LVR) and high elevation *M. guttatus* (CWF) crosses, we generated reciprocal F1s. We then generated F2 populations by self fertilizing each F1 hybrid (OPBxIMPO: 469 individuals; OPBxEAM: 582 individuals; LVR: 273 and 270 individuals for OPBxLVR and LVRxOPB, respectively; CWF: 206 and 224 individuals for CWFxOPB and OPBxCWF respectively) We additionally grew 15-26 replicates of each inbred parental lines and 2-41 replicates of F1s for each cross. We note that we grew 15-26 replicates of the common low altitude *M. guttatus* line in each F2 growout so that parental lines and hybrids shared a common environment per cross.

### Plants grow conditions and phenotyping

The individuals from the four crosses were planted sequentially. All plants were cultivated in a greenhouse common garden at Yale University. Initially, seeds were sown into 3" pots (50-100 seeds per pot) filled with wet Promix BX soil and then placed in the dark at 4°C for stratification. After 5 days, the seeds were transferred to the greenhouse for germination. On the day of germination, each germinant was transplanted into an individual 4” pot filled with Promix BX soil. In the greenhouse, the plants were grown under natural light supplemented with sodium lamps to maintain a 16-hour day length and were regularly fertilized (once per day).

For each cross, the individuals (F2s, F1s, and parental lines) were phenotyped at three developmental stages, as adapted from Coughlan et al. 2021: three weeks post-stratification (the ’vegetative’ period), on the day of the first flower (the ’early reproductive’ period), and seven weeks post-stratification (the ’late reproductive’ period). During the ’vegetative’ period, without unpotting the plants, we measured the number of stolons (ST1), the highest node of stolon emergence (SEM1), the number of rhizomes (RH1), the proportion of stems that were rhizomes (PR1), and the highest node of rhizome emergence (REM1). On the day of first flower, without unpotting the plants, we scored flowering time (FT), the node of the first flower (FN), corolla tube length (CTL), corolla limb length (CLL), corolla limb width (CLW), leaf length (LL) and width (LW) on the first pair of true leaves, internode length (IL; measured between the 1st and 2nd true leaves), internode width (IW), as well as repeatedly measuring traits from our first developmental stage (i.e. ST2, SEM2, RH2, PR2, and REM2). During the ’late reproductive’ period, we destructively sampled plants by unpotting them, washing soil and debris from belowground biomass. We then measured the same set of traits we had measured in the first and second developmental stage (i.e. ST3, SEM3, RH3, REM3, PR3), as well as several more detailed aspects of stolon and rhizome development, measured on the longest stolon or rhizome. These include: stolon length (SL), stolon width (SW), the number of stolon nodes (SN), the number of stolon branches (SB), average branches per stolon node (SBA), the maximum stolon branches per node (MSBR), stolon leaf length (SLL; measured as the first pair of proximal leaves), stolon leaf width (SLW); rhizome length (RL), rhizome width (RW), the number of rhizome nodes (RN), the number of rhizome branches (RB), average branches per rhizome node (RBA), the maximum rhizome branches per node (MRBR), rhizome leaf length (RLL; measured as the first pair of proximal leaves), and rhizome leaf width (RLW).

In total, we scored 40 traits across three time points. Notably, we did not unpot the plants when evaluating whether rhizomes were produced during the ’vegetative’ and ’early reproductive’ stages, rather we carefully dug a small portion of the soil to expose a small part of the rhizome and recovered the exposed area with soil after scoring. Visual representations of all traits are provided in Supplementary Fig. S7.

### Trait heritability, dominance, and correlation structure

We first sought to understand aspects of genetic architecture and modularity that can be assessed strictly through looking at phenotypes. We leveraged the phenotypic variation (σ2) among replicate F1 versus F2 hybrids to estimate broad-sense heritability (*h^2^*) for each trait, calculated as *h^2^= [σ^2^(F2s) - mean (σ^2^(F1s), σ^2^(P1s), σ^2^(P2s))] / σ^2^(F2s)* and also calculated dominance coefficients (*d*) for each trait as *d*= (*ZPARENT1* - *ZF1*) / (*ZPARENT1 - ZPARENT2*), where *Z* denotes the phenotypic mean for that genotype.

To determine modules that we could compare across mapping populations, we adopted a common approach in evolutionary biology by defining putative trait modules *a priori* based on their functional roles (Diggle 2014; Melo et al. 2016; Chen et al. 2025), classifying them into five categories: floral traits, phenological traits, size traits, stolon traits, and rhizome traits (Table S1). These modules were confirmed by computing the pairwise correlation coefficients for all traits, and assessing whether these predefined modules corresponded to highly correlated groups of traits using a pairwise Spearman’s correlation coefficients (*r*) (using the *psych::corr.test* function in R). We then calculated the mean correlation coefficients within and between modules. We tested whether the extent of within-module correlation was greater than the between-modules correlation in two ways. First, we performed a linear regression with the absolute correlation coefficient as the response variable and module type (i.e. floral, phenology, stolon, rhizome, or size), module status (i.e. within module versus between module), and their interaction as independent variables. We then assessed the significance of these factors using a Type III Anova (*Car::Anova* function in R), and assessed whether within versus between module correlation coefficients significantly differed for each category in each cross using the *Emmeans::emmeans* in R. Second, we performed 1000 permutations in which traits were randomly assigned to different modules and assessed whether correlation coefficients were higher for pre-defined modules than randomly selected groups of traits.

To determine whether modularity changed across development, we leveraged a subset of stolon-related and rhizome-related traits that were phenotyped at each of the 3 developmental stages (i.e. ST1, ST2, ST3, SEM1, SEM2, and SEM3 for stolon-related traits and RH1, RH2, RH3, REM1, REM2, and REM3 for rhizome-related traits). We then compared the correlation coefficient of stolon-related or rhizome-related traits across different stages. We used a linear regression to determine whether correlation coefficients within stolon- or rhizome-related traits differed across developmental stages, using the absolute correlation coefficient as the response variable, and the developmental stage, trait category (i.e. rhizome or stolon), and their interaction as independent variables. We assessed significance of these variables using a Type III Anova, and the significance of specific contrasts using the estimated marginal means, as above. Additionally, we performed a 1000 permutations of traits across development to determine if traits measured at the same developmental stage were more correlated than traits measured between developmental stages for rhizome and stolon-related traits separately.

Lastly, to determine whether populations derived from more closely related taxa exhibit more similar phenotypic correlation patterns, we conducted a Mantel test (Mantel 1967) to analyze the relationships among the correlation matrices of the four populations. We then completed a linear regression with the Mantel test correlation coefficient as the response variable and genomic divergence (DXY; see below for description of calculation) as the independent variable.

### DNA extraction, library construction and sequencing

We extracted DNA from flower buds using a modified CTAB method (Doyle and Doyle 1987; dx.doi.org/10.17504/protocols.io.3rrgm56). We constructed ddRADseq libraries following the BestRAD method (https://doi.org/10.17504/protocols.io.6awhafe) for all F2s, F1s, and each parental line. In brief, approximately 100 ng of genomic DNA from each individual sample was digested with BfaI and PstI enzymes and then ligated to a unique in-line barcode adapter. We pooled 48 samples per library, which were prepared using NEBNext Ultra II library preparation kits for Illumina (New England BioLabs, Ipswich, MA). Each library was indexed with an i5 universal adapter containing a molecular barcode and a unique NEBNext i7 adapter. The adapter-ligated fragments were then enriched through 12 cycles of PCR. Following library preparation, the libraries were size-selected to a range of 250–850 bp using BluePippin 1.5% agarose cassettes (Sage Science, Beverly, MA). We prepared a total of 45 libraries from 4 crosses (2,160 individuals) and generated 150bp paired-end reads using a NovaSeq xPlus machine at the Yale Center for Genome Analysis.

### Sequence processing, divergence estimation, and linkage map construction

To process our sequence data, we removed PCR duplicates with a custom Python script (https://doi.org/10.17504/protocols.io.bjnbkman), demultiplexed reads using process_radtags program in the Stacks pipeline (v2.59) (Catchen et al. 2013), trimmed Illumina adaptors using Trimmomatic (v0.39) (Bolger et al. 2014), and aligned reads reads to the *Mimulus guttatus* IM62 v3 reference genome (Lovell et al. 2025) using BWA-MEM (version 0.7.17) (Li and Durbin 2009). We then used SAMtools (v1.20) (Li et al. 2009) to sort and filter reads with alignment qualities below Q30. We used *mpileup* in BCFtools to call genotypes (v1.16) (Danecek et al. 2021). We applied a filtering process to the resulting VCFs to retain only biallelic single nucleotide polymorphisms (SNPs) meeting the following criteria: a mapping quality (MQ) above 20, a minimum of 50 effective samples, an allele frequency ranging between 0.2 and 0.8, a median read depth below 50 across samples, and a minimum spacing of 50 base pairs between SNPs within each scaffold to prevent dense clustering.

Following SNP calling and filtering, we used genotypes derived from individual SNPs to assign preliminary genotypes within 100KB windows for each individual. A preliminary windowed genotype was assigned if the window contained at least 1 SNP and 5 reads; it was considered homozygous if 95% or more of the reads aligned with a single parent, and heterozygous if 90% or fewer reads aligned with a single parent. We then employed the Genome Order Optimization by Genetic Algorithm (GOOGA) (Flagel et al. 2019) to estimate individual genotype error rates by evaluating the probability of erroneously identifying a homozygous genotype as heterozygous, the probability of erroneously identifying a homozygous genotype as the alternative homozygous genotype, and the probability of erroneously identifying a heterozygous genotype as homozygous. To ensure the use of precise genotypes in constructing the linkage map, we excluded any individual with an error rate over 20% in any of the three types. Then we used these filtered individuals to perform GOOGA to calculate recombination rates among markers and removed poor or misplaced markers based on the rate results (e.g., markers that exhibited a recombination rate of 0.25 or greater). The process of calculation and removal is repeated iteratively until all the remaining markers exhibit reasonable recombination rates. Based on the polished genetic markers, we reevaluated individual genotype errors and continued to exclude any individual with over 20% error. We then recalculated recombination rates among markers using GOOGA, resulting in the final linkage map output. This resulted in 464, 418, 539, and 23 individuals (the latter being very small due to significant segregation distortion; Fig. S3), and 1622, 1695, 1711, and 1731 markers being used to construct the linkage map for the IMPO × OPB, CWF × OPB, LVR × OPB, and EAM × OPB populations, respectively (Fig. S8). Finally, we estimated the genotyping error rates for all individuals (even those with higher error rates) based on the recombination rates defined by the linkage map and then used GOOGA to estimate individual genotype posterior probabilities at each genetic marker. We called final genotypes as those that resulted in a posterior probability of 90% or higher.

We estimated pairwise nucleotide divergence (DXY) among taxa using *pixy* v2.0.0.beta8 (Korunes and Samuk 2021). Briefly, we generated a multi-sample VCF of the parental lines (LVR, CWF, EAM, IMPO, and OPB) from whole-genome aligned BAM files using bcftools mpileup and bcftools call (BCFtools v1.16; Danecek et al. 2021), retaining both variant and invariant sites. We indexed the resulting VCF using Tabix v0.2.6. We restricted sites to fourfold degenerate sites to best estimate neutral divergence, then calculated DXY using sliding windows of 2,500 bp. We obtained a genome-wide DXY estimate by computing the weighted mean of avg_dxy, using count_comparisons as weights to account for variation in data availability across windows.

### QTL mapping and QTL overlap

We performed QTL analysis in the *R/qtl* package (Broman et al. 2003; Arends et al. 2010) using the Haley-Knott regression method (Haley and Knott 1992). We determined significance thresholds of the LOD score (*p* = 0.01) through 1000 permutations. For traits with more than one significant QTL identified, and assessed pairwise non-additive interactions using the *addint* function in the *R/qtl* package. We computed the average additive effects (*a*), dominance effects (*d*), and the percentage of phenotypic variation explained (*r^2^*) by each QTL among F2s using the *makeqtl()* and *fitqtl()* functions in the *R/qtl* package. Additionally, we defined the 1.5 LOD-drop confidence intervals for each QTL using the *lodint()* function within the *R/qtl* package.

The degree of QTL overlap between each pair of traits within and between populations was determined using the Jaccard index, following (Liao et al. 2022; Chen et al. 2025). We calculated the means (and standard errors) of QTL overlap within and between trait modules for each population. To test whether the within-module overlap was greater than the between-module overlap, we performed a linear regression with overlap as the response variable, module type (i.e. floral, phenological, stolon, rhizome, and size), module status (i.e. within versus between), and their interaction as independent variables for each cross separately. We assessed significance of these variables using a Type III Anova (*Car::Anova* in R) and determined significance of specific contrasts (i.e. within versus between modules) for each module type using the *Emmeans::emmeans* function in R. Additionally, we performed 1000 permutations in which traits were randomly assigned to different modules to determine if QTL overlap was higher than expected by chance.

For each cross, we also assessed whether the extent of QTL overlap changed across development for stolon- and rhizome-related traits. As in above, we categorized stolon- and rhizome-related traits that were measured at each developmental stage. We asked whether overlap for a given trait changed across development using a linear regression with overlap as the response variable and developmental transition (i.e. vegetative to early reproductive stages and early-reproductive to late reproductive stages as the levels), trait (i.e. number of stolons, number of rhizomes, the highest node at which stolons emerge, and the highest node at which rhizomes emerge), and cross as independent variables. We assessed the significance of these variables using a Type III Anova as above. In this case, overlap differed only across developmental transitions (development: *F*=10.69, *p*=0.0067; trait: *F*=1.55, *p*=0.68; and cross: *F*=1.5, *p*=0.27), and so no further contrasts were calculated.

Lastly, to examine the genetic parallelism of the same traits across different populations, we calculated the degree of QTL overlap for each trait between each pair of crosses using Jaccard indices. We evaluated the significance of the observed overlap between QTLs by performing 1000 permutations, in which QTLs were randomly reassigned to different traits. Lastly, we assessed whether the extent of QTL overlap declined with divergence time with a linear regression using QTL overlap as the response variable and DXY, trait-module, and their interaction as independent variables. We assessed the significance of each of these independent variables using a Type III Anova, as above.

### Assessing the factors that influence individual QTL overlap

We next sought to understand if certain aspects of genetic architecture or divergence influenced the probability that individual QTLs overlap across the four F2 mapping populations. We defined QTL overlap as the presence of a QTL associated with the same trait on the same chromosome in another population, with overlapping genomic intervals (i.e., Mbmin < Mbmax and Mbmax > Mbmin). For each factor, we constructed a logistic regression with the overlap for each QTL as a binary response variable (1= overlapping, 0= no overlap), and aspects of genetic architecture (i.e. a QTL’s additive effect (*a*, standardized by the difference between parental lines), dominance deviation (*d/a*), effect size (*r²*)), trait module, and their interaction as the independent variables. In each case, we used a binomial generalized linear model (GLM) using the *glm* function in R with family=binomial. For aspects of genetic architecture that we coded as binary (i.e. whether a QTL was implicated in more than one trait (out measure of pleiotropy), and whether a QTL had an epistatic effect), we used a Fisher’s Exact test, comparing the count of QTLs that were/were not identified in more than one cross for each pleiotropic or epistatic status.

## Acknowledgements

We are grateful to Christopher Bolick and Nathan Guzzo for plant care in the Yale Science Building (YSB) Greenhouse. We thank Ben Eissler, Hagar Soliman, and Dr. Megan Frayer for help with the phenotyping. We appreciate TJ Johnson’s support in library preparation. We are grateful to all members of the Coughlan lab for helpful feedback on earlier drafts of this manuscript. This work was supported by an NIH grant to J.M.C. (NIH R35GM150907), Yale’s Brown endowed fellowship which supported H.C. for one year, and the STARs program which supported P.J. for one summer.

## Author contributions

H.C. and J.M.C. planned and designed the research. H.C. performed the experiments and generated the data. P.J. helped with data collection. H.C. and J.M.C. analyzed and interpreted the data, visualized the results, and drafted the manuscript. All authors approved the manuscript before submission.

