## supplemental tables for "Unique genetic bases of repeated life-history divergence associated with high altitude adaptation in *Mimulus* perennials"

**Table S1. Variation in life-history traits (means ± SD) for *M. tilingii*, CWF, *M. corallinus*, *M. decorus*, the coastal *M. guttatus*, and their $F_1$ and $F_2$ hybrids.** A) *M. tilingii*, the coastal *M. guttatus*, and their $F_1$ and $F_2$ hybrids. B) CWF, the coastal *M. guttatus*, and their $F_1$ and $F_2$ hybrids. C) *M. corallinus*, the coastal *M. guttatus*, and their $F_1$ and $F_2$ hybrids. D) *M. decorus*, the coastal *M. guttatus*, and their $F_1$ and $F_2$ hybrids. Broad-sense heritability ($h^2$) and dominance coefficients ($d$) were calculated following Coughlan et al. (2021).

**A**

| Trait | *M. tilingii* LVR (n=23) | *M. guttatus* OPB (n=24) | F1 hybrids (n=22) | F2 hybrids (n=543) | *d* | $h^2$ |
|---|---|---|---|---|---|---|
| **Floral traits** | | | | | | |
| Corolla tube length (CTL) | 14.68 ± 0.9 | 21.66 ± 0.85 | 19.88 ± 2.13 | 19.98 ± 2.6 | 0.74 | 0.71 |
| Corolla limb length (CLL) | 20.53 ± 1.81 | 21.11 ± 2.02 | 27.13 ± 4.31 | 23.67 ± 4.51 | 11.4 | 0.59 |
| Corolla limb width (CLW) | 21.91 ± 2.15 | 20.57 ± 2.06 | 24.3 ± 3.77 | 23.28 ± 3.94 | -1.79 | 0.53 |
| **Phenological traits** | | | | | | |
| Flower node (FN) | 4 ± 0.75 | 5.63 ± 0.49 | 3.41 ± 0.5 | 4.44 ± 1.55 | -0.36 | 0.86 |
| Flower time (FT) | 27.5 ± 2.24 | 40.67 ± 1.43 | 26.14 ± 4.69 | 31.15 ± 7.45 | -0.1 | 0.83 |
| Internode length (IL) | 6.4 ± 2.22 | 11.02 ± 3.43 | 14.76 ± 10.21 | 12.77 ± 8.26 | 1.81 | 0.43 |
| **Size traits** | | | | | | |
| Leaf length (LL) | 18.33 ± 4.69 | 30.93 ± 4.26 | 36.21 ± 11.23 | 35.58 ± 11.61 | 1.42 | 0.61 |
| Leaf width (LW) | 10.16 ± 2.36 | 29.27 ± 6.2 | 22.28 ± 6.75 | 22.11 ± 7.55 | 0.63 | 0.50 |
| Internode width (IW) | 2.28 ± 0.3 | 6.14 ± 0.78 | 3.91 ± 0.62 | 4.39 ± 1.07 | 0.42 | 0.70 |
| **Stolon traits** | | | | | | |
| Stolon number 1 (ST1) | 3.78 ± 1.31 | 0.79 ± 0.98 | 3.09 ± 2.07 | 2.88 ± 1.75 | 0.23 | 0.27 |
| Stolon number 2 (ST2) | 5.73 ± 1.42 | 4.92 ± 1.28 | 5.77 ± 2.05 | 5.67 ± 2.32 | -0.06 | 0.54 |
| Stolon number 3 (ST3) | 6.16 ± 1.3 | 6.58 ± 1.32 | 7.3 ± 2.32 | 7.09 ± 2.51 | 2.68 | 0.56 |
| Emergence node 1 (SEM1) | 1.91 ± 0.29 | 1 ± 0 | 1.72 ± 0.57 | 1.55 ± 0.57 | 0.21 | 0.59 |
| Emergence node 2 (SEM2) | 2.27 ± 0.46 | 2.58 ± 0.5 | 2.23 ± 0.43 | 2.49 ± 0.9 | -0.15 | 0.74 |
| Emergence node 3 (SEM3) | 2.35 ± 0.49 | 3.29 ± 0.69 | 2.55 ± 0.6 | 2.89 ± 0.9 | 0.21 | 0.58 |
| Stolon length (SL) | 19.1 ± 3.53 | 54.52 ± 10.61 | 43.39 ± 6.14 | 41.22 ± 11.01 | 0.69 | 0.57 |
| Stolon width (SW) | 1.73 ± 0.29 | 4.04 ± 0.81 | 2.55 ± 0.38 | 2.75 ± 0.61 | 0.35 | 0.23 |
| Stolon node (SN) | 12.42 ± 2.36 | 13.25 ± 2.19 | 16.33 ± 3.71 | 13.86 ± 3.66 | 4.72 | 0.43 |
| Stolon branch (SB) | 4.5 ± 1.51 | 6.39 ± 1.23 | 4.33 ± 1.59 | 5.43 ± 2.62 | -0.09 | 0.71 |
| Stolon branch/node (SBA) | 0.43 ± 0.19 | 0.49 ± 0.11 | 0.28 ± 0.08 | 0.43 ± 0.26 | -2.2 | 0.74 |
| Maximum branch (MSBR) | 2.2 ± 0.42 | 2.26 ± 0.54 | 2.93 ± 0.96 | 2.49 ± 0.74 | 12.05 | 0.21 |
| Stolon leaf length (SLL) | NA | 43.57 ± 8.55 | 53.37 ± 9.68 | 42.25 ± 15.88 | 1.23 | 0.72 |
| Stolon leaf width (SLW) | NA | 31.68 ± 7.12 | 30.01 ± 4.13 | 24.84 ± 9.86 | 0.95 | 0.70 |
| **Rhizome traits** | | | | | | |
| Rhizome number 1 (RH1) | 0.52 ± 0.79 | 0 | 0.09 ± 0.29 | 0.1 ± 0.34 | 0.83 | 0 |
| Rhizome number 2 (RH2) | 1.18 ± 0.73 | 0 | 1.23 ± 0.81 | 0.6 ± 0.8 | -0.04 | 0.41 |
| Rhizome number 3 (RH3) | 1.67 ± 1.08 | 0 | 1.95 ± 0.92 | 0.74 ± 0.91 | -0.17 | 0.23 |
| Emergence node 1 (REM1) | 1 ± 0 | NA | 1 ± 0 | 1 ± 0 | NA | NA |
| Emergence node 2 (REM2) | 1.06 ± 0.24 | NA | 1.06 ± 0.24 | 1.08 ± 0.27 | NA | 0.51 |
| Emergence node 3 (REM3) | 1 ± 0 | NA | 1.21 ± 0.42 | 1.12 ± 0.35 | NA | NA |
| Rhizome length (RL) | 1.52 ± 1.96 | NA | 18.53 ± 13.02 | 8.59 ± 11.13 | NA | NA |
| Rhizome width (RW) | 1.06 ± 0.16 | NA | 2.35 ± 0.59 | 1.89 ± 0.57 | NA | NA |
| Rhizome node (RN) | 3.08 ± 1.31 | NA | 7.43 ± 3.34 | 4.32 ± 3.07 | NA | NA |
| Rhizome branch (RB) | 4.75 ± 3.14 | NA | 7.86 ± 2.96 | 4.52 ± 4.19 | NA | NA |
| Rhizome branch/node (RBA) | 1.6 ± 0.9 | NA | 1.18 ± 0.45 | 1.01 ± 0.64 | NA | NA |
| Maximum branch (MRBR) | 2.08 ± 1.08 | NA | 2.14 ± 0.53 | 1.77 ± 0.9 | NA | NA |
| Rhizome leaf length (RLL) | NA | NA | 2.53 ± 0.86 | 3.15 ± 1.96 | NA | NA |
| Rhizome leaf width (RLW) | NA | NA | 1.63 ± 0.5 | 2.06 ± 1.35 | NA | NA |
| Rhizome proportion 1 (PR1) | 0.1 ± 0.15 | 0 | 0.06 ± 0.23 | 0.02 ± 0.08 | 0.38 | 0 |
| Rhizome proportion 2 (PR2) | 0.18 ± 0.11 | 0 | 0.2 ± 0.16 | 0.1 ± 0.13 | -0.11 | 0.31 |
| Rhizome proportion 3 (PR3) | 0.2 ± 0.12 | 0 | 0.22 ± 0.11 | 0.09 ± 0.12 | -0.12 | 0.39 |

**B**

| Trait | CWF (n=15) | *M. guttatus* OPB (n=18) | F1 hybrids (n=41) | F2 hybrids (n=430) | *d* | $h^2$ |
|---|---|---|---|---|---|---|
| **Floral traits** | | | | | | |
| Corolla tube length (CTL) | 12.93 ± 1.34 | 21.93 ± 1.23 | 18.87 ± 1.01 | 18.91 ± 2.35 | 0.66 | 0.75 |
| Corolla limb length (CLL) | 18.39 ± 1.99 | 22.45 ± 2.01 | 23.56 ± 2.52 | 21.37 ± 3.13 | 1.28 | 0.54 |
| Corolla limb width (CLW) | 19.39 ± 3.42 | 21.81 ± 1.82 | 23.7 ± 2.07 | 22.72 ± 3.67 | 1.78 | 0.57 |
| **Phenological traits** | | | | | | |
| Flower node (FN) | 5.13 ± 0.99 | 5.65 ± 0.86 | 5.15 ± 0.77 | 5.84 ± 1.25 | 0.03 | 0.53 |
| Flower time (FT) | 29.43 ± 4.67 | 43.67 ± 3.36 | 30.39 ± 4.06 | 35.48 ± 6.86 | 0.07 | 0.67 |
| Internode length (IL) | 10.89 ± 9.57 | 14.44 ± 7.01 | 13.87 ± 9.42 | 11.15 ± 8.69 | 0.84 | 0.03 |
| **Size traits** | | | | | | |
| Leaf length (LL) | 22.35 ± 7.58 | 29.17 ± 5.16 | 32.9 ± 11.69 | 28.11 ± 9.62 | 1.55 | 0.24 |
| Leaf width (LW) | 20.2 ± 5.73 | 26.35 ± 4.63 | 31.13 ± 11.41 | 25.47 ± 9.3 | 1.78 | 0.31 |
| Internode width (IW) | 3.36 ± 0.63 | 5.97 ± 0.64 | 5.4 ± 0.75 | 5.04 ± 1.01 | 0.78 | 0.57 |
| **Stolon traits** | | | | | | |
| Stolon number 1 (ST1) | 5.13 ± 2.39 | 1.22 ± 0.94 | 3.73 ± 1.91 | 3.13 ± 2.48 | 0.36 | 0.47 |
| Stolon number 2 (ST2) | 10.13 ± 3.07 | 4.41 ± 1.42 | 9.05 ± 2.87 | 9.11 ± 3.47 | 0.19 | 0.48 |
| Stolon number 3 (ST3) | 15.6 ± 3.78 | 5.65 ± 1.69 | 15 ± 2.03 | 12.63 ± 4.24 | 0.06 | 0.64 |
| Emergence node 1 (SEM1) | 1.73 ± 0.46 | 1 ± 0 | 1.47 ± 0.51 | 1.52 ± 0.53 | 0.35 | 0.47 |
| Emergence node 2 (SEM2) | 2.73 ± 0.8 | 2.71 ± 0.47 | 2.63 ± 0.7 | 3.03 ± 0.98 | 3.95 | 0.55 |
| Emergence node 3 (SEM3) | 4.9 ± 0.99 | 3.35 ± 0.7 | 4.48 ± 0.71 | 4.27 ± 1.09 | 0.27 | 0.48 |
| Stolon length (SL) | 30.5 ± 7.02 | 45.65 ± 8.18 | 55.82 ± 7.04 | 44.87 ± 12.45 | 1.67 | 0.67 |
| Stolon width (SW) | 2 ± 0.49 | 4.1 ± 0.61 | 2.91 ± 0.52 | 2.67 ± 0.61 | 0.43 | 0.25 |
| Stolon node (SN) | 12.33 ± 2.12 | 12.24 ± 1.95 | 14 ± 2.02 | 11.69 ± 2.75 | -17 | 0.49 |
| Stolon branch (SB) | 21.33 ± 5.68 | 6.47 ± 1.81 | 10.81 ± 3.44 | 11.9 ± 5.1 | 0.71 | 0.44 |
| Stolon branch/node (SBA) | 1.76 ± 0.48 | 0.55 ± 0.18 | 0.79 ± 0.28 | 1.05 ± 0.53 | 0.80 | 0.64 |
| Maximum branch (MSBR) | 5.56 ± 1.13 | 2.24 ± 0.56 | 3.81 ± 0.54 | 3.68 ± 0.84 | 0.52 | 0.20 |
| Stolon leaf length (SLL) | 23.67 ± 5.95 | 45.07 ± 12.49 | 37.42 ± 14.43 | 34.57 ± 14.68 | 0.64 | 0.42 |
| Stolon leaf width (SLW) | 20.78 ± 3.1 | 29.74 ± 9.75 | 27.95 ± 10.37 | 24.42 ± 9.39 | 0.80 | 0.25 |
| **Rhizome traits** | | | | | | |
| Rhizome number 1 (RH1) | 0.47 ± 0.52 | 0 | 0.07 ± 0.26 | 0.09 ± 0.34 | 0.84 | 0.08 |
| Rhizome number 2 (RH2) | 0.87 ± 1.06 | 0 | 0.28 ± 0.55 | 0.38 ± 0.73 | 0.68 | 0.14 |
| Rhizome number 3 (RH3) | 3.25 ± 2.49 | 0 | 0.75 ± 0.94 | 0.92 ± 1.28 | 0.77 | 0 |
| Emergence node 1 (REM1) | 1 ± 0 | NA | 1 ± 0 | 1.03 ± 0.18 | NA | NA |
| Emergence node 2 (REM2) | 1.11 ± 0.33 | NA | 1.11 ± 0.33 | 1.3 ± 0.5 | NA | NA |
| Emergence node 3 (REM3) | 2.08 ± 0.67 | NA | 1.68 ± 0.58 | 1.61 ± 0.67 | NA | NA |
| Rhizome length (RL) | 7.24 ± 2.84 | NA | 10.24 ± 12.69 | 6.5 ± 8.96 | NA | NA |
| Rhizome width (RW) | 1.33 ± 0.38 | NA | 1.74 ± 0.43 | 1.65 ± 0.48 | NA | NA |
| Rhizome node (RN) | 9.11 ± 3.48 | NA | 5.72 ± 3.14 | 4.66 ± 3.03 | NA | NA |
| Rhizome branch (RB) | 17.67 ± 8.08 | NA | 7.83 ± 6.01 | 5.56 ± 5.69 | NA | NA |
| Rhizome branch/node (RBA) | 1.97 ± 0.57 | NA | 1.18 ± 0.55 | 0.93 ± 0.68 | NA | NA |
| Maximum branch (MRBR) | 3.33 ± 1.41 | NA | 2.06 ± 1.11 | 1.63 ± 1.18 | NA | NA |
| Rhizome leaf length (RLL) | NA | NA | 4.19 ± 2.91 | 2.7 ± 2.47 | NA | NA |
| Rhizome leaf width (RLW) | NA | NA | 3.31 ± 2.1 | 2.26 ± 2.21 | NA | NA |
| Rhizome proportion 1 (PR1) | 0.08 ± 0.1 | 0 | 0.02 ± 0.07 | 0.02 ± 0.09 | 0.77 | 0.45 |
| Rhizome proportion 2 (PR2) | 0.07 ± 0.08 | 0 | 0.03 ± 0.07 | 0.04 ± 0.08 | 0.57 | 0.46 |
| Rhizome proportion 3 (PR3) | 0.17 ± 0.13 | 0 | 0.05 ± 0.06 | 0.06 ± 0.09 | 0.71 | 0.22 |

C

| Trait | *M. corallinus* EAM (n=23) | *M. guttatus* OPB (n=36) | F1 hybrids (n=2) | F2 hybrids (n=582) | *d* | $h^2$ |
|---|---|---|---|---|---|---|
| **Floral traits** | | | | | | |
| Corolla tube length (CTL) | 16.05±1.14 | 21.71±1.50 | 19.82±1.60 | 20.30±1.31 | 0.67 | 0.11 |
| Corolla limb length (CLL) | 24.19±3.61 | 20.70±2.55 | 30.40±1.19 | 28.64±3.17 | -1.78 | 0.37 |
| Corolla limb width (CLW) | 21.53±3.00 | 21.71±2.34 | 31.08±0.55 | 33.45±2.83 | 54.69 | 0.43 |
| **Phenological traits** | | | | | | |
| Flower node (FN) | 7.50±0.86 | 6.50±1.18 | 6.00±1.41 | 6.20±0.93 | 1.50 | 0 |
| Flower time (FT) | 36.76±3.95 | 46.91±4.51 | 34.50±4.95 | 33.90±3.59 | -0.22 | 0 |
| Internode length (IL) | 8.40±1.89 | 25.87±15.42 | 14.14±6.72 | 13.33±5.48 | 0.33 | 0 |
| **Size traits** | | | | | | |
| Leaf length (LL) | 35.65±5.91 | 26.94±3.00 | 57.02±NA | 44.68±12.72 | -2.45 | 0.92 |
| Leaf width (LW) | 22.42±4.79 | 26.05±3.35 | 41.72±NA | 35.27±8.63 | 5.31 | 0.86 |
| Internode width (IW) | 3.65±0.61 | 5.81±0.52 | 5.54±0.57 | 5.75±0.65 | 0.87 | 0.41 |
| **Stolon traits** | | | | | | |
| Stolon number 1 (ST1) | 3.48±1.95 | 1.31±0.95 | 4.00±1.41 | 4.09±1.33 | -0.24 | 0 |
| Stolon number 2 (ST2) | 14.56±4.15 | 4.56±2.10 | 7.50±2.12 | 8.03±2.07 | 0.71 | 0 |
| Stolon number 3 (ST3) | 21.47±5.67 | 6.35±2.26 | 13.00±7.07 | 9.48±2.39 | 0.56 | 0 |
| Emergence node 1 (SEM1) | 3.16±0.50 | 1±0 | 2.50±0.71 | 2.15±0.44 | 0.30 | 0.18 |
| Emergence node 2 (SEM2) | 6.56±0.86 | 2.6±0.96 | 4.00±1.41 | 3.75±0.74 | 0.65 | 0 |
| Emergence node 3 (SEM3) | 9.47±2.21 | 3.46±0.95 | 5.50±2.12 | 4.26±0.82 | 0.66 | 0 |
| Stolon length (SL) | 30.25±6.52 | 52.67±8.28 | 59.90±1.56 | 50.38±11.82 | 1.32 | 0.74 |
| Stolon width (SW) | 1.67±0.32 | 3.55±0.41 | 2.86±0.94 | 2.82±0.43 | 0.63 | 0 |
| Stolon node (SN) | 10.94±2.41 | 13.35±2.56 | 14.00±0.00 | 13.99±2.60 | 1.27 | 0.42 |
| Stolon branch (SB) | 18.31±11.26 | 5.81±1.27 | 14.50±6.36 | 12.38±4.38 | 0.30 | 0 |
| Stolon branch/node (SBA) | 1.55±0.68 | 0.45±0.14 | 1.04±0.45 | 0.90±0.31 | 0.47 | 0 |
| Maximum branch (MSBR) | 3.20±0.86 | 2.08±0.27 | 3.00±1.41 | 2.31±0.62 | 0.18 | 0 |
| Stolon leaf length (SLL) | 43.38±13.25 | 37.78±9.12 | NA | 35.96±11.20 | NA | NA |
| Stolon leaf width (SLW) | 29.13±9.25 | 27.84±7.72 | NA | 23.49±6.72 | NA | NA |
| **Rhizome traits** | | | | | | |
| Rhizome number 1 (RH1) | 3.00±2.28 | 0 | 0.50±0.71 | 1.22±0.95 | 0.83 | 0 |
| Rhizome number 2 (RH2) | 8.11±3.10 | 0 | 2.00±2.83 | 2.26±1.10 | 0.75 | 0 |
| Rhizome number 3 (RH3) | 13.58±3.61 | 0 | 4.50±3.54 | 3.03±1.33 | 0.67 | 0 |
| Emergence node 1 (REM1) | 1.89±0.81 | NA | NA | 1.07±0.25 | NA | NA |
| Emergence node 2 (REM2) | 3.33±1.08 | NA | NA | 1.53±0.51 | NA | NA |
| Emergence node 3 (REM3) | 3.82±1.33 | NA | 1.50±0.71 | 1.79±0.60 | NA | NA |
| Rhizome length (RL) | 15.78±3.78 | NA | 22.05±6.29 | 19.13±10.67 | NA | NA |
| Rhizome width (RW) | 1.31±0.25 | NA | 2.41±0.33 | 1.78±0.64 | NA | NA |
| Rhizome node (RN) | 11.33±1.91 | NA | 12.50±3.54 | 8.64±2.46 | NA | NA |
| Rhizome branch (RB) | 18.06±3.61 | NA | 17.00±2.83 | 9.59±4.19 | NA | NA |
| Rhizome branch/node (RBA) | 1.62±0.32 | NA | 1.38±0.16 | 1.11±0.41 | NA | NA |
| Maximum branch (MRBR) | 3.67±1.08 | NA | 3.00±0.00 | 2.00±0.47 | NA | NA |
| Rhizome leaf length (RLL) | 2.84±2.14 | NA | NA | 3.91±3.55 | NA | NA |
| Rhizome leaf width (RLW) | 1.34±0.44 | NA | NA | 3.46±3.01 | NA | NA |
| Rhizome proportion 1 (PR1) | 0.45±0.23 | 0 | 0.08±0.12 | 0.22±0.18 | 0.81 | 0.37 |
| Rhizome proportion 2 (PR2) | 0.37±0.14 | 0 | 0.20±0.28 | 0.22±0.10 | 0.45 | 0 |
| Rhizome proportion 3 (PR3) | 0.39±0.11 | 0 | 0.24±0.06 | 0.24±0.10 | 0.38 | 0.58 |

**D**

| Trait | *M. decorus* IMPO (n=26) | *M. guttatus* OPB (n=34) | F1 hybrids (n=21) | F2 hybrids (n=469) | *d* | $h^2$ |
|---|---|---|---|---|---|---|
| **Floral traits** | | | | | | |
| Corolla tube length (CTL) | 24.47±1.63 | 20.98±1.32 | 24.69±0.79 | 22.46±2.58 | -0.06 | 0.76 |
| Corolla limb length (CLL) | 22.12±1.75 | 19.90±2.75 | 22.45±3.15 | 20.43±3.19 | -0.15 | 0.36 |
| Corolla limb width (CLW) | 22.63±3.05 | 20.79±2.65 | 24.27±1.37 | 22.74±3.28 | -0.89 | 0.46 |
| **Phenological traits** | | | | | | |
| Flower node (FN) | 5.52±0.67 | 6.09±1.09 | 5.40±0.75 | 6.31±1.40 | -0.21 | 0.64 |
| Flower time (FT) | 33.50±4.27 | 44.28±4.91 | 32.50±3.93 | 37.34±7.15 | -0.09 | 0.64 |
| Internode length (IL) | 8.55±4.16 | 9.87±9.09 | 9.61±3.53 | 8.63±5.24 | 0.80 | 0 |
| **Size traits** | | | | | | |
| Leaf length (LL) | 26.71±6.97 | 24.59±5.64 | 38.38±7.51 | 31.56±9.85 | -5.50 | 0.55 |
| Leaf width (LW) | 15.10±3.62 | 21.31±6.79 | 29.64±6.18 | 23.42±6.84 | 2.34 | 0.34 |
| Internode width (IW) | 3.40±0.51 | 5.79±0.63 | 6.44±1.19 | 5.61±1.31 | 1.27 | 0.62 |
| **Stolon traits** | | | | | | |
| Stolon number 1 (ST1) | 2.88±1.70 | 1.06±0.95 | 4.52±2.02 | 3.35±2.17 | -0.90 | 0.46 |
| Stolon number 2 (ST2) | 7.46±2.19 | 4.94±1.50 | 9.60±1.98 | 9.21±3.38 | -0.85 | 0.69 |
| Stolon number 3 (ST3) | 6.59±2.27 | 6.17±1.53 | 11.40±2.50 | 10.34±3.81 | -11.41 | 0.70 |
| Emergence node 1 (SEM1) | 1.83±0.38 | 1±0 | 1.85±0.37 | 1.77±0.47 | -0.02 | 0.59 |
| Emergence node 2 (SEM2) | 4.08±0.83 | 2.91±0.78 | 3.05±0.39 | 3.38±1.00 | 0.88 | 0.53 |
| Emergence node 3 (SEM3) | 4.67±1.08 | 3.43±0.86 | 3.87±0.74 | 3.90±1.23 | 0.65 | 0.49 |
| Stolon length (SL) | 37.63±10.16 | 56.08±6.78 | 70.22±7.50 | 53.76±12.13 | 1.77 | 0.56 |
| Stolon width (SW) | 1.94±0.42 | 3.59±0.51 | 2.98±0.41 | 2.74±0.56 | 0.63 | 0.40 |
| Stolon node (SN) | 11.20±2.18 | 12.17±1.98 | 15.00±2.63 | 12.40±2.98 | 3.91 | 0.45 |
| Stolon branch (SB) | 13.14±6.02 | 5.44±1.58 | 12.67±3.03 | 9.83±4.37 | 0.06 | 0.22 |
| Stolon branch/node (SBA) | 1.09±0.54 | 0.46±0.16 | 0.88±0.31 | 0.86±0.43 | 0.32 | 0.31 |
| Maximum branch (MSBR) | 3.87±0.52 | 2.00±0.00 | 3.43±1.02 | 2.97±0.87 | 0.23 | 0.47 |
| Stolon leaf length (SLL) | 53.37±9.67 | 42.48±10.51 | 46.79±12.96 | 43.71±16.87 | 0.60 | 0.63 |
| Stolon leaf width (SLW) | 23.99±4.16 | 30.09±9.84 | 24.27±4.72 | 24.96±10.41 | 0.05 | 0.63 |
| **Rhizome traits** | | | | | | |
| Rhizome number 1 (RH1) | 1.65±1.41 | 0 | 0.19±0.40 | 0.07±0.32 | 0.88 | 0 |
| Rhizome number 2 (RH2) | 5.96±1.65 | 0 | 0.5±0.51 | 0.55±0.99 | 0.92 | 0.02 |
| Rhizome number 3 (RH3) | 8.43±1.80 | 0 | 1.31±1.14 | 0.94±1.36 | 0.84 | 0.22 |
| Emergence node 1 (REM1) | 1.11±0.32 | NA | 1±0 | 1.03±0.18 | NA | 0 |
| Emergence node 2 (REM2) | 2.63±0.65 | NA | 1±0 | 1.64±0.78 | NA | 0.78 |
| Emergence node 3 (REM3) | 3.50±0.76 | NA | 1.69±0.85 | 1.93±1.02 | NA | NA |
| Rhizome length (RL) | 15.50±3.02 | NA | 33.88±19.22 | 21.04±15.35 | NA | NA |
| Rhizome width (RW) | 1.26±0.18 | NA | 1.79±0.51 | 2.03±1.73 | NA | NA |
| Rhizome node (RN) | 7.90±1.45 | NA | 7.50±1.60 | 5.60±2.58 | NA | NA |
| Rhizome branch (RB) | 13.33±4.60 | NA | 7.78±4.92 | 6.43±5.33 | NA | NA |
| Rhizome branch/node (RBA) | 1.66±0.40 | NA | 0.93±0.62 | 1.00±0.64 | NA | NA |
| Maximum branch (MRBR) | 2.67±0.91 | NA | 1.56±0.88 | 1.67±0.92 | NA | NA |
| Rhizome leaf length (RLL) | 4.19±3.49 | NA | 6.28±6.16 | 10.81±10.51 | NA | NA |
| Rhizome leaf width (RLW) | 2.65±1.97 | NA | 4.59±5.21 | 6.50±5.44 | NA | NA |
| Rhizome proportion 1 (PR1) | 0.33±0.22 | 0 | 0.03±0.06 | 0.02±0.07 | 0.90 | 0 |
| Rhizome proportion 2 (PR2) | 0.45±0.10 | 0 | 0.05±0.06 | 0.05±0.09 | 0.88 | 0.44 |
| Rhizome proportion 3 (PR3) | 0.58±0.08 | 0 | 0.10±0.08 | 0.08±0.10 | 0.83 | 0.63 |

**Table S2. Fraser's *v*-test results for detecting selection on measured traits. *p*-values are based on F-tests comparing observed trait divergence to the neutral expectation.** Holm's sequential Bonferroni correction was applied to account for multiple testing. Asterisks indicate significance levels: * $p < 0.05$, ** $p < 0.01$, *** $p < 0.001$.

| Trait | *v* (LVR) | *v* (CWF) | *v* (EAM) | *v* (IMPO) |
|---|---|---|---|---|
| **Floral traits** | | | | |
| Corolla tube length (CTL) | 5.06 | 9.74* | 85.69*** | 1.21 |
| Corolla limb length (CLL) | 0.01 | 1.55 | 1.59 | 0.67 |
| Corolla limb width (CLW) | 0.10 | 0.35 | NA | 0.33 |
| **Phenological traits** | | | | |
| Flower node (FN) | 0.64 | 0.14 | NA | 0.13 |
| Flower time (FT) | 1.88 | 3.22 | NA | 1.78 |
| Internode length (IL) | 0.36 | 2.00 | NA | NA |
| **Size traits** | | | | |
| Leaf length (LL) | 0.97 | 1.04 | 0.25 | 0.03 |
| Leaf width (LW) | 6.41 | 0.69 | 0.10 | 1.21 |
| Internode width (IW) | 9.31 | 5.85 | 13.42** | 2.70 |
| **Stolon traits** | | | | |
| Stolon number 1 (ST1) | 5.39 | 2.64 | NA | 0.76 |
| Stolon number 2 (ST2) | 0.11 | 2.82 | NA | 0.40 |
| Stolon number 3 (ST3) | 0.02 | 4.31 | NA | 0.00 |
| Emergence node 1 (SEM1) | 2.14 | 2.05 | 67.19*** | 2.67 |
| Emergence node 2 (SEM2) | 0.08 | NA | NA | 1.30 |
| Emergence node 3 (SEM3) | 0.93 | 2.06 | NA | 1.01 |
| Stolon length (SL) | 9.09 | 1.11 | 2.42 | 2.05 |
| Stolon width (SW) | 31.33*** | 24.14*** | NA | 10.88* |
| Stolon node (SN) | 0.05 | NA | 0.99 | 0.11 |
| Stolon branch (SB) | 0.36 | 9.58* | NA | 6.98 |
| Stolon branch/node (SBA) | 0.03 | 4.36 | NA | 3.31 |
| Maximum branch (MSBR) | NA | 38.92*** | NA | 4.90 |
| Stolon leaf length (SLL) | NA | 2.52 | NA | 0.31 |
| Stolon leaf width (SLW) | NA | 1.81 | NA | 0.26 |
| **Rhizome traits** | | | | |
| Rhizome number 1 (RH1) | NA | 12.00* | NA | NA |
| Rhizome number 2 (RH2) | 2.67 | 4.84 | NA | NA |
| Rhizome number 3 (RH3) | 7.35 | NA | NA | 86.68*** |
| Emergence node 1 (REM1) | NA | NA | NA | NA |
| Emergence node 2 (REM2) | NA | NA | NA | NA |
| Emergence node 3 (REM3) | NA | NA | NA | NA |
| Rhizome length (RL) | NA | NA | NA | NA |
| Rhizome width (RW) | NA | NA | NA | NA |
| Rhizome node (RN) | NA | NA | NA | NA |
| Rhizome branch (RB) | NA | NA | NA | NA |
| Rhizome branch/node (RBA) | NA | NA | NA | NA |
| Maximum branch (MRBR) | NA | NA | NA | NA |
| Rhizome leaf length (RLL) | NA | NA | NA | NA |
| Rhizome leaf width (RLW) | NA | NA | NA | NA |
| Rhizome proportion 1 (PR1) | NA | 0.81 | 8.73** | NA |
| Rhizome proportion 2 (PR2) | 2.78 | 0.77 | NA | 30.11*** |
| Rhizome proportion 3 (PR3) | 3.63 | 8.62 | 13.30** | 25.04*** |

**Table S3. Average correlation coefficients (absolute values; mean ± SE) within and between trait categories across the four mapping populations.** A) The coastal *M. guttatus* × *M. tilingii* population. B) The coastal *M. guttatus* × CWF population. C) The coastal *M. guttatus* × *M. corallinus* population. D) The coastal *M. guttatus* × *M. decorus* population. Significance levels: ∗ p < 0.05, ∗∗ p < 0.01, ∗∗∗< 0.001, determined by permutation test with 1000 permutations.

A

| Pair of trait category | Average correlation coefficients (mean ± SE) | *p* values |
|---|---|---|
| Floral – Floral | 0.53 ± 0.04 | 0.002** |
| Floral – Phenology | 0.06 ± 0.02 | |
| Floral – Size | 0.28 ± 0.03 | |
| Floral – Stolon | 0.09 ± 0.01 | |
| Floral – Rhizome | 0.08 ± 0.01 | |
| Phenology – Phenology | 0.59 ± 0.12 | 0.009** |
| Phenology – Size | 0.44 ± 0.04 | |
| Phenology – Stolon | 0.27 ± 0.03 | |
| Phenology – Rhizome | 0.07 ± 0.01 | |
| Size – Size | 0.33 ± 0.26 | 0.187 |
| Size – Stolon | 0.24± 0.02 | |
| Size – Rhizome | 0.11 ± 0.01 | |
| Stolon – Stolon | 0.25 ± 0.03 | 0.012* |
| Stolon – Rhizome | 0.09± 0.01 | |
| Rhizome – Rhizome | 0.30 ± 0.03 | 0*** |

B

| Pair of trait category | Average correlation coefficients (mean ± SE) | *p* values |
|---|---|---|
| Floral – Floral | 0.51 ± 0.04 | 0.016* |
| Floral – Phenology | 0.12 ± 0.03 | |
| Floral – Size | 0.31 ± 0.04 | |
| Floral – Stolon | 0.18 ± 0.02 | |
| Floral – Rhizome | 0.10 ± 0.01 | |
| Phenology – Phenology | 0.70 ± 0.02 | 0.003** |

| | | |
|---|---|---|
| Phenology – Size | 0.41 ± 0.08 | |
| Phenology – Stolon | 0.21 ± 0.03 | |
| Phenology – Rhizome | 0.11 ± 0.01 | |
| Size – Size | 0.49 ± 0.22 | 0.037* |
| Size – Stolon | 0.20 ± 0.03 | |
| Size – Rhizome | 0.06 ± 0.01 | |
| Stolon – Stolon | 0.26 ± 0.03 | 0.01* |
| Stolon – Rhizome | 0.09 ± 0.01 | |
| Rhizome – Rhizome | 0.36 ± 0.04 | 0*** |

C

| **Pair of trait category** | **Average correlation coefficients (mean ± SE)** | ***p* values** |
|---|---|---|
| Floral – Floral | 0.38 ± 0.01 | 0.012* |
| Floral – Phenology | 0.06 ± 0.01 | |
| Floral – Size | 0.10 ± 0.01 | |
| Floral – Stolon | 0.05 ± 0.01 | |
| Floral – Rhizome | 0.05 ± 0.01 | |
| Phenology – Phenology | 0.62 ± 0.06 | 0*** |
| Phenology – Size | 0.31 ± 0.07 | |
| Phenology – Stolon | 0.17 ± 0.02 | |
| Phenology – Rhizome | 0.07 ± 0.01 | |
| Size – Size | 0.64 ± 0.13 | 0.001** |
| Size – Stolon | 0.14 ± 0.02 | |
| Size – Rhizome | 0.09 ± 0.01 | |
| Stolon – Stolon | 0.22 ± 0.03 | 0.003** |
| Stolon – Rhizome | 0.09 ± 0.01 | |
| Rhizome – Rhizome | 0.23 ± 0.03 | 0*** |

D

| **Pair of trait category** | **Average correlation coefficients (mean ± SE)** | ***p* values** |
|---|---|---|
| Floral – Floral | 0.54 ± 0.06 | 0.005** |
| Floral – Phenology | 0.14 ± 0.03 | |
| Floral – Size | 0.19 ± 0.03 | |
| Floral – Stolon | 0.14 ± 0.02 | |
| Floral – Rhizome | 0.07 ± 0.01 | |
| Phenology – Phenology | 0.54 ± 0.09 | 0.013* |
| Phenology – Size | 0.31 ± 0.06 | |
| Phenology – Stolon | 0.21 ± 0.03 | |
| Phenology – Rhizome | 0.10 ± 0.01 | |
| Size – Size | 0.61 ± 0.11 | 0.004** |
| Size – Stolon | 0.19 ± 0.02 | |
| Size – Rhizome | 0.16 ± 0.02 | |
| Stolon – Stolon | 0.25 ± 0.03 | 0.013* |
| Stolon – Rhizome | 0.11 ± 0.01 | |
| Rhizome – Rhizome | 0.47 ± 0.03 | 0*** |

**Table S4. Average correlation coefficients (absolute values; mean ± SE) between trait categories of stolon-related traits (the number of stolons and the highest node of stolon emergence) across the four mapping populations, consistently measured at three developmental stages.** Trait categories are defined based on developmental stages: vegetative, early reproductive, and late reproductive. A) The coastal *M. guttatus* × *M. tilingii* population. B) The coastal *M. guttatus* × CWF population. C) The coastal *M. guttatus* × *M. corallinus* population. D) The coastal *M. guttatus* × *M. decorus* population. Significance levels were determined by a permutation test with 1000 permutations.

A

| **Trait category pair for stolon traits** | **Average correlation coefficients (mean ± SE)** | ***p* values** |
|---|---|---|
| vegetative – vegetative | 0.78 ± NA | 0.073 |
| vegetative – early reproductive | 0.25 ± 0.05 | |
| vegetative – late reproductive | 0.12 ± 0.03 | |
| early reproductive – early reproductive | 0.71 ± NA | 0.305 |
| early reproductive – late reproductive | 0.65 ± 0.06 | |
| late reproductive – late reproductive | 0.70 ± NA | 0.263 |

B

| **Trait category pair for stolon traits** | **Average correlation coefficients (mean ± SE)** | ***p* values** |
|---|---|---|
| vegetative – vegetative | 0.77 ± NA | 0.043 |
| vegetative – early reproductive | 0.11 ± 0.06 | |
| vegetative – late reproductive | 0.09 ± 0.02 | |
| early reproductive – early reproductive | 0.73 ± NA | 0.227 |
| early reproductive – late reproductive | 0.59 ± 0.06 | |
| late reproductive – late reproductive | 0.66 ± NA | 0.275 |

C

| **Trait category pair for stolon traits** | **Average correlation coefficients (mean ± SE)** | ***p* values** |
|---|---|---|
| vegetative – vegetative | 0.71 ± NA | 0.06 |
| vegetative – early reproductive | 0.09 ± 0.03 | |
| vegetative – late reproductive | 0.05 ± 0.02 | |
| early reproductive – early reproductive | 0.81 ± NA | 0.125 |

| | | |
|---|---|---|
| early reproductive – late reproductive | 0.39 ± 0.04 | |
| late reproductive – late reproductive | 0.70 ± NA | 0.198 |

D

| **Trait category pair for stolon traits** | **Average correlation coefficients (mean ± SE)** | ***p* values** |
|---|---|---|
| vegetative – vegetative | 0.71 ± NA | 0.05 |
| vegetative – early reproductive | 0.09 ± 0.03 | |
| vegetative – late reproductive | 0.07 ± 0.01 | |
| early reproductive – early reproductive | 0.72 ± NA | 0.214 |
| early reproductive – late reproductive | 0.60 ± 0.06 | |
| late reproductive – late reproductive | 0.75 ± NA | 0.141 |

**Table S5. Average correlation coefficients (absolute values; mean ± SE) between trait categories of rhizome-related traits (the number of rhizomes and the highest node of rhizome emergence) across the four mapping populations, consistently measured at three developmental stages.** Trait categories are defined based on developmental stages: vegetative, early reproductive, and late reproductive. A) The coastal *M. guttatus* × *M. tilingii* population. B) The coastal *M. guttatus* × CWF population. C) The coastal *M. guttatus* × *M. corallinus* population. D) The coastal *M. guttatus* × *M. decorus* population. Significance levels: ∗ $p < 0.05$, determined by permutation test with 1000 permutations.

A

| Trait category pair for rhizome traits | Average correlation coefficients (mean ± SE) | *p* values |
|---|---|---|
| vegetative – vegetative | NA ± NA | NA |
| vegetative – early reproductive | NA ± NA | |
| vegetative – late reproductive | NA ± NA | |
| early reproductive – early reproductive | 0.29 ± NA | 0.64 |
| early reproductive – late reproductive | 0.47 ± 0.17 | |
| late reproductive – late reproductive | 0.33 ± NA | 0.47 |

B

| Trait category pair for rhizome traits | Average correlation coefficients (mean ± SE) | *p* values |
|---|---|---|
| vegetative – vegetative | 0.11 ± NA | 0.42 |
| vegetative – early reproductive | 0.23 ± 0.11 | |
| vegetative – late reproductive | 0.19 ± 0.12 | |
| early reproductive – early reproductive | 0.21 ± NA | 0.34 |
| early reproductive – late reproductive | 0.31 ± 0.15 | |
| late reproductive – late reproductive | 0.39 ± NA | 0.19 |

C

| Trait category pair for rhizome traits | Average correlation coefficients (mean ± SE) | *p* values |
|---|---|---|
| vegetative – vegetative | 0.33 ± NA | 0.32 |
| vegetative – early reproductive | 0.25 ± 0.06 | |
| vegetative – late reproductive | 0.16 ± 0.06 | |
| early reproductive – early reproductive | 0.72 ± NA | 0.12 |
| early reproductive – late reproductive | 0.58 ± 0.05 | |

| | | |
|---|---|---|
| late reproductive – late reproductive | 0.69 ± NA | 0.07 |

D

| **Trait category pair for rhizome traits** | **Average correlation coefficients (mean ± SE)** | ***p* values** |
|---|---|---|
| vegetative – vegetative | 0.29 ± NA | 0.31 |
| vegetative – early reproductive | 0.20 ± 0.05 | |
| vegetative – late reproductive | 0.14 ± 0.04 | |
| early reproductive – early reproductive | 0.56 ± NA | 0.25 |
| early reproductive – late reproductive | 0.57 ± 0.12 | |
| late reproductive – late reproductive | 0.67 ± NA | 0.21 |

**Table S6. Quantitative Trait Loci (QTLs) associated with life-history trait divergence in the four mapping populations.** (A) QTLs identified in the LVR × OPB population. (B) QTLs identified in the CWF × OPB population.

(C) QTLs identified in the EAM × OPB population. (D) QTLs identified in the IMPO × OPB population. QTLs are organized by trait category as in Fig. 3: (1) Floral traits, (2) Phenological traits, (3) Size traits, (4) Stolon traits, and (5) Rhizome traits. For each QTL, the additive effect (*a*), dominance effect (*d*), the proportion of phenotypic variance explained in the $F_2$ population ($r^2$), peak logarithm of odds (LOD) score, and linkage group (LG) are reported. The QTL position is given in centiMorgans (cM) based on the linkage map, including the peak position and its 1.5 LOD-drop confidence interval (cMmin, cMpeak, cMmax), along with the corresponding chromosome number (Chrom) and genomic coordinates of the peak and confidence interval (bmin, bpeak, bmax; in units of 100,000 bp).

**A:** LVR × OPB

### 1. Floral traits

| Trait | QTL | *a* | *d* | $r^2$ (%) | LOD | LG | cMmin | cMpeak | cMmax | Chrom | bmin | bpeak | bmax |
|---|---|---|---|---|---|---|---|---|---|---|---|---|---|
| CTL | CTL2 | 0.67 | -0.09 | 2.22 | 3.57 | 2 | 0.00 | 10.26 | 45.57 | 2 | 0 | 13 | 21 |
| CTL | CTL5 | 0.36 | 0.63 | 2.29 | 3.64 | 5 | 36.19 | 49.00 | 58.30 | 5 | 39 | 57 | 217 |
| CTL | CTL6 | -0.59 | 0.49 | 2.54 | 4.87 | 6 | 32.01 | 79.55 | 85.27 | 6 | 39 | 188 | 198 |
| CTL | CTL8 | -1.53 | 0.51 | 15.11 | 22.09 | 8 | 10.57 | 11.98 | 16.06 | 8 | 7 | 14 | 159 |
| CTL | CTL10 | 0.70 | -0.44 | 3.35 | 3.58 | 10 | 0.00 | 32.01 | 72.83 | 10 | 0 | 13 | 60 |
| CTL | CTL13 | -1.41 | 0.02 | 13.36 | 18.75 | 13 | 43.51 | 44.27 | 46.43 | 13 | 251 | 253 | 301 |
| CLL | CLL5 | 2.07 | 0.89 | 8.43 | 17.65 | 5 | 36.19 | 50.68 | 55.87 | 5 | 39 | 65 | 160 |
| CLL | CLL7 | 0.24 | 0.20 | 0.08 | 4.85 | 7 | 74.04 | 83.97 | 93.12 | 7 | 132 | 136 | 142 |
| CLL | CLL8 | 1.08 | 0.72 | 3.68 | 4.70 | 8 | 56.28 | 80.15 | 91.13 | 8 | 239 | 252 | 276 |
| CLL | CLL10 | 0.83 | -0.92 | 1.89 | 3.53 | 10 | 0.00 | 70.80 | 97.25 | 10 | 0 | 54 | 86 |
| CLL | CLL11 | 0.87 | -0.38 | 1.95 | 3.72 | 11 | 33.40 | 55.23 | 69.75 | 11 | 85 | 280 | 334 |
| CLL | CLL12 | 1.04 | 0.17 | 2.58 | 6.27 | 12 | 25.40 | 35.31 | 61.43 | 12 | 184 | 212 | 250 |
| CLL | CLL13 | -1.0 | 0.53 | 2.58 | 4.27 | 13 | 0.00 | 0.00 | 55.02 | 13 | 0 | 0 | 306 |
| CLL | CLL14 | -1.15 | 1.00 | 2.84 | 5.48 | 14 | 101.05 | 107.58 | 137.72 | 14 | 168 | 192 | 260 |
| CLW | CLW1 | -0.94 | -0.28 | 2.88 | 3.92 | 1 | 34.39 | 38.80 | 61.66 | 1 | 39 | 45 | 119 |
| CLW | CLW5 | 2.08 | 0.74 | 13.44 | 18.15 | 5 | 38.70 | 41.00 | 49.88 | 5 | 46 | 52 | 57 |
| CLW | CLW11 | -0.79 | 0.85 | 2.77 | 3.72 | 11 | 2.08 | 9.64 | 24.62 | 11 | 3 | 10 | 29 |
| CLW | CLW13 | -0.96 | -0.09 | 2.71 | 4.27 | 13 | 0.00 | 2.00 | 12.75 | 13 | 0 | 2 | 10 |
| CLW | CLW14 | -1.09 | 0.84 | 3.73 | 3.53 | 14 | 9.47 | 103.65 | 120.52 | 14 | 11 | 180 | 227 |

### 2. Phenological traits

| Trait | QTL | *a* | *d* | $r^2$ (%) | LOD | LG | cMmin | cMpeak | cMmax | Chrom | bmin | bpeak | bmax |
|---|---|---|---|---|---|---|---|---|---|---|---|---|---|
| FN | FN5 | -0.49 | -0.36 | 4.42 | 6.94 | 5 | 52.28 | 57.76 | 58.30 | 5 | 78 | 177 | 217 |
| FN | FN6 | -0.52 | -0.29 | 6.94 | 10.26 | 6 | 79.55 | 81.72 | 88.00 | 6 | 188 | 193 | 202 |
| FN | FN7 | 0.32 | -0.26 | 3.55 | 4.97 | 7 | 24.36 | 83.97 | 103.63 | 7 | 25 | 136 | 179 |
| FN | FN8 | 0.69 | -0.51 | 11.82 | 14.29 | 8 | 87.42 | 90.59 | 91.13 | 8 | 271 | 274 | 276 |
| FN | FN12 | 0.31 | -0.42 | 3.25 | 4.09 | 12 | 2.95 | 21.47 | 31.10 | 12 | 7 | 170 | 201 |
| FT | FT4 | -1.80 | -2.26 | 4.51 | 5.44 | 4 | 8.03 | 19.59 | 22.76 | 4 | 15 | 32 | 37 |
| FT | FT5 | -3.92 | -3.87 | 15.37 | 25.93 | 5 | 55.08 | 57.46 | 58.21 | 5 | 127 | 172 | 200 |
| FT | FT6 | -1.45 | -2.10 | 4.37 | 7.36 | 6 | 49.82 | 83.71 | 89.63 | 6 | 142 | 195 | 204 |
| FT | FT7 | 1.39 | -1.15 | 2.74 | 9.48 | 7 | 74.04 | 83.00 | 93.12 | 7 | 132 | 136 | 142 |
| FT | FT8 | 1.77 | -2.01 | 4.17 | 5.89 | 8 | 83.27 | 87.42 | 91.13 | 8 | 263 | 271 | 276 |
| FT | FT14 | 1.01 | -2.13 | 2.09 | 3.75 | 14 | 28.97 | 91.00 | 107.58 | 14 | 59 | 118 | 192 |
| IL | IL4 | 3.83 | -0.34 | 8.76 | 12.96 | 4 | 40.33 | 45.00 | 51.31 | 4 | 80 | 156 | 178 |
| IL | IL6 | 0.96 | 1.46 | 1.65 | 4.10 | 6 | 81.15 | 88.00 | 91.88 | 6 | 192 | 202 | 208 |
| IL | IL7 | -2.88 | -0.58 | 5.03 | 5.91 | 7 | 26.18 | 48.00 | 63.10 | 7 | 29 | 34 | 54 |
| IL | IL8 | -4.26 | 0.46 | 12.57 | 15.31 | 8 | 79.48 | 83.76 | 87.02 | 8 | 248 | 264 | 269 |
| IL | IL14 | -1.41 | -1.38 | 2.17 | 4.32 | 14 | 28.97 | 94.00 | 127.83 | 14 | 59 | 118 | 238 |

### 3. Size traits

| Trait | QTL | *a* | *d* | $r^2$ (%) | LOD | LG | cMmin | cMpeak | cMmax | Chrom | bmin | bpeak | bmax |
|---|---|---|---|---|---|---|---|---|---|---|---|---|---|
| LL | LL4 | -0.19 | -0.12 | 1.54 | 4.54 | 4 | 21.63 | 32.98 | 51.31 | 4 | 36 | 55 | 178 |

| | | | | | | | | | | | | | |
|---|---|---|---|---|---|---|---|---|---|---|---|---|---|
| LL | LL5 | -0.09 | 0.09 | 0.45 | 4.15 | 5 | 21.28 | 25.03 | 35.79 | 5 | 22 | 27 | 38 |
| LL | LL6 | -0.30 | 0.01 | 3.67 | 3.59 | 6 | 62.71 | 82.22 | 91.88 | 6 | 175 | 194 | 208 |
| LL | LL8 | -0.31 | -0.04 | 4.39 | 11.40 | 8 | 0.00 | 16.43 | 26.46 | 8 | 0 | 161 | 179 |
| LL | LL11 | 0.09 | -0.17 | 1.04 | 4.74 | 11 | 19.70 | 45.84 | 60.21 | 11 | 23 | 236 | 321 |
| LL | LL13 | -0.04 | -0.15 | 0.57 | 5.09 | 13 | 33.06 | 44.36 | 49.59 | 13 | 240 | 264 | 303 |
| LL | LL14 | 0.15 | -0.16 | 1.02 | 6.24 | 14 | 107.58 | 137.72 | 138.83 | 14 | 192 | 260 | 262 |
| LW | LW6 | -1.85 | 1.11 | 3.41 | 4.67 | 6 | 3.57 | 17.21 | 30.63 | 6 | 7 | 20 | 37 |
| LW | LW7 | -1.41 | -0.49 | 1.37 | 4.00 | 7 | 0.00 | 55.00 | 60.76 | 7 | 1 | 37 | 49 |
| LW | LW8 | -2.24 | 0.81 | 3.82 | 5.93 | 8 | 0.00 | 0.00 | 61.43 | 8 | 0 | 0 | 242 |
| LW | LW9 | -1.10 | 0.08 | 0.90 | 3.53 | 9 | 6.18 | 16.59 | 55.30 | 9 | 5 | 21 | 215 |
| LW | LW11 | 2.07 | -0.09 | 3.34 | 4.51 | 11 | 13.28 | 46.00 | 60.21 | 11 | 14 | 238 | 321 |
| LW | LW12 | -1.84 | 1.11 | 3.18 | 3.47 | 12 | 0.00 | 13.93 | 61.43 | 12 | 1 | 29 | 250 |
| LW | LW13 | -3.36 | -0.48 | 8.78 | 9.11 | 13 | 41.24 | 44.97 | 46.43 | 13 | 249 | 300 | 301 |
| LW | LW14 | -2.28 | 1.32 | 3.72 | 5.16 | 14 | 97.61 | 103.74 | 116.19 | 14 | 159 | 181 | 215 |
| IW | IW2 | -0.28 | -0.03 | 2.86 | 3.90 | 2 | 55.80 | 62.05 | 68.57 | 2 | 51 | 141 | 162 |
| IW | IW4 | -0.22 | -0.23 | 2.48 | 6.00 | 4 | 40.33 | 47.35 | 51.31 | 4 | 80 | 163 | 178 |
| IW | IW6 | -0.28 | -0.05 | 3.45 | 7.41 | 6 | 81.15 | 85.27 | 89.63 | 6 | 192 | 198 | 204 |
| IW | IW7 | 0.35 | -0.17 | 6.29 | 8.26 | 7 | 27.00 | 39.00 | 63.10 | 7 | 30 | 32 | 54 |
| IW | IW8 | -0.34 | 0.03 | 4.80 | 7.61 | 8 | 8.33 | 11.98 | 91.13 | 8 | 6 | 72 | 276 |
| IW | IW12 | -0.30 | -0.06 | 3.92 | 4.67 | 12 | 58.66 | 63.12 | 66.94 | 12 | 247 | 252 | 258 |
| IW | IW13 | -0.33 | -0.08 | 4.56 | 8.17 | 13 | 0.00 | 0.00 | 4.92 | 13 | 0 | 0 | 4 |

**4. Stolon traits**

| Trait | QTL | *a* | *d* | $r^2$ (%) | LOD | LG | cMmin | cMpeak | cMmax | Chrom | bmin | bpeak | bmax |
|---|---|---|---|---|---|---|---|---|---|---|---|---|---|
| ST1 | ST1_5 | 0.84 | 0.73 | 12.71 | 23.48 | 5 | 55.08 | 56.23 | 58.21 | 5 | 127 | 161 | 200 |
| ST1 | ST1_6 | 0.37 | 0.10 | 2.30 | 4.12 | 6 | 26.53 | 44.20 | 76.60 | 6 | 34 | 81 | 177 |
| ST1 | ST1_7 | 0.13 | 0.23 | 0.29 | 8.10 | 7 | 74.04 | 83.00 | 93.12 | 7 | 132 | 136 | 142 |
| ST1 | ST1_13 | 0.39 | 0.45 | 3.98 | 4.17 | 13 | 44.36 | 48.00 | 55.02 | 13 | 299 | 302 | 306 |
| ST1 | ST1_14 | 0.62 | 0.11 | 6.59 | 7.59 | 14 | 122.98 | 138.36 | 141.42 | 14 | 230 | 261 | 265 |
| ST2 | ST2_7 | 0.68 | -0.06 | 3.93 | 8.36 | 7 | 17.67 | 37.00 | 59.56 | 7 | 19 | 32 | 46 |
| ST2 | ST2_8 | 0.72 | -0.34 | 5.18 | 5.43 | 8 | 86.14 | 91.00 | 91.13 | 8 | 268 | 275 | 276 |
| ST2 | ST2_13 | 0.63 | 0.59 | 5.30 | 5.41 | 13 | 43.61 | 46.43 | 55.02 | 13 | 252 | 301 | 306 |
| ST2 | ST2_14 | 0.91 | 0.01 | 6.67 | 10.29 | 14 | 122.98 | 125.76 | 135.28 | 14 | 230 | 233 | 253 |
| ST3 | ST3_5 | -0.76 | -0.15 | 4.43 | 4.93 | 5 | 8.76 | 21.00 | 58.30 | 5 | 9 | 21 | 217 |
| ST3 | ST3_7 | 0.48 | -0.31 | 2.48 | 4.87 | 7 | 26.18 | 59.56 | 64.59 | 7 | 29 | 46 | 65 |
| ST3 | ST3_8 | -0.91 | 0.02 | 5.72 | 4.77 | 8 | 3.33 | 10.57 | 91.13 | 8 | 3 | 7 | 276 |
| ST3 | ST3_12 | -0.64 | 0.02 | 3.07 | 3.91 | 12 | 16.58 | 48.00 | 54.81 | 12 | 33 | 235 | 242 |
| ST3 | ST3_13 | 0.59 | 0.47 | 3.45 | 3.64 | 13 | 41.24 | 47.28 | 55.02 | 13 | 249 | 302 | 307 |
| ST3 | ST3_14 | 0.71 | 0.02 | 3.42 | 5.73 | 14 | 99.32 | 125.76 | 135.28 | 14 | 163 | 233 | 253 |
| SEM1 | SEM1_4 | -0.19 | 0.07 | 5.87 | 8.08 | 4 | 42.55 | 46.06 | 49.12 | 4 | 138 | 161 | 169 |
| SEM1 | SEM1_5 | 0.22 | 0.18 | 6.79 | 13.46 | 5 | 55.30 | 57.17 | 58.21 | 5 | 135 | 170 | 200 |
| SEM1 | SEM1_7 | 0.07 | 0.09 | 0.47 | 5.92 | 7 | 74.04 | 83.00 | 93.12 | 7 | 132 | 136 | 142 |
| SEM1 | SEM1_14 | 0.15 | 0.11 | 5.47 | 5.68 | 14 | 122.98 | 137.60 | 141.42 | 14 | 230 | 259 | 265 |
| SEM2 | SEM2_4 | -0.31 | -0.19 | 5.51 | 5.51 | 4 | 28.50 | 41.52 | 51.31 | 4 | 46 | 89 | 178 |
| SEM2 | SEM2_5 | -0.23 | -0.21 | 4.58 | 4.58 | 5 | 20.15 | 50.28 | 58.30 | 5 | 20 | 58 | 217 |
| SEM2 | SEM2_7 | 0.29 | -0.01 | 5.71 | 5.71 | 7 | 6.84 | 37.00 | 63.10 | 7 | 9 | 32 | 54 |
| SEM2 | SEM2_8 | 0.40 | -0.12 | 10.96 | 10.96 | 8 | 87.42 | 90.59 | 91.13 | 8 | 271 | 274 | 276 |
| SEM2 | SEM2_12 | 0.17 | -0.24 | 3.83 | 3.83 | 12 | 13.93 | 22.41 | 66.94 | 12 | 29 | 175 | 258 |
| SEM3 | SEM3_4 | -0.26 | -0.23 | 4.50 | 5.24 | 4 | 28.50 | 41.52 | 46.98 | 4 | 46 | 89 | 162 |
| SEM3 | SEM3_5 | -0.28 | -0.28 | 7.23 | 7.88 | 5 | 49.88 | 57.76 | 58.30 | 5 | 57 | 175 | 217 |
| SEM3 | SEM3_7 | 0.23 | -0.12 | 4.10 | 4.65 | 7 | 10.99 | 61.40 | 93.12 | 7 | 13 | 52 | 142 |
| SEM3 | SEM3_8 | 0.45 | -0.16 | 12.80 | 13.14 | 8 | 88.57 | 91.00 | 91.13 | 8 | 272 | 275 | 276 |
| SEM3 | SEM3_12 | -0.22 | -0.00 | 2.76 | 4.01 | 12 | 46.27 | 58.66 | 66.94 | 12 | 233 | 247 | 258 |
| SL | SL8 | -5.08 | 2.18 | 8.61 | 6.84 | 8 | 8.33 | 19.86 | 26.46 | 8 | 6 | 174 | 179 |
| SL | SL9 | -3.42 | -0.81 | 4.92 | 4.08 | 9 | 14.31 | 19.00 | 109.78 | 9 | 16 | 28 | 250 |

| | | | | | | | | | | | | | |
|---|---|---|---|---|---|---|---|---|---|---|---|---|---|
| SW | SW6 | -0.20 | -0.07 | 6.27 | 4.13 | 6 | 3.57 | 10.69 | 18.31 | 6 | 7 | 15 | 22 |
| SW | SW14 | -0.15 | -0.17 | 6.27 | 4.13 | 14 | 115.37 | 136.50 | 138.36 | 14 | 212 | 256 | 261 |
| SN | SN2 | -1.25 | 1.95 | 4.94 | 3.49 | 2 | 85.14 | 87.45 | 89.48 | 2 | 199 | 201 | 205 |
| SN | SN4 | -1.28 | 0.40 | 5.53 | 3.99 | 4 | 36.99 | 71.00 | 80.80 | 4 | 68 | 210 | 228 |
| SN | SN5 | 1.01 | 0.75 | 4.84 | 4.33 | 5 | 45.40 | 55.30 | 58.30 | 5 | 55 | 135 | 217 |
| SB | SB6 | -0.49 | -0.63 | 3.88 | 3.51 | 6 | 48.02 | 83.00 | 91.88 | 6 | 137 | 195 | 208 |
| SB | SB8 | 1.59 | -0.47 | 19.60 | 15.34 | 8 | 89.94 | 91.13 | 91.13 | 8 | 273 | 276 | 276 |
| SBA | SBA8 | 0.15 | -0.07 | 17.95 | 12.9 | 8 | 89.94 | 91.13 | 91.13 | 8 | 273 | 276 | 276 |
| MSBR | MSBR14 | 0.29 | 0.06 | 7.98 | 5.48 | 14 | 128.01 | 130.25 | 141.42 | 14 | 239 | 244 | 265 |
| SLL | SLL6 | 11.03 | -5.64 | 17.31 | 3.92 | 6 | 51.58 | 80.46 | 91.88 | 6 | 163 | 190 | 208 |

**5. Rhizome traits**

| Trait | QTL | *a* | *d* | $r^2$ (%) | LOD | LG | cMmin | cMpeak | cMmax | Chrom | bmin | bpeak | bmax |
|---|---|---|---|---|---|---|---|---|---|---|---|---|---|
| RH1 | RH1_9 | 0.10 | -0.04 | 4.12 | 4.95 | 9 | 6.18 | 15.16 | 57.87 | 9 | 5 | 18 | 221 |
| RH2 | RH2_3 | 0.14 | 0.16 | 2.93 | 3.47 | 3 | 117.50 | 127.87 | 133.75 | 3 | 215 | 225 | 232 |
| RH3 | RH3_3 | 0.20 | 0.14 | 3.46 | 4.32 | 3 | 115.78 | 125.16 | 130.25 | 3 | 212 | 223 | 228 |
| RH3 | RH3_14 | -0.26 | 0.01 | 3.82 | 4.69 | 14 | 0.72 | 8.00 | 16.21 | 14 | 3 | 8 | 23 |
| REM1 | REM1_13 | 0.00 | 0.00 | - | 8.76 | 13 | 44.36 | 44.36 | 44.36 | 13 | 256 | 291 | 297 |
| RL | RL9 | 4.53 | -1.15 | 6.87 | 4.14 | 9 | 0.00 | 2.26 | 35.42 | 9 | 0 | 3 | 41 |
| RL | RL11 | -4.00 | 0.41 | 5.48 | 3.55 | 11 | 22.21 | 32.07 | 60.21 | 11 | 26 | 71 | 321 |
| RN | RN9 | 1.26 | -0.45 | 7.38 | 3.90 | 9 | 0.00 | 3.00 | 35.42 | 9 | 0 | 4 | 41 |
| RN | RN11 | -1.42 | 0.09 | 8.60 | 4.42 | 11 | 23.23 | 54.00 | 61.34 | 11 | 27 | 276 | 322 |
| RB | RB9 | 2.06 | 0.39 | 10.79 | 4.17 | 9 | 12.50 | 16.59 | 60.58 | 9 | 13 | 21 | 226 |
| RBA | RBA8 | 0.29 | -0.06 | 9.16 | 3.76 | 8 | 22.04 | 26.46 | 38.55 | 8 | 177 | 179 | 194 |
| RBA | RBA9 | 0.16 | 0.45 | 17.88 | 7.43 | 9 | 52.30 | 58.00 | 60.96 | 9 | 206 | 221 | 227 |
| MRBR | MRBR9 | 0.25 | 0.53 | 15.18 | 6 | 9 | 49.71 | 52.30 | 56.89 | 9 | 187 | 206 | 219 |
| RLL | RLL2 | - | - | - | 5.82 | 2 | 68.57 | 69.09 | 69.30 | 2 | 162 | 164 | 165 |
| RLW | RLW2 | - | - | - | 5.92 | 2 | 68.57 | 69.09 | 69.30 | 2 | 162 | 164 | 165 |
| PR1 | PR1_9 | 0.03 | -0.01 | 5.08 | 5.38 | 9 | 6.18 | 15.16 | 53.56 | 9 | 5 | 18 | 210 |
| PR2 | PR2_5 | 0.04 | 0.01 | 3.28 | 3.77 | 5 | 31.21 | 50.68 | 58.30 | 5 | 34 | 65 | 217 |

**B:** CWF × OPB

**1. Floral traits**

| Trait | QTL | *a* | *d* | $r^2$ (%) | LOD | LG | cMmin | cMpeak | cMmax | Chrom | bmin | bpeak | bmax |
|---|---|---|---|---|---|---|---|---|---|---|---|---|---|
| CTL | CTL2 | -0.55 | 0.06 | 2.17 | 4.57 | 2 | 88.73 | 102.00 | 124.02 | 2 | 145 | 149 | 156 |
| CTL | CTL3 | -0.81 | -0.19 | 4.96 | 14.59 | 3 | 68.61 | 72.00 | 76.93 | 3 | 218 | 223 | 228 |
| CTL | CTL4 | -0.81 | -0.22 | 5.16 | 5.36 | 4 | 43.48 | 54.00 | 64.28 | 4 | 73 | 171 | 197 |
| CTL | CTL5 | 1.03 | 1.03 | 10.49 | 13.41 | 5 | 36.37 | 43.00 | 65.59 | 5 | 59 | 170 | 188 |
| CTL | CTL7 | -0.57 | -0.04 | 2.91 | 3.55 | 7 | 0.00 | 1.00 | 177.08 | 7 | 0 | 3 | 198 |
| CTL | CTL8 | -0.24 | -0.41 | 1.38 | 4.26 | 8 | 0.84 | 75.51 | 86.81 | 8 | 6 | 208 | 211 |
| CTL | CTL9 | -0.44 | 0.05 | 1.08 | 3.91 | 9 | 7.29 | 62.85 | 72.38 | 9 | 11 | 232 | 236 |
| CTL | CTL12 | -0.31 | 0.40 | 0.86 | 6.77 | 12 | 136.60 | 144.61 | 144.61 | 12 | 249 | 258 | 258 |
| CTL | CTL14 | -0.77 | 0.27 | 3.31 | 5.3 | 14 | 94.75 | 101.71 | 141.04 | 14 | 229 | 238 | 280 |
| CLL | CLL4 | -1.03 | 0.11 | 5.13 | 4.39 | 4 | 33.73 | 37.00 | 54.20 | 4 | 53 | 58 | 171 |
| CLL | CLL5 | 0.97 | 1.04 | 6.61 | 6.94 | 5 | 30.74 | 39.52 | 65.59 | 5 | 48 | 161 | 188 |
| CLL | CLL12 | -1.07 | 0.52 | 3.84 | 3.45 | 12 | 100.14 | 117.43 | 123.45 | 12 | 219 | 232 | 233 |
| CLL | CLL13 | -0.86 | -0.29 | 4.24 | 4.89 | 13 | 33.97 | 53.10 | 77.80 | 13 | 245 | 265 | 279 |
| CLW | CLW3 | -1.18 | -0.20 | 4.43 | 8.82 | 3 | 71.03 | 76.00 | 84.39 | 3 | 222 | 228 | 237 |
| CLW | CLW5 | 2.37 | 2.05 | 18.90 | 24.44 | 5 | 38.71 | 44.00 | 65.59 | 5 | 151 | 170 | 188 |
| CLW | CLW6 | 0.47 | 0.80 | 1.91 | 4.77 | 6 | 45.80 | 80.18 | 91.64 | 6 | 53 | 195 | 207 |
| CLW | CLW9 | -1.10 | -0.29 | 3.38 | 6.43 | 9 | 56.19 | 64.00 | 72.38 | 9 | 231 | 233 | 236 |
| CLW | CLW13 | -0.58 | 0.65 | 1.84 | 4.28 | 13 | 20.87 | 27.48 | 46.24 | 13 | 193 | 238 | 255 |
| CLW | CLW14 | -0.79 | 0.46 | 1.11 | 4.38 | 14 | 94.75 | 136.66 | 141.04 | 14 | 229 | 276 | 280 |

### 2. Phenological traits

| Trait | QTL | *a* | *d* | $r^2$ (%) | LOD | LG | cMmin | cMpeak | cMmax | Chrom | bmin | bpeak | bmax |
|---|---|---|---|---|---|---|---|---|---|---|---|---|---|
| FN | FN1 | -0.36 | -0.02 | 3.14 | 4.17 | 1 | 0.00 | 4.00 | 23.66 | 1 | 0 | 0 | 8 |
| FN | FN4 | -0.45 | 0.14 | 6.72 | 4.49 | 4 | 37.04 | 41.00 | 56.85 | 4 | 58 | 64 | 178 |
| FN | FN6 | 0.86 | -0.26 | 17.20 | 17.98 | 6 | 80.60 | 84.00 | 87.89 | 6 | 197 | 199 | 200 |
| FN | FN14 | 0.27 | -0.36 | 1.95 | 4.27 | 14 | 132.25 | 137.41 | 141.04 | 14 | 274 | 278 | 280 |
| FT | FT2 | -1.77 | -1.41 | 4.62 | 3.73 | 2 | 18.71 | 161.00 | 173.51 | 2 | 21 | 191 | 205 |
| FT | FT5 | -3.61 | -3.17 | 13.12 | 12.39 | 5 | 36.37 | 44.00 | 93.48 | 5 | 59 | 170 | 217 |
| FT | FT6 | 2.58 | -0.43 | 4.55 | 4.94 | 6 | 73.49 | 81.00 | 87.89 | 6 | 192 | 197 | 200 |
| FT | FT14 | 1.58 | -1.75 | 1.83 | 3.73 | 14 | 128.88 | 138.00 | 141.04 | 14 | 273 | 279 | 281 |
| IL | IL5 | 2.19 | 1.99 | 4.20 | 4.22 | 5 | 36.37 | 93.48 | 93.48 | 5 | 59 | 217 | 217 |
| IL | IL7 | -0.97 | -2.55 | 3.03 | 3.66 | 7 | 39.81 | 48.01 | 56.33 | 7 | 25 | 32 | 40 |
| IL | IL8 | -2.60 | -0.32 | 4.20 | 4.47 | 8 | 61.27 | 124.00 | 149.65 | 8 | 193 | 241 | 268 |

### 3. Size traits

| Trait | QTL | *a* | *d* | $r^2$ (%) | LOD | LG | cMmin | cMpeak | cMmax | Chrom | bmin | bpeak | bmax |
|---|---|---|---|---|---|---|---|---|---|---|---|---|---|
| LL | LL5 | 3.26 | 4.62 | 10.22 | 7.38 | 5 | 36.37 | 40.59 | 93.48 | 5 | 59 | 170 | 217 |
| LL | LL6 | -5.19 | 0.97 | 12.29 | 9.33 | 6 | 2.06 | 19.78 | 33.47 | 6 | 5 | 28 | 33 |
| LW | LW5 | 4.04 | 5.00 | 14.13 | 11.2 | 5 | 36.37 | 41.00 | 65.59 | 5 | 59 | 170 | 188 |
| LW | LW6 | -4.06 | 0.59 | 8.00 | 5.5 | 6 | 0.00 | 19.78 | 38.91 | 6 | 0 | 28 | 42 |
| IW | IW5 | 0.30 | 0.49 | 9.07 | 10.89 | 5 | 36.37 | 40.59 | 65.59 | 5 | 59 | 170 | 188 |
| IW | IW6 | 0.43 | -0.10 | 8.24 | 9.73 | 6 | 88.21 | 91.64 | 91.64 | 6 | 201 | 207 | 207 |
| IW | IW13 | -0.29 | 0.06 | 4.16 | 5.44 | 13 | 8.24 | 77.80 | 95.79 | 13 | 13 | 279 | 283 |

### 4. Stolon traits

| Trait | QTL | *a* | *d* | $r^2$ (%) | LOD | LG | cMmin | cMpeak | cMmax | Chrom | bmin | bpeak | bmax |
|---|---|---|---|---|---|---|---|---|---|---|---|---|---|
| ST1 | ST1_2 | 0.60 | 0.54 | 4.50 | 3.71 | 2 | 88.24 | 160.92 | 173.51 | 2 | 135 | 191 | 205 |
| ST1 | ST1_5 | 1.40 | 1.28 | 18.61 | 18.36 | 5 | 36.37 | 42.00 | 91.67 | 5 | 59 | 170 | 216 |
| ST2 | ST2_6 | 1.04 | 0.25 | 3.49 | 4.61 | 6 | 73.49 | 80.60 | 87.89 | 6 | 192 | 197 | 200 |
| ST2 | ST2_7 | 0.31 | 1.02 | 2.60 | 3.97 | 7 | 124.18 | 142.10 | 197.46 | 7 | 132 | 159 | 207 |
| ST2 | ST2_10 | 0.76 | 0.80 | 3.45 | 4.20 | 10 | 1.59 | 14.68 | 39.68 | 10 | 5 | 18 | 71 |
| ST2 | ST2_11 | 1.23 | 0.25 | 6.62 | 6.05 | 11 | 29.22 | 33.92 | 44.92 | 11 | 22 | 26 | 44 |
| ST2 | ST2_14 | 1.43 | -0.17 | 5.97 | 6.38 | 14 | 101.71 | 104.87 | 120.27 | 14 | 238 | 241 | 256 |
| ST3 | ST3_6 | 1.12 | -0.25 | 3.06 | 3.82 | 6 | 18.94 | 41.35 | 85.84 | 6 | 27 | 43 | 199 |
| ST3 | ST3_11 | 0.89 | 2.33 | 11.42 | 9.85 | 11 | 32.94 | 42.41 | 44.92 | 11 | 25 | 31 | 44 |
| ST3 | ST3_14 | 1.79 | 0.14 | 7.31 | 5.98 | 14 | 99.53 | 104.35 | 113.86 | 14 | 236 | 240 | 246 |
| SEM1 | SEM1_3 | -0.04 | -0.20 | 4.04 | 3.53 | 3 | 67.30 | 83.48 | 84.39 | 3 | 215 | 234 | 237 |
| SEM1 | SEM1_5 | 0.23 | 0.31 | 13.11 | 10.73 | 5 | 35.49 | 91.00 | 93.48 | 5 | 55 | 216 | 217 |
| SEM2 | SEM2_1 | -0.31 | -0.07 | 3.94 | 4.83 | 1 | 0.00 | 5.00 | 23.66 | 1 | 0 | 0 | 8 |
| SEM2 | SEM2_6 | 0.44 | -0.12 | 7.34 | 7.62 | 6 | 34.32 | 84.00 | 88.91 | 6 | 35 | 199 | 202 |
| SEM2 | SEM2_7 | 0.18 | 0.22 | 2.86 | 4.13 | 7 | 39.81 | 141.00 | 197.46 | 7 | 25 | 150 | 207 |
| SEM2 | SEM2_11 | 0.07 | 0.33 | 3.26 | 4.57 | 11 | 15.80 | 20.00 | 61.20 | 11 | 12 | 15 | 60 |
| SEM3 | SEM3_1 | -0.35 | 0.00 | 4.20 | 3.56 | 1 | 0.00 | 0.00 | 58.59 | 1 | 0 | 0 | 44 |
| SEM3 | SEM3_6 | 0.40 | 0.02 | 5.25 | 4.40 | 6 | 37.91 | 85.00 | 89.20 | 6 | 40 | 199 | 203 |
| SL | SL5 | 5.84 | 3.22 | 10.56 | 6.89 | 5 | 33.34 | 43.00 | 93.48 | 5 | 52 | 170 | 217 |
| SL | SL8 | -5.23 | 3.30 | 4.83 | 3.65 | 8 | 0.00 | 16.00 | 153.86 | 8 | 0 | 155 | 276 |
| SL | SL13 | -4.84 | -1.02 | 7.26 | 5.05 | 13 | 0.00 | 1.18 | 56.48 | 13 | 3 | 3 | 270 |
| SW | SW4 | -0.18 | -0.08 | 4.16 | 3.82 | 4 | 41.94 | 53.00 | 58.42 | 4 | 68 | 171 | 185 |
| SW | SW10 | -0.04 | 0.28 | 4.71 | 3.41 | 10 | 0.49 | 6.22 | 39.68 | 10 | 4 | 12 | 71 |
| SW | SW12 | -0.21 | 0.23 | 5.17 | 4.68 | 12 | 70.14 | 93.00 | 96.82 | 12 | 208 | 213 | 215 |
| SW | SW13 | -0.16 | -0.11 | 4.28 | 4.13 | 13 | 0.00 | 15.38 | 91.81 | 13 | 0 | 33 | 281 |
| SN | SN4 | -0.90 | -0.20 | 5.03 | 4.99 | 4 | 14.62 | 22.00 | 50.78 | 4 | 21 | 36 | 161 |
| SN | SN5 | 1.07 | 0.77 | 9.05 | 8.81 | 5 | 33.34 | 40.59 | 93.48 | 5 | 52 | 170 | 217 |
| SN | SN7 | 0.13 | -1.18 | 4.18 | 4.38 | 7 | 46.61 | 53.12 | 187.55 | 7 | 29 | 35 | 206 |
| SN | SN13 | -0.89 | -0.02 | 4.84 | 5.17 | 13 | 12.45 | 67.00 | 95.79 | 13 | 16 | 278 | 283 |

| | | | | | | | | | | | | | |
|---|---|---|---|---|---|---|---|---|---|---|---|---|---|
| SB | SB11 | 3.47 | 0.32 | 22.49 | 17.5 | 11 | 30.65 | 58.00 | 61.20 | 11 | 23 | 57 | 60 |
| SBA | SBA11 | 0.35 | 0.04 | 22.08 | 17.1 | 11 | 47.97 | 58.00 | 61.20 | 11 | 47 | 57 | 60 |
| MSBR | MSBR5 | 0.32 | -0.07 | 6.57 | 4.66 | 5 | 18.04 | 25.51 | 30.74 | 5 | 22 | 35 | 48 |
| SLW | SLW13 | -3.57 | -4.23 | 11.30 | 3.98 | 13 | 0.00 | 13.36 | 19.33 | 13 | 0 | 18 | 46 |

**5. Rhizome traits**

| Trait | QTL | *a* | *d* | $r^2$ (%) | LOD | LG | cMmin | cMpeak | cMmax | Chrom | bmin | bpeak | bmax |
|---|---|---|---|---|---|---|---|---|---|---|---|---|---|
| RH1 | RH1_12 | -0.03 | -0.12 | 4.09 | 3.87 | 12 | 18.11 | 19.37 | 23.09 | 12 | 35 | 37 | 66 |
| RH2 | RH2_2 | 0.19 | 0.12 | 4.18 | 3.89 | 2 | 87.63 | 153.77 | 173.51 | 2 | 124 | 183 | 205 |
| RH3 | RH3_2 | 0.50 | -0.02 | 6.92 | 6.83 | 2 | 163.83 | 170.00 | 173.51 | 2 | 196 | 203 | 205 |
| RH3 | RH3_14 | 0.39 | -0.10 | 4.07 | 4.19 | 14 | 114.06 | 123.97 | 141.04 | 14 | 248 | 266 | 281 |
| PR2 | PR2_2 | 0.03 | 0.01 | 4.75 | 4.43 | 2 | 125.45 | 157.42 | 160.92 | 2 | 160 | 186 | 191 |
| PR3 | PR3_2 | 0.03 | 0.00 | 5.91 | 4.47 | 2 | 148.83 | 169.00 | 173.51 | 2 | 182 | 202 | 205 |
| PR3 | PR3_5 | 0.04 | -0.01 | 6.01 | 4.54 | 5 | 32.72 | 52.00 | 88.49 | 5 | 50 | 170 | 214 |

**C:** EAM × OPB

**3. Size traits**

| Trait | QTL | *a* | *d* | $r^2$ (%) | LOD | LG | cMmin | cMpeak | cMmax | Chrom | bmin | bpeak | bmax |
|---|---|---|---|---|---|---|---|---|---|---|---|---|---|
| LW | LW12 | 2.52 | 4.07 | 2.55 | 3.13 | 12 | 6.89 | 11.00 | 17.16 | 12 | 170 | 179 | 199 |
| IW | IW2 | -1.58 | 0.22 | 4.82 | 4.38 | 2 | 21.66 | 23.88 | 26.10 | 2 | 162 | 163 | 184 |
| IW | IW4 | -0.82 | -0.50 | 4.69 | 4.21 | 4 | 0.00 | 0.00 | 3.33 | 4 | 0 | 0 | 46 |

**4. Stolon traits**

| Trait | QTL | *a* | *d* | $r^2$ (%) | LOD | LG | cMmin | cMpeak | cMmax | Chrom | bmin | bpeak | bmax |
|---|---|---|---|---|---|---|---|---|---|---|---|---|---|
| ST1 | ST1_5 | 1.71 | 0.47 | 2.72 | 3.48 | 5 | 31.47 | 33.69 | 33.69 | 5 | 206 | 217 | 217 |
| ST3 | ST3_13 | 3.18 | -0.78 | 2.76 | 3.42 | 13 | 31.44 | 36.00 | 37.39 | 13 | 226 | 278 | 307 |
| SEM1 | SEM1_5 | 0.62 | 0.29 | 3.20 | 4.02 | 5 | 8.65 | 33.69 | 33.69 | 5 | 68 | 217 | 217 |
| SW | SW14 | 0.68 | -0.09 | 3.72 | 4.48 | 14 | 0.00 | 0.00 | 2.67 | 14 | 0 | 0 | 31 |
| SBA | SBA11 | 0.04 | 0.22 | 2.81 | 3.29 | 11 | 2.78 | 14.00 | 31.13 | 11 | 58 | 90 | 241 |
| SLW | SLW4 | 4.96 | -4.84 | 4.62 | 3.47 | 4 | 4.43 | 32.33 | 34.49 | 4 | 54 | 197 | 198 |

**5. Rhizome traits**

| Trait | QTL | *a* | *d* | $r^2$ (%) | LOD | LG | cMmin | cMpeak | cMmax | Chrom | bmin | bpeak | bmax |
|---|---|---|---|---|---|---|---|---|---|---|---|---|---|
| REM2 | REM2_12 | 0.02 | -0.50 | 2.73 | 3.24 | 12 | 0.00 | 0.00 | 12.58 | 12 | 0 | 0 | 179 |

**D:** IMPO × OPB

**1. Floral traits**

| Trait | QTL | *a* | *d* | $r^2$ (%) | LOD | LG | cMmin | cMpeak | cMmax | Chrom | bmin | bpeak | bmax |
|---|---|---|---|---|---|---|---|---|---|---|---|---|---|
| CTL | CTL1 | 0.77 | -0.08 | 4.55 | 1.72 | 1 | 0.00 | 1.72 | 16.26 | 1 | 0 | 5 | 25 |
| CTL | CTL5 | 1.14 | 0.16 | 9.41 | 38.88 | 5 | 11.71 | 38.88 | 50.04 | 5 | 13 | 44 | 217 |
| CTL | CTL7 | -0.64 | 0.26 | 3.22 | 0.65 | 7 | 0.00 | 0.65 | 11.41 | 7 | 0 | 2 | 14 |
| CTL | CTL8 | -0.86 | -0.02 | 3.29 | 16.00 | 8 | 3.06 | 16.00 | 34.81 | 8 | 6 | 77 | 175 |
| CTL | CTL10 | 0.76 | 0.06 | 3.69 | 40.30 | 10 | 12.86 | 40.30 | 43.37 | 10 | 18 | 89 | 170 |
| CTL | CTL14 | -1.21 | -0.11 | 10.54 | 83.00 | 14 | 79.36 | 83.00 | 93.69 | 14 | 241 | 245 | 265 |
| CLL | CLL2 | 1.38 | -0.20 | 9.06 | 10.37 | 2 | 63.33 | 68.19 | 72.29 | 2 | 185 | 192 | 197 |
| CLL | CLL6 | -0.74 | 0.33 | 2.51 | 3.96 | 6 | 12.68 | 21.69 | 40.94 | 6 | 21 | 33 | 61 |
| CLL | CLL11 | 0.94 | 0.05 | 4.48 | 5.43 | 11 | 30.87 | 48.19 | 70.21 | 11 | 69 | 233 | 328 |
| CLL | CLL14 | -0.90 | -0.05 | 3.72 | 4.91 | 14 | 14.43 | 98.41 | 114.58 | 14 | 34 | 274 | 282 |
| CLW | CLW2 | 1.03 | -0.23 | 5.05 | 5.09 | 2 | 63.33 | 70.92 | 77.65 | 2 | 185 | 194 | 205 |
| CLW | CLW5 | 1.47 | 0.45 | 10.40 | 10.04 | 5 | 24.56 | 38.88 | 50.04 | 5 | 24 | 44 | 173 |
| CLW | CLW6 | -0.79 | 0.79 | 3.43 | 3.83 | 6 | 6.20 | 15.00 | 22.94 | 6 | 13 | 24 | 35 |

**2. Phenological traits**

| Trait | QTL | *a* | *d* | $r^2$ (%) | LOD | LG | cMmin | cMpeak | cMmax | Chrom | bmin | bpeak | bmax |
|---|---|---|---|---|---|---|---|---|---|---|---|---|---|
| FN | FN1 | -0.21 | 0.26 | 1.99 | 3.64 | 1 | 13.64 | 26.16 | 48.32 | 1 | 23 | 38 | 72 |
| FN | FN6 | 0.55 | -0.10 | 7.75 | 9.83 | 6 | 66.03 | 66.00 | 66.69 | 6 | 195 | 206 | 208 |
| FN | FN8 | 0.67 | -0.26 | 10.32 | 12.89 | 8 | 69.54 | 78.77 | 80.21 | 8 | 254 | 273 | 274 |
| FN | FN14 | 0.44 | -0.18 | 5.25 | 6.04 | 14 | 83.74 | 87.87 | 92.53 | 14 | 246 | 253 | 263 |
| FT | FT5 | -2.14 | -2.81 | 8.81 | 9.92 | 5 | 41.65 | 47.28 | 50.04 | 5 | 55 | 80 | 217 |
| FT | FT8 | 2.07 | -0.72 | 3.66 | 4.27 | 8 | 66.72 | 78.77 | 80.49 | 8 | 245 | 273 | 276 |
| FT | FT14 | 1.76 | -0.62 | 3.24 | 3.49 | 14 | 74.72 | 82.03 | 114.58 | 14 | 229 | 242 | 282 |
| IL | IL3 | 1.50 | -0.25 | 3.61 | 4.74 | 3 | 13.30 | 22.00 | 25.31 | 3 | 196 | 211 | 212 |
| IL | IL8 | -1.83 | -0.99 | 7.75 | 9.10 | 8 | 3.06 | 31.85 | 47.38 | 8 | 6 | 164 | 204 |

**3. Size traits**

| Trait | QTL | *a* | *d* | $r^2$ (%) | LOD | LG | cMmin | cMpeak | cMmax | Chrom | bmin | bpeak | bmax |
|---|---|---|---|---|---|---|---|---|---|---|---|---|---|
| LL | LL4 | 3.30 | -1.58 | 6.72 | 6.06 | 4 | 3.15 | 7.00 | 15.09 | 4 | 8 | 13 | 27 |
| LL | LL5 | 1.63 | 4.59 | 7.27 | 7.40 | 5 | 45.28 | 47.28 | 50.04 | 5 | 72 | 80 | 217 |
| LL | LL10 | 4.10 | 0.83 | 7.54 | 7.82 | 10 | 12.86 | 42.11 | 47.68 | 10 | 18 | 158 | 183 |
| LL | LL14 | 2.59 | -1.35 | 3.72 | 3.74 | 14 | 23.49 | 69.05 | 75.85 | 14 | 50 | 217 | 230 |
| LW | LW2 | -1.67 | -0.94 | 3.50 | 3.54 | 2 | 54.06 | 63.33 | 77.65 | 2 | 171 | 185 | 205 |
| LW | LW5 | 2.12 | 3.61 | 12.66 | 13.03 | 5 | 46.96 | 50.04 | 50.04 | 5 | 78 | 217 | 217 |
| LW | LW10 | 2.39 | 1.06 | 5.73 | 5.59 | 10 | 12.86 | 42.11 | 50.75 | 10 | 18 | 158 | 194 |
| IW | IW5 | 0.37 | 0.47 | 7.63 | 8.68 | 5 | 43.26 | 50.04 | 50.04 | 5 | 58 | 217 | 217 |
| IW | IW13 | -0.35 | -0.00 | 3.12 | 3.86 | 13 | 27.77 | 77.85 | 84.97 | 13 | 202 | 250 | 307 |
| IW | IW14 | 0.37 | 0.23 | 4.89 | 6.06 | 14 | 17.33 | 25.43 | 40.13 | 14 | 35 | 55 | 80 |

**4. Stolon traits**

| Trait | QTL | *a* | *d* | $r^2$ (%) | LOD | LG | cMmin | cMpeak | cMmax | Chrom | bmin | bpeak | bmax |
|---|---|---|---|---|---|---|---|---|---|---|---|---|---|
| ST1 | ST1_5 | 1.04 | 0.90 | 18.62 | 21.82 | 5 | 46.11 | 48.00 | 50.04 | 5 | 76 | 125 | 173 |
| ST1 | ST1_11 | 0.45 | 0.04 | 2.74 | 3.58 | 11 | 0.00 | 25.02 | 66.00 | 11 | 0 | 52 | 321 |
| ST2 | ST2_11 | 0.78 | 0.58 | 3.80 | 3.66 | 11 | 0.00 | 22.57 | 73.55 | 11 | 0 | 34 | 332 |
| ST2 | ST2_13 | 1.52 | 0.18 | 8.04 | 8.33 | 13 | 33.93 | 44.37 | 82.89 | 13 | 230 | 243 | 304 |
| ST2 | ST2_14 | 1.00 | 0.16 | 4.30 | 3.68 | 14 | 40.13 | 65.46 | 94.08 | 14 | 80 | 210 | 266 |
| ST3 | ST3_13 | 1.59 | -0.46 | 8.62 | 6.67 | 13 | 27.77 | 39.00 | 80.75 | 13 | 202 | 239 | 302 |
| SEM1 | SEM1_5 | 0.24 | 0.27 | 21.75 | 22.4 | 5 | 47.72 | 49.72 | 50.04 | 5 | 125 | 171 | 217 |
| SEM2 | SEM2_6 | 0.42 | -0.17 | 8.85 | 8.75 | 6 | 57.69 | 66.69 | 66.69 | 6 | 194 | 208 | 208 |
| SEM2 | SEM2_8 | 0.27 | -0.01 | 3.39 | 3.74 | 8 | 65.51 | 78.77 | 80.49 | 8 | 243 | 273 | 276 |
| SEM2 | SEM2_13 | 0.29 | -0.12 | 4.61 | 3.81 | 13 | 32.94 | 44.37 | 84.97 | 13 | 224 | 243 | 307 |
| SEM3 | SEM3_6 | 0.35 | 0.20 | 5.21 | 3.73 | 6 | 45.84 | 60.82 | 66.69 | 6 | 165 | 197 | 208 |
| SEM3 | SEM3_13 | 0.48 | -0.02 | 6.26 | 4.57 | 13 | 19.41 | 44.85 | 84.97 | 13 | 35 | 245 | 307 |
| SL | SL6 | 3.78 | 0.31 | 5.14 | 4.70 | 6 | 29.51 | 62.27 | 66.69 | 6 | 42 | 200 | 208 |
| SL | SL14 | -3.20 | 0.61 | 3.52 | 3.51 | 14 | 69.05 | 78.40 | 89.97 | 14 | 217 | 238 | 255 |
| SW | SW7 | -0.17 | 0.06 | 5.02 | 3.58 | 7 | 0.00 | 3.38 | 12.55 | 7 | 0 | 6 | 15 |
| SN | SN5 | 1.00 | 0.59 | 6.91 | 6.02 | 5 | 36.81 | 40.52 | 50.04 | 5 | 40 | 49 | 217 |
| SN | SN11 | -0.96 | 0.21 | 5.29 | 4.82 | 11 | 34.44 | 67.57 | 73.87 | 11 | 106 | 323 | 334 |
| SB | SB6 | 1.73 | -0.66 | 8.22 | 7.14 | 6 | 45.51 | 65.00 | 66.69 | 6 | 163 | 205 | 208 |
| SB | SB8 | 1.53 | -0.36 | 5.43 | 4.88 | 8 | 59.40 | 66.72 | 80.49 | 8 | 238 | 245 | 276 |
| SB | SB9 | -1.32 | 0.47 | 3.80 | 4.26 | 9 | 0.00 | 2.16 | 5.47 | 9 | 0 | 4 | 10 |
| SB | SB13 | 1.43 | 0.52 | 4.54 | 4.67 | 13 | 80.28 | 83.89 | 84.97 | 13 | 301 | 305 | 307 |
| SBA | SBA6 | 0.17 | -0.03 | 7.50 | 5.23 | 6 | 35.92 | 65.00 | 66.69 | 6 | 49 | 205 | 208 |
| SBA | SBA8 | 0.12 | -0.00 | 3.81 | 3.72 | 8 | 53.28 | 67.86 | 80.49 | 8 | 225 | 248 | 276 |
| SBA | SBA11 | 0.12 | 0.03 | 4.29 | 4.38 | 11 | 16.94 | 50.65 | 71.95 | 11 | 26 | 248 | 330 |
| SBA | SBA13 | 0.14 | -0.02 | 4.37 | 4.35 | 13 | 44.50 | 83.89 | 84.97 | 13 | 244 | 305 | 307 |

### 5. Rhizome traits

| Trait | QTL | *a* | *d* | $r^2$ (%) | LOD | LG | cMmin | cMpeak | cMmax | Chrom | bmin | bpeak | bmax |
|---|---|---|---|---|---|---|---|---|---|---|---|---|---|
| RH1 | RH1_11 | 0.08 | -0.03 | 3.48 | 3.59 | 11 | 3.29 | 17.59 | 73.87 | 11 | 7 | 27 | 334 |
| RH2 | RH2_11 | 0.40 | -0.08 | 8.86 | 9.34 | 11 | 16.40 | 50.65 | 61.78 | 11 | 25 | 248 | 318 |
| RH3 | RH3_11 | 0.77 | -0.21 | 17.57 | 15.5 | 11 | 15.01 | 22.98 | 27.92 | 11 | 23 | 45 | 58 |
| REM3 | REM3_6 | 0.36 | -0.34 | 7.65 | 3.97 | 6 | 13.73 | 20.17 | 60.03 | 6 | 22 | 30 | 195 |
| REM3 | REM3_11 | 0.44 | -0.02 | 8.74 | 4.44 | 11 | 8.48 | 16.40 | 56.00 | 11 | 13 | 25 | 280 |
| RL | RL11 | 6.22 | 8.46 | 10.62 | 3.61 | 11 | 0.00 | 4.67 | 11.72 | 11 | 0 | 8 | 18 |
| RB | RB8 | 2.81 | -0.89 | 12.12 | 4.01 | 8 | 36.58 | 46.94 | 52.31 | 8 | 178 | 202 | 223 |
| RLL | RLL6 | 8.93 | -7.30 | 41.19 | 7.86 | 6 | 21.69 | 29.04 | 47.62 | 6 | 33 | 41 | 173 |
| RLW | RLW6 | 4.36 | -2.19 | 28.25 | 4.90 | 6 | 21.69 | 47.00 | 60.82 | 6 | 33 | 171 | 197 |
| PR1 | PR1_9 | 0.02 | -0.01 | 4.52 | 3.72 | 9 | 35.35 | 52.44 | 54.18 | 9 | 225 | 248 | 251 |
| PR1 | PR1_11 | 0.02 | -0.01 | 4.31 | 3.52 | 11 | 3.29 | 17.59 | 73.87 | 11 | 7 | 27 | 334 |
| PR2 | PR2_11 | 0.03 | -0.01 | 8.11 | 8.47 | 11 | 16.40 | 50.65 | 61.78 | 11 | 25 | 248 | 318 |
| PR3 | PR3_11 | 0.06 | -0.02 | 20.23 | 16.6 | 11 | 15.01 | 22.98 | 27.92 | 11 | 23 | 45 | 58 |

**Table S7. Epistatic interactions between QTLs influencing measured life-history traits across the four mapping populations.** The "P/T cat" denotes the population and trait category. F: floral traits, P: phenological traits, S: size traits, T: stolon traits, R: rhizome traits. Epistatic effects are classified into four types: *aa* (additive-additive)*, da* (dominance-additive)*, ad* (additive-dominance), and *dd* (dominance-dominance). Significant level: . < 0.1, ∗ p < 0.05, ∗∗ p < 0.01, ∗∗∗< 0.001.

| P/T cat | trait | QTL_i | QTL_j | *aa* | *da* | *ad* | *dd* | *P* value |
|---|---|---|---|---|---|---|---|---|
| LVR/F | CTL | CTL2 | CTL10 | 0.19 | 0.46 | 1.21 | -0.70 | 0.0192 * |
| | | CTL5 | CTL6 | -0.12 | 0.70 | -0.77 | 0.54 | 0.0216 * |
| | CLL | CLL5 | CLL10 | -1.39 | -0.04 | 0.83 | 0.84 | 0.0254 * |
| | | CLL5 | CLL13 | 0.95 | -0.30 | 0.02 | 1.49 | 0.068006 . |
| | | CLL8 | CLL13 | 1.18 | -0.47 | 0.27 | -0.55 | 0.042 * |
| | | CLL11 | CLL13 | 0.22 | -1.40 | -0.61 | 0.75 | 0.061712 . |
| | | CLL11 | CLL14 | -1.11 | -1.20 | 1.43 | 0.56 | 0.00571 ** |
| | | CLL12 | CLL13 | -0.41 | -0.85 | 1.38 | 0.36 | 0.0363 * |
| LVR/P | FN | FN5 | FN8 | -0.39 | -0.28 | 0.22 | 0.43 | 0.00446 ** |
| | | FN6 | FN8 | -0.34 | -0.15 | 0.26 | -0.29 | 0.026 * |
| | | FN7 | FN8 | -0.10 | -0.49 | -0.35 | 0.22 | 0.0241 * |
| | | FN8 | FN12 | 0.23 | 0.08 | -0.66 | 0.09 | 0.00273 ** |
| | FT | FT4 | FT5 | 2.72 | 0.96 | 2.06 | 1.91 | 7.23e-06 *** |
| | | FT4 | FT6 | 2.41 | 3.24 | -1.30 | -1.18 | 0.000754 *** |
| | | FT5 | FT7 | -3.04 | 0.88 | -0.99 | 1.83 | 0.0176 * |
| | | FT5 | FT8 | -0.55 | -1.49 | 2.57 | 1.01 | 0.00741 ** |
| | | FT6 | FT7 | 0.54 | -1.42 | 1.64 | 1.64 | 0.053 . |
| | | FT7 | FT14 | -1.18 | -0.42 | -0.56 | 3.45 | 0.08489 . |
| | IL | IL4 | IL8 | -2.23 | 0.41 | 0.08 | -0.24 | 0.0339 * |
| | | IL4 | IL14 | -1.53 | 1.36 | -1.26 | 0.64 | 0.0505 . |
| | | IL6 | IL7 | -1.19 | -2.18 | -0.34 | -3.89 | 0.0221 * |
| | | IL7 | IL8 | 3.30 | 0.38 | 1.93 | 0.63 | 0.000031 *** |
| | | IL8 | IL14 | 1.87 | 1.34 | 0.88 | 0.14 | 0.0286 * |
| LVR/S | LL | LL4 | LL5 | -1.29 | 0.63 | 1.51 | 6.35 | 0.0292 * |
| | | LL5 | LL11 | 1.50 | 3.25 | -1.65 | -2.53 | 0.064221 . |
| | IW | IW2 | IW7 | -0.10 | -0.24 | 0.03 | -0.45 | 0.099420 . |
| | | IW2 | IW12 | 0.28 | -0.14 | -0.35 | -0.05 | 0.0532 . |
| LVR/T | ST1 | ST1_5 | ST1_13 | 0.08 | 0.44 | 0.06 | 0.46 | 0.0757 . |
| | | ST1_6 | ST1_14 | 0.46 | -0.25 | -0.07 | 0.20 | 0.0529 . |
| | ST2 | ST2_13 | ST2_14 | 0.34 | -0.21 | 0.43 | -0.35 | 0.0597 . |
| | ST3 | ST3_5 | ST3_8 | 0.81 | -0.03 | -0.46 | -0.37 | 0.0603 . |
| | | ST3_5 | ST3_14 | -0.27 | 0.45 | -0.17 | -1.19 | 0.0948 . |
| | SL | SL8 | SL9 | 4.28 | -1.62 | -5.10 | 1.40 | 0.00403 ** |
| | SEM2 | SEM2_5 | SEM2_7 | -0.34 | -0.04 | -0.15 | 0.11 | 0.00327 ** |
| | | SEM2_5 | SEM2_8 | -0.18 | -0.21 | 0.13 | 0.20 | 0.0139 * |
| | | SEM2_7 | SEM2_8 | 0.03 | -0.05 | -0.34 | 0.05 | 0.0285 * |
| | SEM3 | SEM3_8 | SEM3_12 | -0.22 | 0.20 | 0.14 | -0.18 | 0.0521 . |
| | SN | SN4 | SN5 | -1.17 | 0.62 | 0.06 | -0.10 | 0.079097 . |
| CWF/F | CTL | CTL3 | CTL9 | -0.30 | 0.67 | -0.66 | -0.20 | 0.08926 . |
| | CTL | CTL4 | CTL12 | 0.02 | -0.81 | -0.42 | 0.96 | 0.0732 . |
| | CTL | CTL5 | CTL12 | 0.75 | 0.52 | -0.50 | -0.36 | 0.0366 * |
| | CLW | CLW9 | CLW14 | 0.94 | -2.45 | -0.16 | 1.19 | 0.00311 ** |
| CWF/P | FN | FN4 | FN14 | 0.10 | -0.61 | -0.20 | 0.86 | 0.00671 ** |
| | FN | FN6 | FN14 | 0.02 | 0.57 | 0.06 | -0.65 | 0.05580 . |
| | FT | FT5 | FT6 | 0.98 | 2.64 | 2.25 | 1.39 | 0.00649 ** |
| | FT | FT5 | FT14 | -2.05 | 3.32 | 0.52 | -5.83 | 0.000713 *** |
| CWF/T | ST2 | ST2_6 | ST2_14 | 0.12 | 1.20 | 0.73 | -1.89 | 0.0386 * |
| | SW | SW12 | SW13 | 0.04 | 0.01 | 0.05 | -0.40 | 0.081375 . |
| | SN | SN4 | SN7 | -0.46 | -0.23 | 1.20 | 0.12 | 0.042 * |
| | SN | SN4 | SN13 | 1.03 | 0.24 | 0.24 | -0.67 | 0.00716 ** |
| CWF/R | RH3 | RH3_2 | RH3_14 | 0.29 | -0.19 | -0.27 | 0.56 | 0.0291 * |
| | PR3 | PR3_2 | PR3_5 | 0.05 | 0.00 | -0.04 | 0.06 | 0.00*** |
| IMPO/F | CTL | CTL5 | CTL7 | -0.37 | 0.50 | -0.66 | 0.58 | 0.0431 * |

| | | | | | | | | |
|---|---|---|---|---|---|---|---|---|
| | CTL | CTL5 | CTL8 | -0.55 | 0.38 | -0.96 | 0.51 | 0.0833 . |
| | CTL | CTL5 | CTL14 | 0.16 | -0.32 | 0.46 | -1.13 | 0.0932 . |
| | CTL | CTL7 | CTL10 | -0.57 | -0.45 | -0.06 | 0.68 | 0.0564 . |
| | CLW | CLW5 | CLW6 | 0.11 | 0.22 | 0.89 | -1.48 | 0.0537 . |
| IMPO/P | FT | FT5 | FT14 | -0.53 | 0.13 | 0.34 | -3.52 | 0.083624 . |
| IMPO/T | SEM3 | SEM3_6 | SEM3_13 | 0.37 | -0.32 | -0.28 | 0.17 | 0.00838 ** |
| | PR1 | PR1_9 | PR1_11 | 0.02 | -0.01 | -0.00 | -0.01 | 0.0092 ** |

**Table S8. Average degree of QTL overlap (mean ± SE) within and between trait categories across the three mapping populations.** A) The coastal *M. guttatus* × *M. tilingii* population. B) The coastal *M. guttatus* × CWF population. C) The coastal *M. guttatus* × *M. decorus* population. Significance levels: ∗ $p < 0.05$, ∗∗ $p < 0.01$, ∗∗∗< 0.001, determined by permutation test with 1000 permutations.

A

| **Pair of trait category** | **Average correlation coefficients (mean ± SE)** | ***p* values** |
|---|---|---|
| Floral – Floral | 0.22 ± 0.06 | 0.169 |
| Floral – Phenology | 0.20 ± 0.05 | |
| Floral – Size | 0.23 ± 0.03 | |
| Floral – Stolon | 0.12 ± 0.03 | |
| Floral – Rhizome | 0.03 ± 0.01 | |
| Phenology – Phenology | 0.40 ± 0.09 | 0.054 |
| Phenology – Size | 0.23 ± 0.05 | |
| Phenology – Stolon | 0.18 ± 0.03 | |
| Phenology – Rhizome | 0 ± 0 | |
| Size – Size | 0.30 ± 0.03 | 0.097 |
| Size – Stolon | 0.15 ± 0.03 | |
| Size – Rhizome | 0.06 ± 0.02 | |
| Stolon – Stolon | 0.10 ± 0.03 | 0.662 |
| Stolon – Rhizome | 0.04 ± 0.02 | |
| Rhizome – Rhizome | 0.18 ± 0.06 | 0.001** |

B

| **Pair of trait category** | **Average correlation coefficients (mean ± SE)** | ***p* values** |
|---|---|---|
| Floral – Floral | 0.27 ± 0.05 | 0.132 |
| Floral – Phenology | 0.23 ± 0.03 | |
| Floral – Size | 0.21 ± 0.05 | |
| Floral – Stolon | 0.11 ± 0.03 | |
| Floral – Rhizome | 0.11 ± 0.03 | |
| Phenology – Phenology | 0.17 ± 0.10 | 0.448 |
| Phenology – Size | 0.14 ± 0.04 | |
| Phenology – Stolon | 0.09 ± 0.03 | |
| Phenology – Rhizome | 0.24 ± 0.09 | |
| Size – Size | 0.5 ± 0.25 | 0.007** |

| | | |
|---|---|---|
| Size – Stolon | 0.13 ± 0.03 | |
| Size – Rhizome | 0.15 ± 0.07 | |
| Stolon – Stolon | 0.09 ± 0.03 | 0.433 |
| Stolon – Rhizome | 0.03± 0.02 | |
| Rhizome – Rhizome | 0.33 ± NA | 0.12 |

C

| **Pair of trait category** | **Average correlation coefficients (mean ± SE)** | ***p* values** |
|---|---|---|
| Floral – Floral | 0.21 ± 0.09 | 0.197 |
| Floral – Phenology | 0.13 ± 0.04 | |
| Floral – Size | 0.22 ± 0.04 | |
| Floral – Stolon | 0.08 ± 0.03 | |
| Floral – Rhizome | 0.09 ± 0.03 | |
| Phenology – Phenology | 0.13 ± 0.13 | 0.286 |
| Phenology – Size | 0.09 ± 0.05 | |
| Phenology – Stolon | 0.10 ± 0.03 | |
| Phenology – Rhizome | 0.04 ± 0.03 | |
| Size – Size | 0.33 ± 0.07 | 0.041* |
| Size – Stolon | 0.09 ± 0.03 | |
| Size – Rhizome | 0 ± 0 | |
| Stolon – Stolon | 0.17 ± 0.05 | 0.072 |
| Stolon – Rhizome | 0.11 ± 0.02 | |
| Rhizome – Rhizome | 0.21 ± 0.07 | 0.005** |

**Table S9. Average degree of QTL overlap (mean ± SE) between trait categories of stolon-related traits (the number of stolons and the highest node of stolon emergence) across the three mapping populations, consistently measured at three developmental stages.** Trait categories are defined based on developmental stages: vegetative, early reproductive, and late reproductive. A) The coastal *M. guttatus* × *M. tilingii* population. B) The coastal *M.*

*guttatus* × CWF population. C) The coastal *M. guttatus* × *M. decorus* population. Significance levels were determined by a permutation test with 1000 permutations.

A

| Trait category pair for stolon traits | Average correlation coefficients (mean ± SE) | *p* values |
|---|---|---|
| Vegetative – vegetative | 0.5 ± NA | 0.219 |
| vegetative – early reproductive | 0.21 ± 0.05 | |
| vegetative – late reproductive | 0.34 ± 0.06 | |
| early reproductive – early reproductive | 0.29 ± NA | 0.605 |
| early reproductive – late reproductive | 0.63 ± 0.15 | |
| late reproductive – late reproductive | 0.57 ± NA | 0.383 |

B

| Trait category pair for stolon traits | Average correlation coefficients (mean ± SE) | *p* values |
|---|---|---|
| vegetative – vegetative | 0.33 ± NA | 0.215 |
| vegetative – early reproductive | 0 ± 0 | |
| vegetative – late reproductive | 0 ± 0 | |
| early reproductive – early reproductive | 0.5 ± NA | 0.285 |
| early reproductive – late reproductive | 0.42 ± 0.09 | |
| late reproductive – late reproductive | 0.25 ± NA | 0.401 |

C

| Trait category pair for stolon traits | Average correlation coefficients (mean ± SE) | *p* values |
|---|---|---|
| vegetative – vegetative | 0.5 ± NA | 0.127 |
| vegetative – early reproductive | 0.06 ± 0.06 | |
| vegetative – late reproductive | 0 ± 0 | |
| early reproductive – early reproductive | 0.2 ± NA | 0.555 |
| early reproductive – late reproductive | 0.40 ± 0.09 | |
| late reproductive – late reproductive | 0.5 ± NA | 0.193 |

**Table S10. Average degree of QTL overlap (mean ± SE) between trait categories of rhizome-related traits (the number of rhizomes, the highest node of rhizome emergence, and the proportion of stems that are rhizomes) across the three mapping populations, consistently measured at three developmental stages.** Trait categories are defined based on developmental stages: vegetative, early reproductive, and late reproductive. A) The coastal *M.*

*guttatus* × *M. tilingii* population. B) The coastal *M. guttatus* × CWF population. C) The coastal *M. guttatus* × *M. decorus* population. Significance levels were determined by permutation test with 1000 permutations. *** $p < 0.001$

A

| **Trait category pair for rhizome traits** | **Average correlation coefficients (mean ± SE)** | ***p* values** |
|---|---|---|
| Vegetative – vegetative | 0.33 ± 0.33 | 0.113 |
| vegetative – early reproductive | 0 ± 0 | |
| vegetative – late reproductive | 0 ± 0 | |
| early reproductive – early reproductive | 0 ± NA | 0.728 |
| early reproductive – late reproductive | 0.25 ± 0.25 | |
| late reproductive – late reproductive | NA ± NA | NA |

B

| **Trait category pair for rhizome traits** | **Average correlation coefficients (mean ± SE)** | ***p* values** |
|---|---|---|
| vegetative – vegetative | NA ± NA | NA |
| vegetative – early reproductive | 0 ± 0 | |
| vegetative – late reproductive | 0 ± 0 | |
| early reproductive – early reproductive | 1 ± NA | 0*** |
| early reproductive – late reproductive | 0.38 ± 0.13 | |
| late reproductive – late reproductive | 0.33 ± NA | 0.312 |

C

| **Trait category pair for rhizome traits** | **Average correlation coefficients (mean ± SE)** | ***p* values** |
|---|---|---|
| vegetative – vegetative | 0.5 ± NA | 1 |
| vegetative – early reproductive | 0.75 ± 0.14 | |
| vegetative – late reproductive | 0.64 ± 0.12 | |
| early reproductive – early reproductive | 1 ± NA | 0*** |
| early reproductive – late reproductive | 0.83 ± 0.11 | |
| late reproductive – late reproductive | 0.67 ± 0.17 | 0.864 |

A

**Table S11. Number of QTL overlaps ($n_{12}$), degree of QTL overlap, and *p*-values between the same traits for each pair of mapping populations: CWF/LVR (A), IMPO/LVR (B), and IMPO/CWF (C).** Significance levels were determined by permutation test with 1000 permutations.

A

| Traits | $n_{12}$ | Overlap degree | *p*_value |
|---|---|---|---|
| FN | 1 | 0.125 | 0.516 |
| CTL | 2 | 0.15384615 | 0.263 |
| CLL | 3 | 0.33333333 | 0.107 |
| CLW | 0 | 0 | 1 |
| FT | 2 | 0.25 | 0.279 |
| IL | 2 | 0.33333333 | 0.251 |
| IW | 1 | 0.11111111 | 0.443 |
| LL | 0 | 0 | 1 |
| LW | 1 | 0.11111111 | 0.396 |
| ST1 | 1 | 0.16666667 | 0.495 |
| ST2 | 1 | 0.125 | 0.569 |
| ST3 | 1 | 0.125 | 0.494 |
| SEM1 | 1 | 0.2 | 0.525 |
| SEM2 | 1 | 0.125 | 0.567 |
| SEM3 | 0 | 0 | 1 |
| SL | 1 | 0.25 | 0.596 |
| SW | 0 | 0 | 1 |
| SN | 2 | 0.4 | 0.362 |
| SB | 0 | 0 | 1 |
| SBA | 0 | 0 | 1 |
| MSBR | 0 | 0 | 1 |
| RH1 | 0 | 0 | 1 |
| RH2 | 0 | 0 | 1 |
| RH3 | 0 | 0 | 1 |
| PR2 | 0 | 0 | 1 |

B

| Traits | $n_{12}$ | Overlap degree | *p*_value |
|---|---|---|---|
| FN | 2 | 0.28571429 | 0.309 |
| CTL | 3 | 0.33333333 | 0.168 |
| CLL | 2 | 0.2 | 0.231 |
| CLW | 1 | 0.14285714 | 0.556 |
| FT | 2 | 0.28571429 | 0.227 |
| IL | 0 | 0 | 1 |
| IW | 0 | 0 | 1 |
| LL | 1 | 0.1 | 0.514 |
| LW | 0 | 0 | 1 |
| ST1 | 1 | 0.16666667 | 0.47 |
| ST2 | 2 | 0.4 | 0.33 |
| ST3 | 1 | 0.16666667 | 0.275 |
| SEM1 | 1 | 0.25 | 0.378 |
| SEM2 | 1 | 0.14285714 | 0.543 |
| SEM3 | 0 | 0 | 1 |
| SL | 0 | 0 | 1 |
| SW | 0 | 0 | 1 |
| SN | 1 | 0.25 | 0.604 |
| SB | 2 | 0.5 | 0.294 |
| SBA | 1 | 0.25 | 0.407 |
| RH1 | 0 | 0 | 1 |
| RH2 | 0 | 0 | 1 |
| RH3 | 0 | 0 | 1 |
| RL | 0 | 0 | 1 |
| RB | 0 | 0 | 1 |

| | | | |
|---|---|---|---|
| RLL | 0 | 0 | 1 |
| RLW | 0 | 0 | 1 |
| PR1 | 0 | 0 | 1 |
| PR2 | 0 | 0 | 1 |

C

| Traits | $n_{12}$ | Overlap degree | *p*_value |
|---|---|---|---|
| FN | 1 | 0.14285714 | 0.61 |
| CTL | 4 | 0.36363636 | 0.054 |
| CLL | 0 | 0 | 1 |
| CLW | 1 | 0.125 | 0.497 |
| FT | 2 | 0.4 | 0.336 |
| IL | 1 | 0.25 | 0.608 |
| IW | 2 | 0.5 | 0.408 |
| LL | 1 | 0.2 | 0.544 |
| LW | 1 | 0.25 | 0.587 |
| ST1 | 1 | 0.33333333 | 0.666 |
| ST2 | 2 | 0.33333333 | 0.296 |
| ST3 | 0 | 0 | 1 |
| SEM1 | 1 | 0.5 | 0.67 |
| SEM2 | 1 | 0.16666667 | 0.544 |
| SEM3 | 1 | 0.33333333 | 0.677 |
| SL | 0 | 0 | 1 |
| SW | 0 | 0 | 1 |
| SN | 1 | 0.2 | 0.504 |
| SB | 0 | 0 | 1 |
| SBA | 1 | 0.25 | 0.395 |
| RH1 | 0 | 0 | 1 |
| RH2 | 0 | 0 | 1 |
| RH3 | 0 | 0 | 1 |
| PR2 | 0 | 0 | 1 |
| PR3 | 0 | 0 | 1 |

**Table S12. Summary of candidate genes.** Shown are QTL intervals ≤ 1 Mb in length; strong candidate genes are indicated in purple.

| QTL | Candidate_locus1 | Annotations 1 | Candidate_locus2 | Annotations 2 |
|---|---|---|---|---|

| | | | | |
|---|---|---|---|---|
| LVR_FN8 | Migut.08G237000 | auxin-induced protein 5NG4, putative, expressed (Zhao 2010) | Migut.08G236200 | CESA3 - cellulose synthase, expressed (Zhou et al. 2024) |
| LVR_CLL7 | Migut.07G084200 | OsFBX433 - F-box domain containing protein, expressed (Borna et al. 2022) | | |
| LVR_CLW13 | Migut.13G001200 | polygalacturonase, putative, expressed (Xiao et al. 2014; Önder et al. 2023) | | |
| LVR_FT7 | Migut.07G084200 | OsFBX433 - F-box domain containing protein, expressed (Mizoguchi and Coupland 2000) | Migut.07G085800 | GEM-ASSOCIATED PROTEIN 2 (spliceosome protein-related) (Schlaen et al. 2015) |
| LVR_IW13 | Migut.13G003500 | cyclin, putative, expressed (Umeda 2013) | | |
| LVR_ST1_7 | Migut.07G084200 | OsFBX433 - F-box domain containing protein, expressed (L. Guo et al. 2021) | | |
| LVR_ST1_13 | Migut.13G178800 | OsFBX48 - F-box domain containing protein, expressed (L. Guo et al. 2021) | | |
| LVR_ST2_8 | Migut.08G233700 | OsFBK19 - F-box domain and kelch repeat containing protein, expressed (L. Guo et al. 2021) | | |
| LVR_SEM1_7 | Migut.07G084200 | OsFBX433 - F-box domain containing protein, expressed (F. Guo et al. 2021) | | |
| LVR_SEM2_8 | Migut.08G237000 | auxin-induced protein 5NG4, putative, expressed (Scofield et al. 2018) | | |
| LVR_SEM3_8 | Migut.08G237000 | auxin-induced protein 5NG4, putative, expressed (Scofield et al. 2018) | | |
| LVR_SN2 | Migut.02G172100 | OsGH3.5 - Probable indole-3-acetic acid-amido synthetase, expressed (Luo et al. 2023) | | |
| LVR_SB8 | Migut.08G242800 | Calcium-dependent phospholipid-binding Copine family protein (Jing et al. 2024) | Migut.08G243000 | CAMK_CAMK_like.43 - CAMK includes calcium/calmodulin depedent protein kinases, expressed (Tripathi et al. 2009) |

| | | | | |
|---|---|---|---|---|
| LVR_SBA8 | Migut.08G238300 | Leucine-rich repeat (LRR) family protein (DeYoung et al. 2006; Kang et al. 2017) | | |
| LVR_RLL2 | Migut.02G118800 | carotenoid cleavage dioxygenase 1 (Auldridge et al. 2006) | | |
| LVR_RLW2 | Migut.02G118900 | CCCH-type zinc finger protein with ARM repeat domain (Kong et al. 2006) | | |
| CWF_FN1 | Migut.01G000800 | CHD3-type chromatin-remodeling factor PICKLE, putative, expressed (Fu et al. 2016; Jing et al. 2019; Yoon et al. 2021) | | |
| CWF_FN6 | Migut.06G188300 | WUSCHEL related homeobox 2 (Van Der Graaff et al. 2009) | | |
| CWF_FN14 | Migut.14G299900 | GASR6 - Gibberellin-regulated GASA/GAST/Snakin family protein precursor, expressed (Roxrud et al. 2007; Bouteraa et al. 2023; Chen et al. 2024) | Migut.14G300800 | COP1 |
| CWF_CTL3 | Migut.03G101400 | GASR3 - Gibberellin-regulated GASA/GAST/Snakin family protein precursor, expressed (Muhammad et al. 2019; Bouteraa et al. 2023) | | |
| CWF_CTL12 | Migut.12G168200 | GRAS family transcription factor (Muhammad et al. 2019; Bouteraa et al. 2023) | | |
| CWF_CLW9 | Migut.09G105000 | OsIAA18 - Auxin-responsive Aux/IAA gene family member, expressed (Talbert et al. 1995; Zhong and Ye 1999; Cao et al. 2024) | Migut.13G002600<br>Migut.13G009600 | GRAS family transcription factor (Jin et al. 2025) |

| | | | | |
|---|---|---|---|---|
| CWF_FT6 | Migut.06G177100 | osFTL2 FT-Like2 homologous to Flowering Locus T gene; contains Pfam profile PF01161: Phosphatidylethanol amine-binding protein, expressed (Kojima et al. 2002; Hayama et al. 2003; Abe et al. 2005; Komiya et al. 2008) | Migut.06G177600 | AP2-like ethylene-responsive transcription factor PLETHORA 2, putative, expressed (Aukerman and Sakai 2003; Chen 2004; Yant et al. 2010) |
| CWF_FT14 | Migut.14G297800 | histone H3, putative, expressed (Tamada et al. 2009; Zhang et al. 2021; Li et al. 2025) | Migut.14G300800 | COP1-interacting protein 8 (Hardtke et al. 2002; Sarid-Krebs et al. 2015; Xu et al. 2016; Lee et al. 2017) |
| CWF_IW6 | Migut.06G200300 | vacuolar ATP synthase subunit H, putative, expressed (Schumacher et al. 1999; Fukao and Ferjani 2011) | | |
| CWF_ST2_6 | Migut.06G177600 | AP2-like ethylene-responsive transcription factor PLETHORA 2, putative, expressed (Prasad et al. 2011) | | |
| CWF_ST3_14 | Migut.14G242900 | auxin response factor 18, putative, expressed (Lavenus et al. 2015; Huang et al. 2016; Kim et al. 2020) | | |
| CWF_SEM2_1 | Migut.01G000800 | CHD3-type chromatin-remodeling factor PICKLE, putative, expressed (Ma et al. 2015; Chen et al. 2018) | | |
| CWF_SW12 | Migut.12G086800 | WRKY117, expressed (Wang et al. 2010; Li et al. 2015) | Migut.12G084500 | BRASSINOSTEROID INSENSITIVE 1-associated receptor kinase 1 precursor (Nam and Li 2002; Zhiponova et al. 2013) |
| CWF_RH3_2 | Migut.02G162200 | K12862 - pleiotropic regulator 1 (PLRG1, PRL1, PRP46) (Flores-Perez et al. 2010) | Migut.02G173500 | DEAD/DEAH box helicase, putative, expressed (Zhang et al. 2022) |
| IMPO_SB9 | Migut.09G000800 | TCP family transcription factor, putative, expressed | | |

| | | |
|---|---|---|
| | | (Takeda et al. 2003; Aguilar-Martínez et al. 2007; Gastaldi et al. 2024) |
| IMPO_SB13 | NA | NA |
