## supplemental figures for "Unique genetic bases of repeated life-history divergence associated with high altitude adaptation in *Mimulus* perennials"

**Fig. S1. Phenotypic distributions for each life-history trait measured in the four mapping populations.** A) LVR × OPB population. B) CWF × OPB population. C) EAM × OPB population. D) IMPO × OPB population. Vertical lines indicate the phenotypic means of the LVR, CWF, EAM, and IMPO (brown, solid), OPB (green, solid) and $F_1$s (blue, dashed) in each population. Significant deviation from normality as determined by the Shapiro-Wilks test are indicated by asterisks (* < 0.05, **< 0.01, *** < 0.001).

A

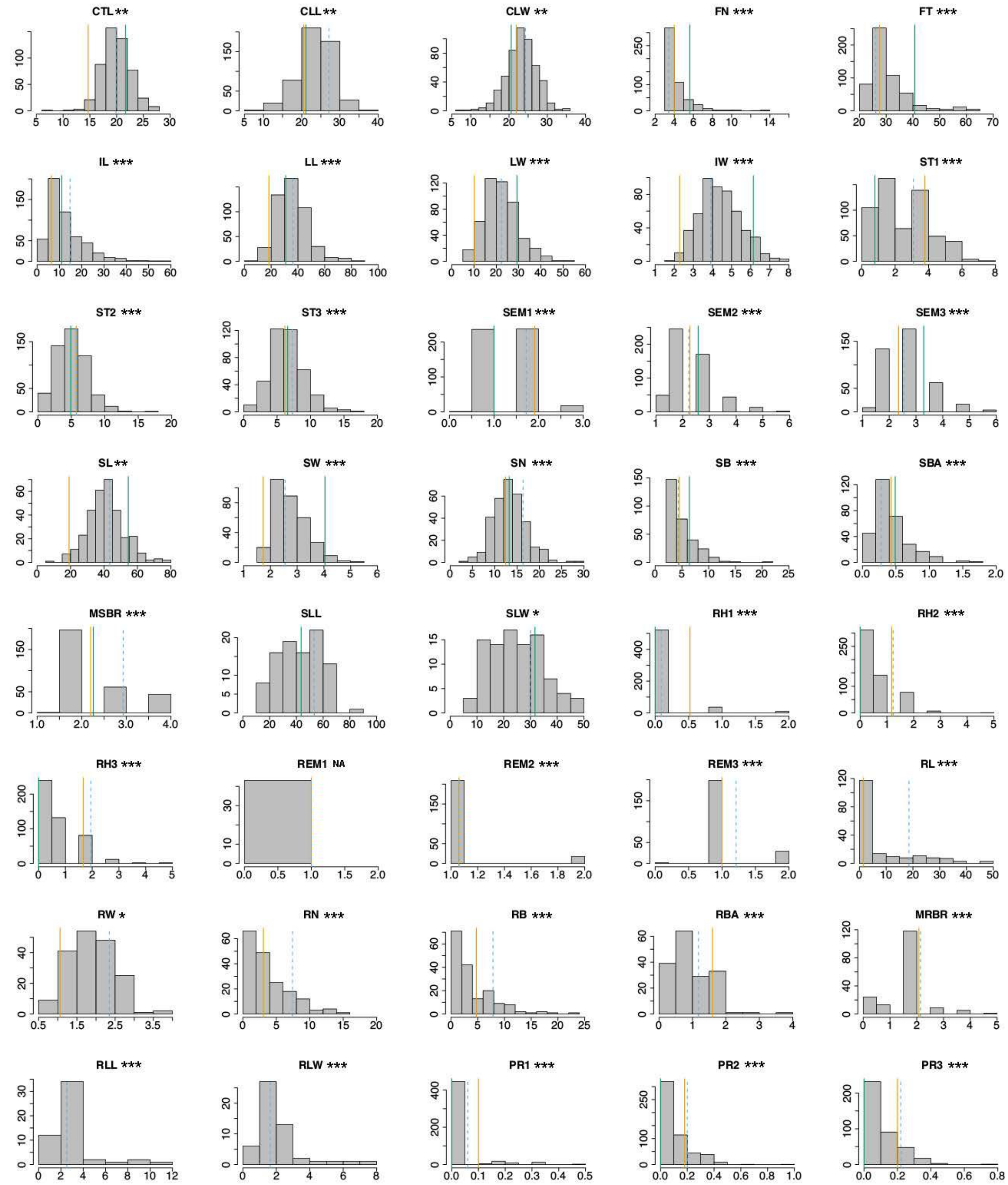


B

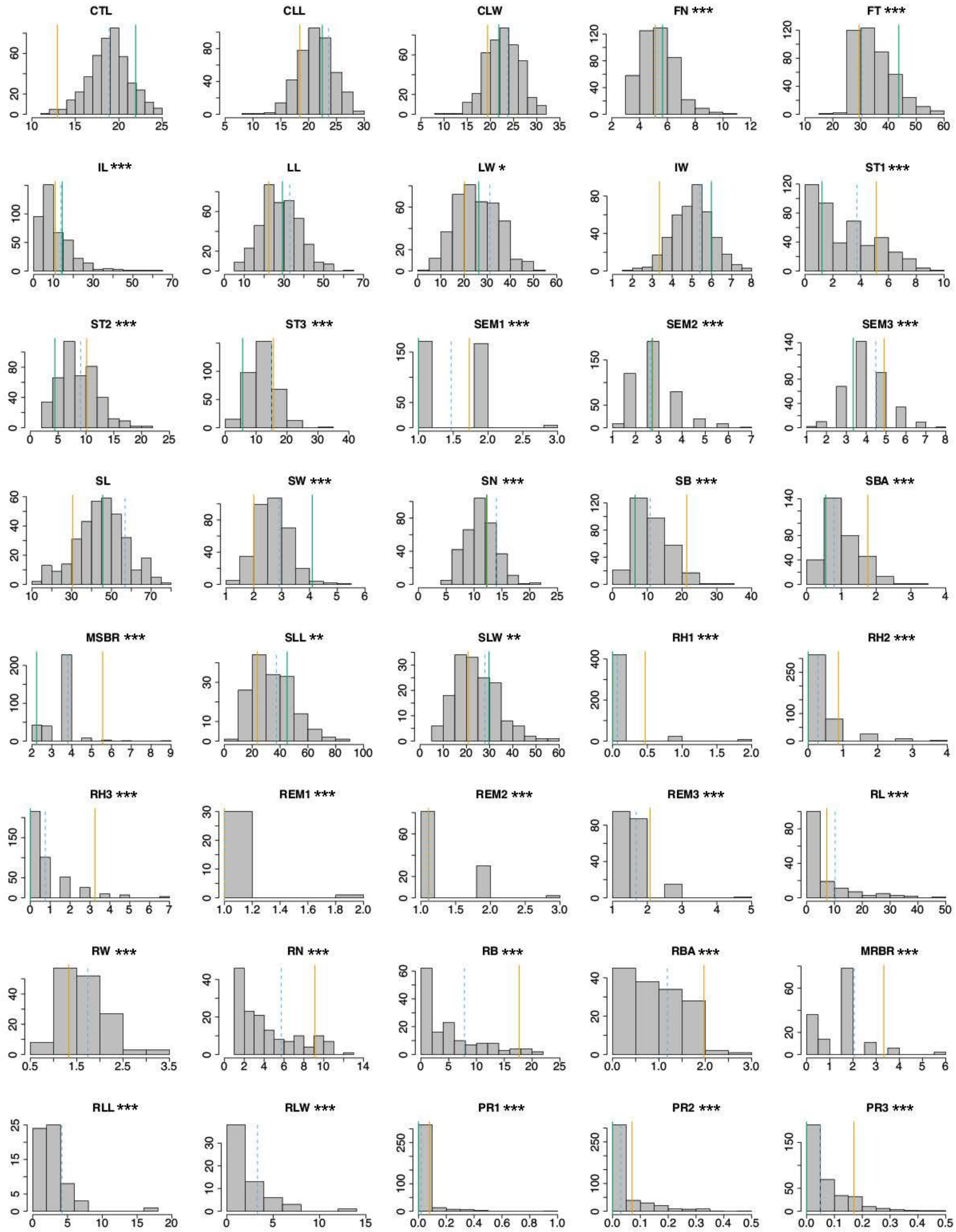
CTL
CLL
CLW
FN ***
FT ***
IL ***
LL
LW *
IW
ST1 ***
ST2 ***
ST3 ***
SEM1 ***
SEM2 ***
SEM3 ***
SL
SW ***
SN ***
SB ***
SBA ***
MSBR ***
SLL **
SLW **
RH1 ***
RH2 ***
RH3 ***
REM1 ***
REM2 ***
REM3 ***
RL ***
RW ***
RN ***
RB ***
RBA ***
MRBR ***
RLL ***
RLW ***
PR1 ***
PR2 ***
PR3 ***

C

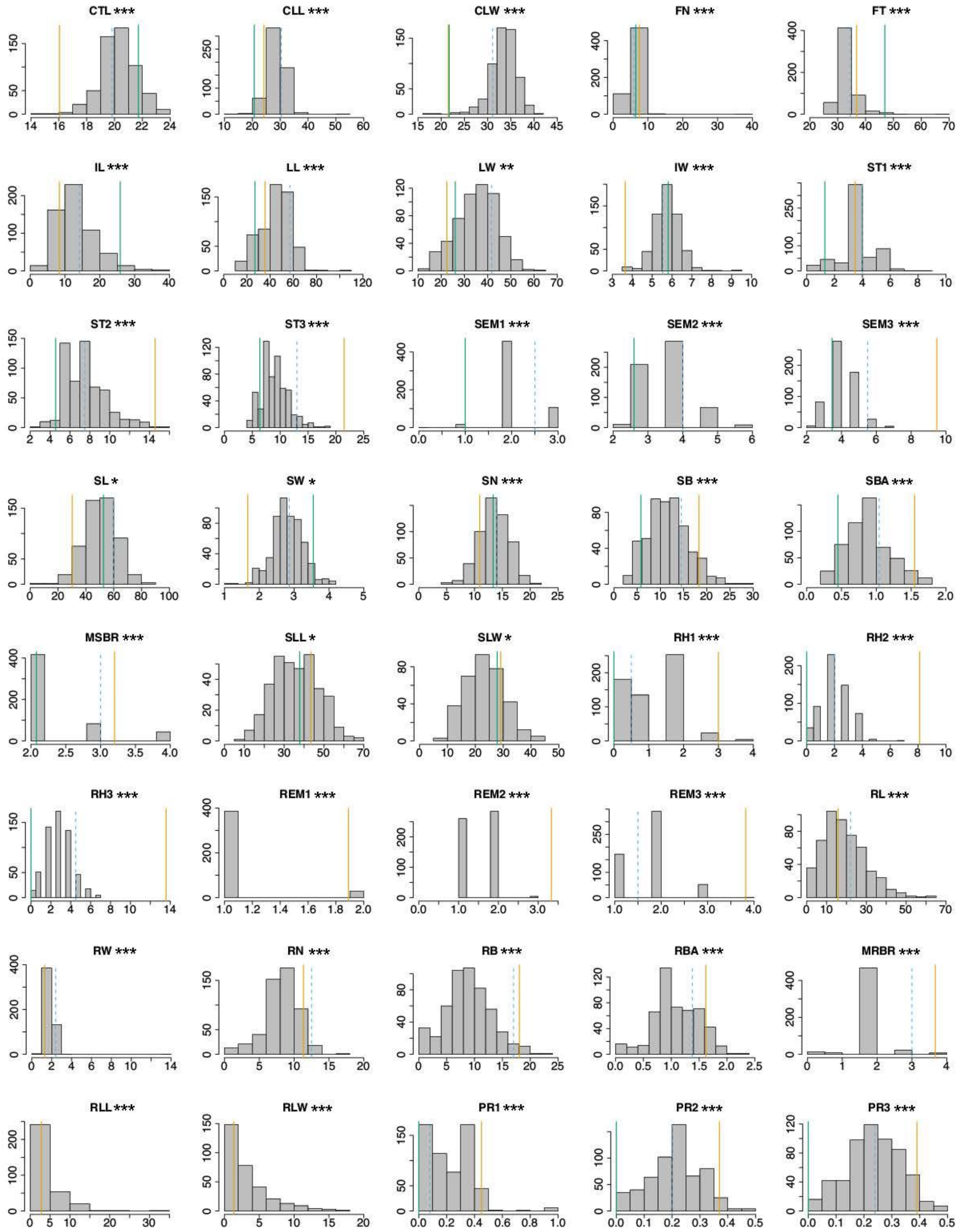
CTL ***
CLL ***
CLW ***
FN ***
FT ***
IL ***
LL ***
LW **
IW ***
ST1 ***
ST2 ***
ST3 ***
SEM1 ***
SEM2 ***
SEM3 ***
SL *
SW *
SN ***
SB ***
SBA ***
MSBR ***
SLL *
SLW *
RH1 ***
RH2 ***
RH3 ***
REM1 ***
REM2 ***
REM3 ***
RL ***
RW ***
RN ***
RB ***
RBA ***
MRBR ***
RLL ***
RLW ***
PR1 ***
PR2 ***
PR3 ***

D

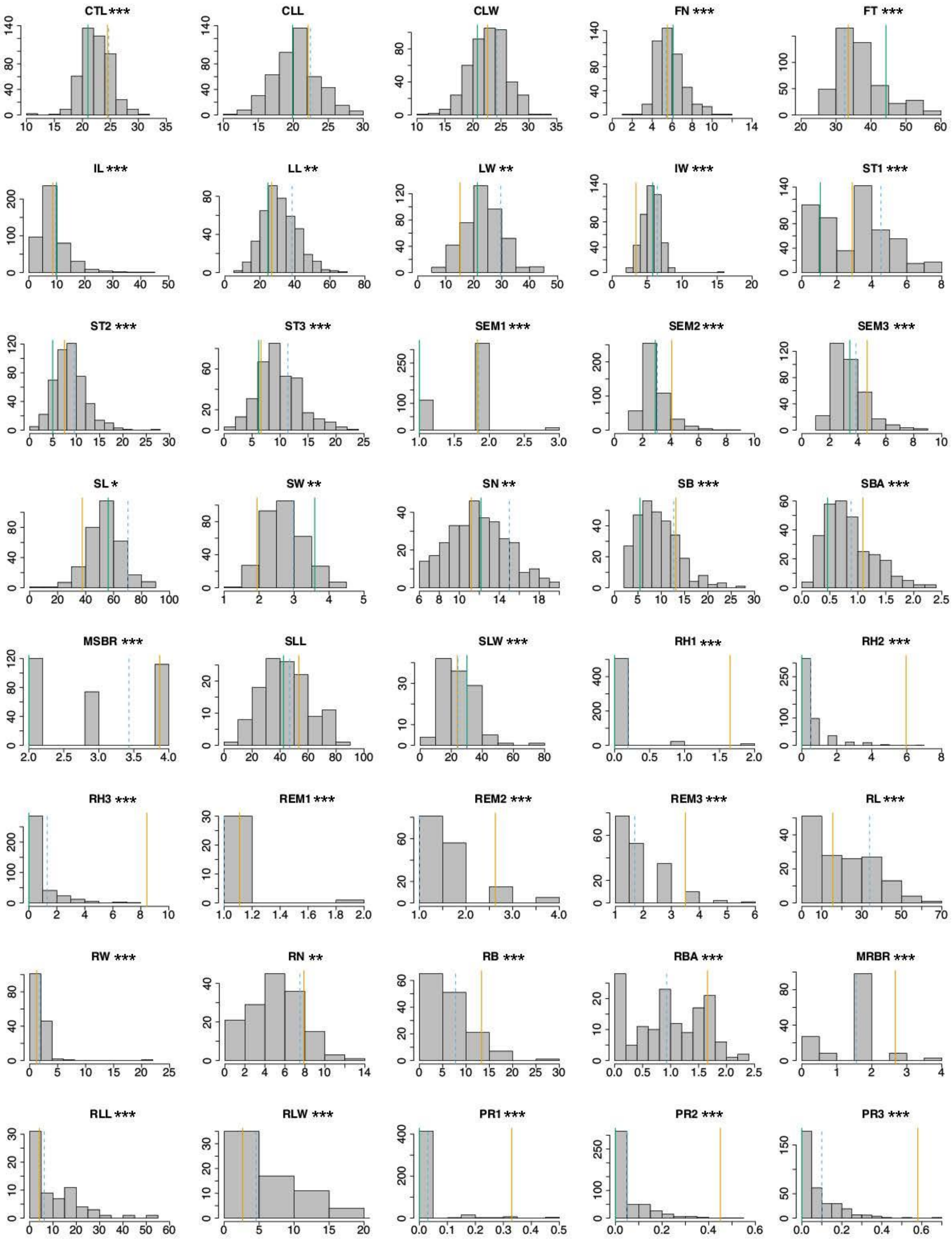
CTL ***
CLL
CLW
FN ***
FT ***
IL ***
LL **
LW **
IW ***
ST1 ***
ST2 ***
ST3 ***
SEM1 ***
SEM2 ***
SEM3 ***
SL *
SW **
SN **
SB ***
SBA ***
MSBR ***
SLL
SLW ***
RH1 ***
RH2 ***
RH3 ***
REM1 ***
REM2 ***
REM3 ***
RL ***
RW ***
RN **
RB ***
RBA ***
MRBR ***
RLL ***
RLW ***
PR1 ***
PR2 ***
PR3 ***

**Fig. S2. Phenotypic correlations among measured life-history traits in the four mapping populations.** Pairwise phenotypic correlations are shown for the LVR × OPB (A), EAM × OPB (B), IMPO × OPB (C), and CWF × OPB (D) populations.

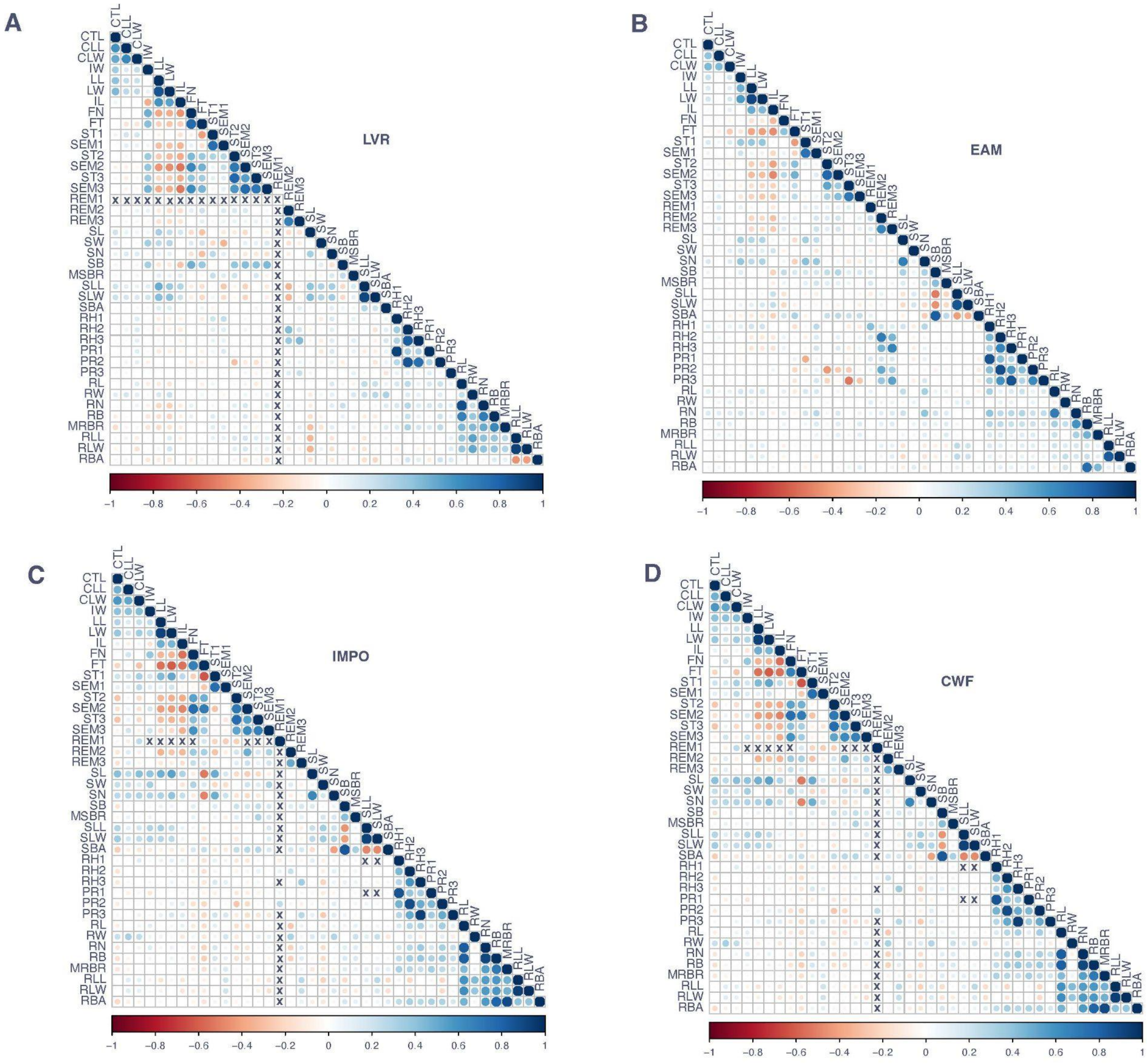

**Fig. S3. Genotype frequencies across fourteen chromosomes in the coastal *M. guttatus* × *M. corallinus* population.** Blue dots represent homozygous *M. corallinus* (EAM), brown dots represent homozygous coastal *M. guttatus* (OPB), and green dots represent heterozygous genotypes.

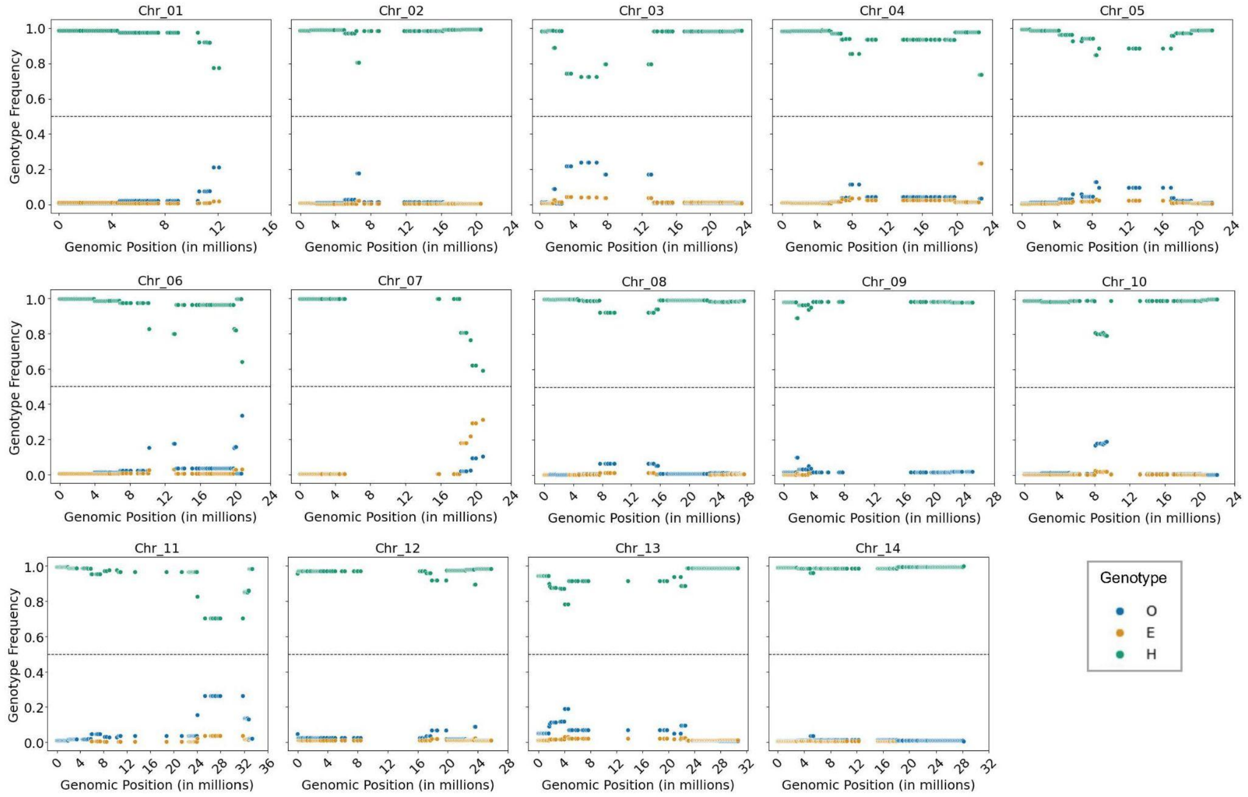

**Fig. S4. Density distributions of QTL effect sizes ($r^2$) for each trait module in each mapping population.**

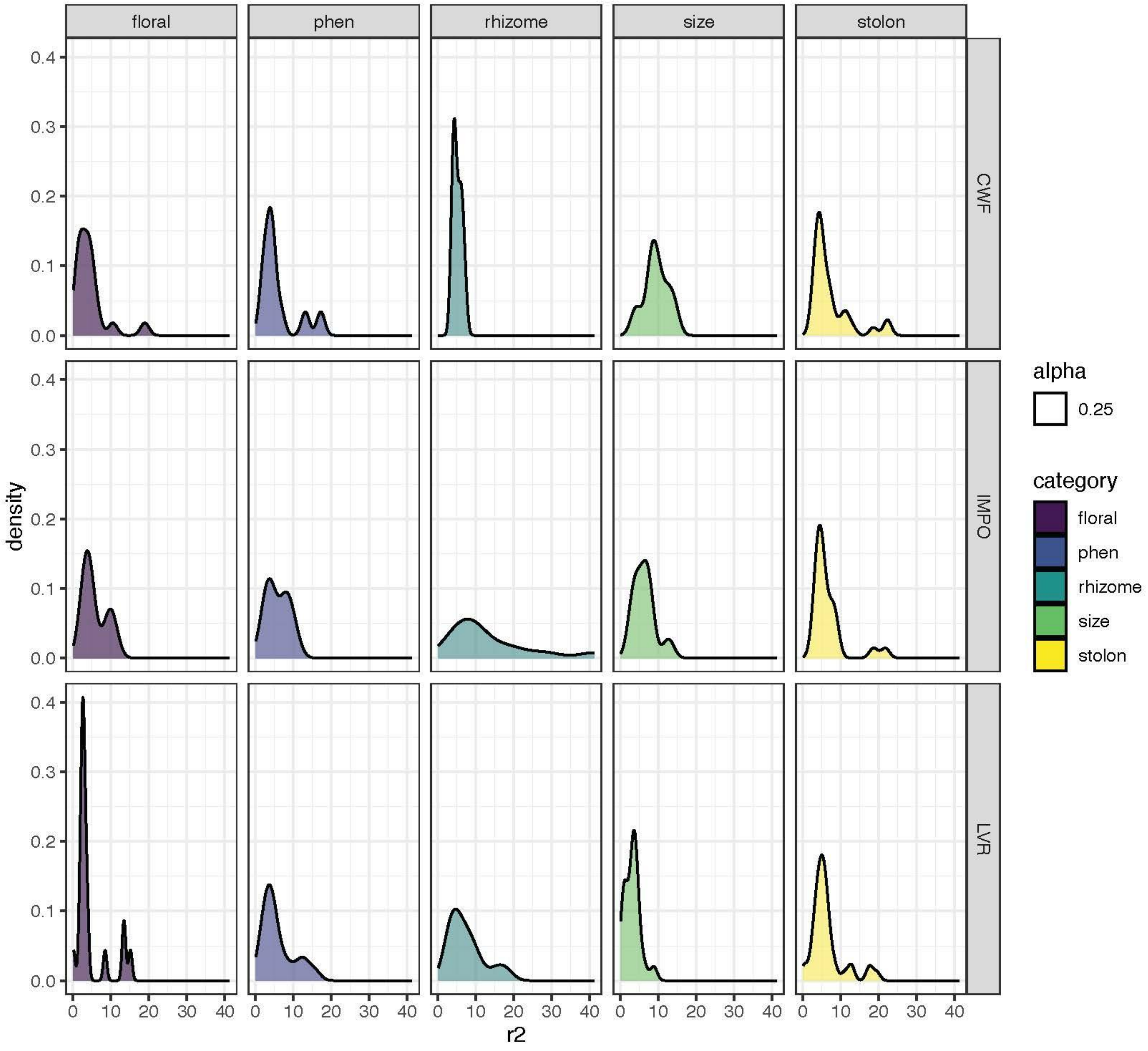


**Fig. S5. Relationship between the number of identified QTLs and effect size ($r^2$) for each trait in each mapping population**

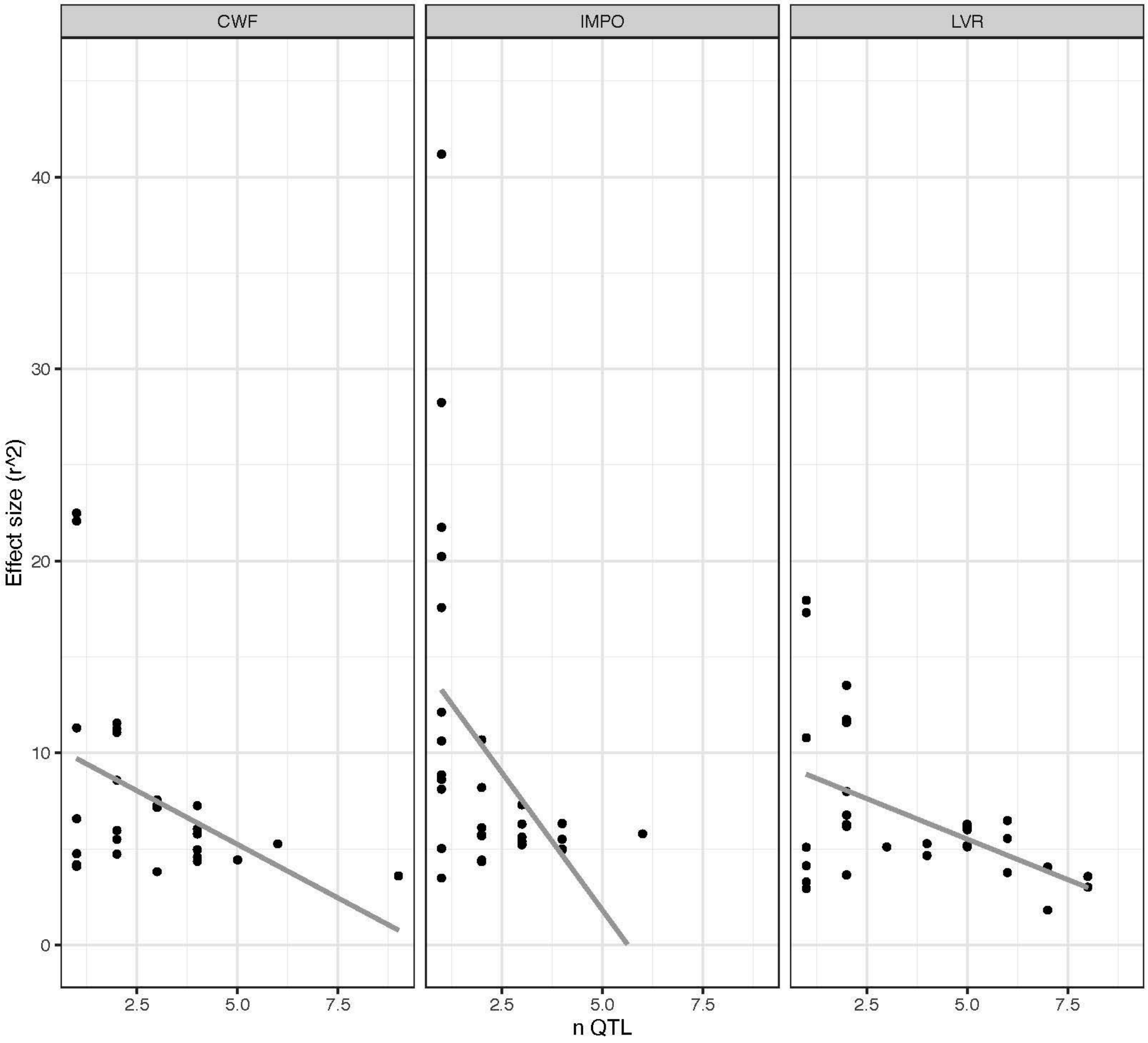


**Fig. S6. Phenotypic correlations between stolon number and flowering time for each of the four mapping populations.**

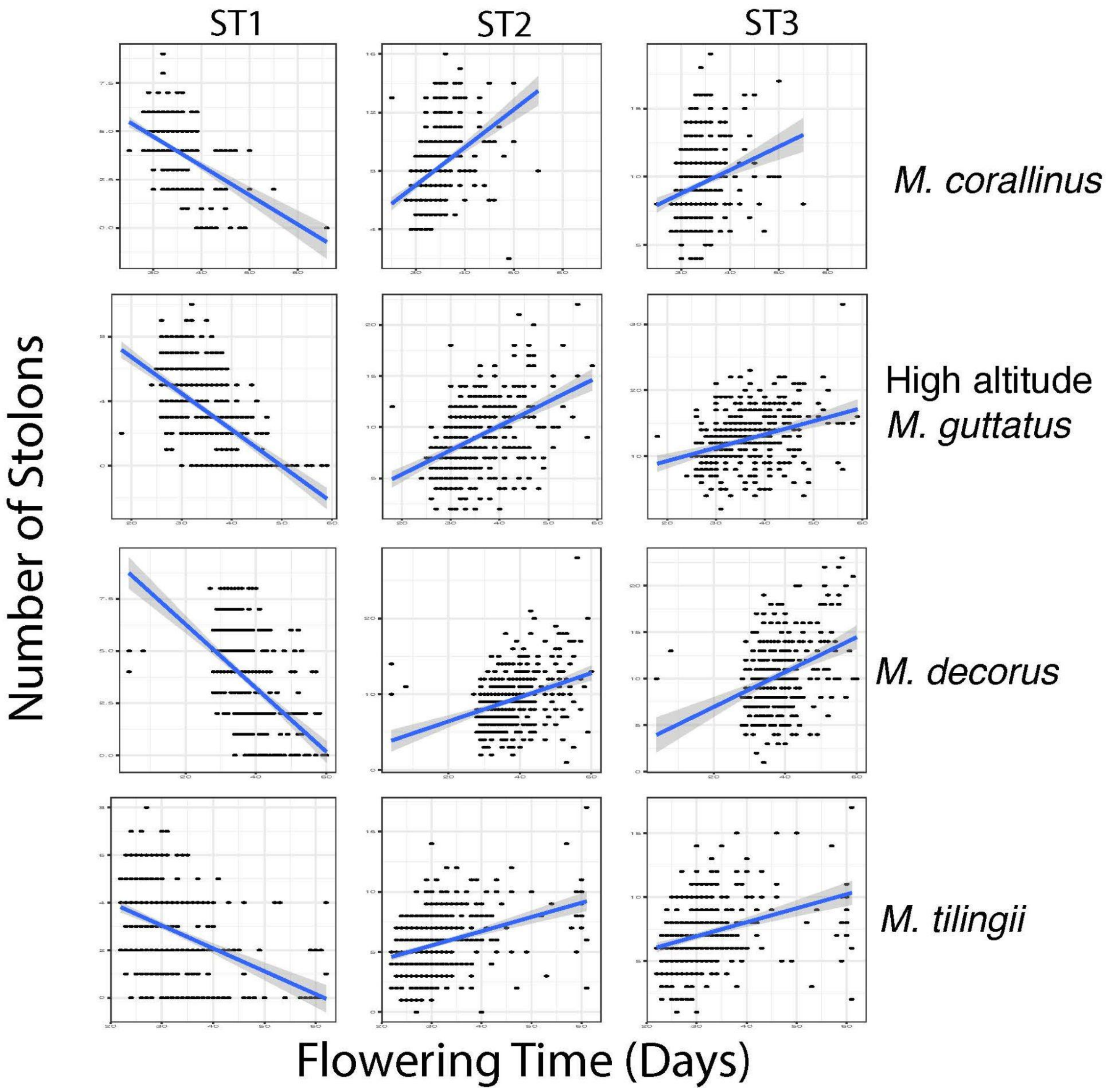


**Fig. S7. Major life-history traits measured in this study.** CLL, corolla limb length; CLW, corolla limb width; CTL, corolla tube length; IL, internode length; IW, internode width; LL, leaf length; LW, leaf width; SB, stolon branch; SN, stolon node; SW, stolon width; RL, rhizome length; RN, rhizome node; RB, rhizome branch; RW, rhizome width. Floral, phenological, and size-related traits are illustrated using representative individuals from various $F_2$ populations; stolon traits are represented by the coastal *M. guttatus* accession (OPB), and rhizome traits by CWF.

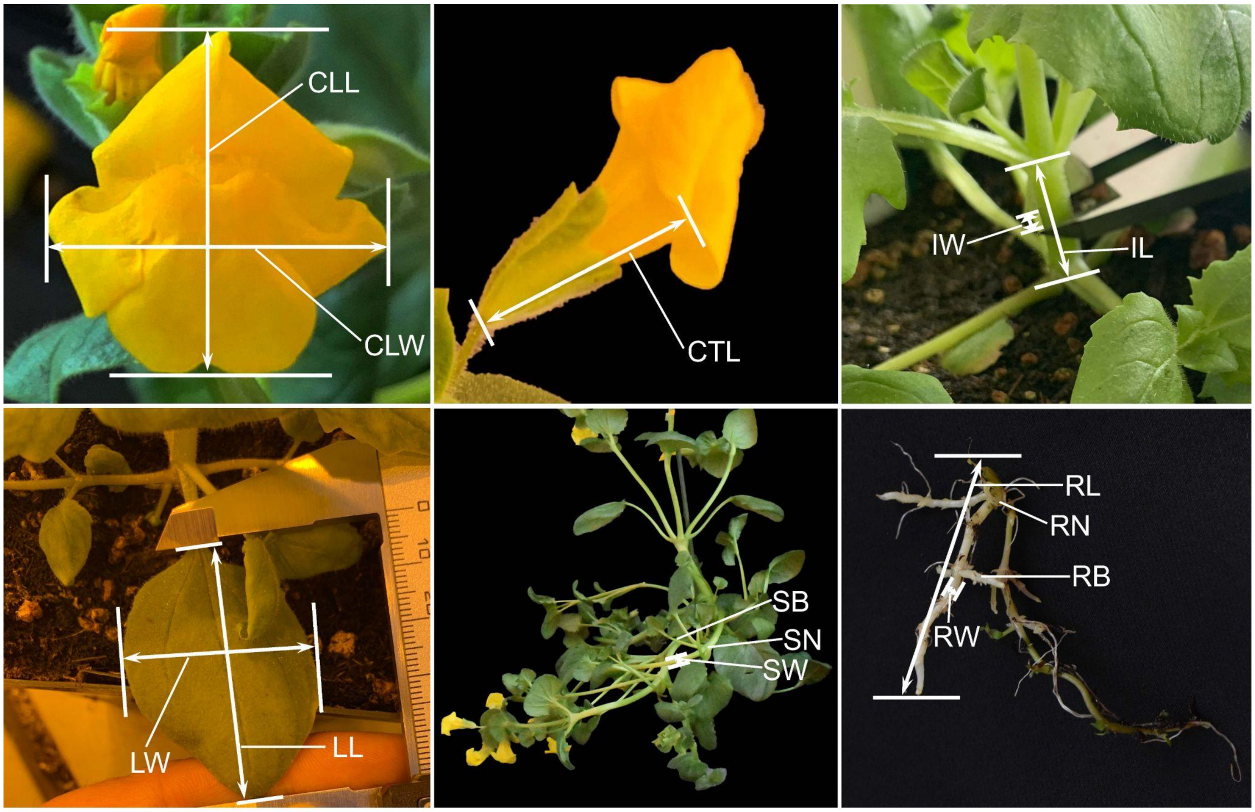


**Fig. S8. Linkage maps for the four mapping populations.** A) The coastal *M. guttatus* × *M. tilingii* population. B) The coastal *M. guttatus* × CWF population. C) The coastal *M. guttatus* × *M. corallinus* population. D) The coastal *M. guttatus* × *M. decorus* population.

A

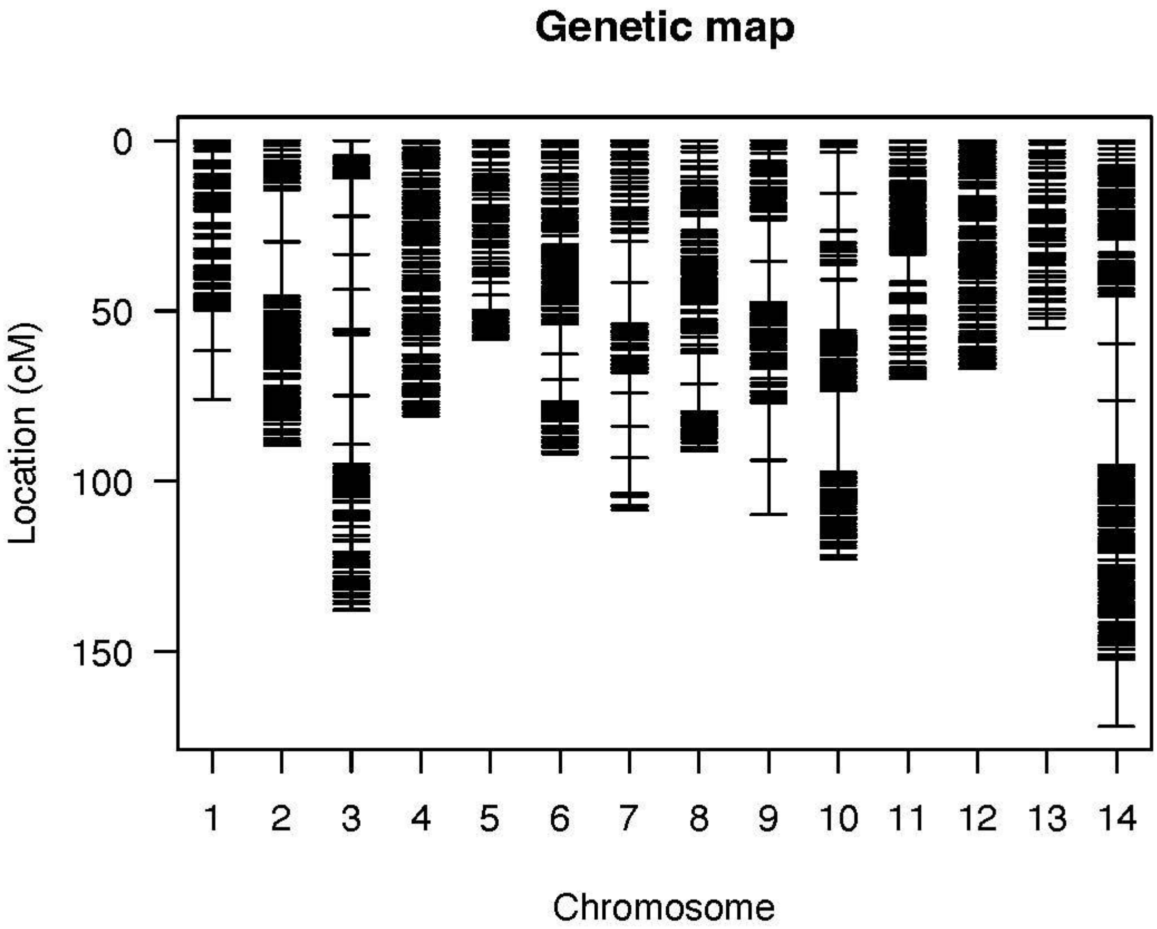
Genetic map
0
50
100
150
Location (cM)
1
2
3
4
5
6
7
8
9
10
11
12
13
14
Chromosome

B

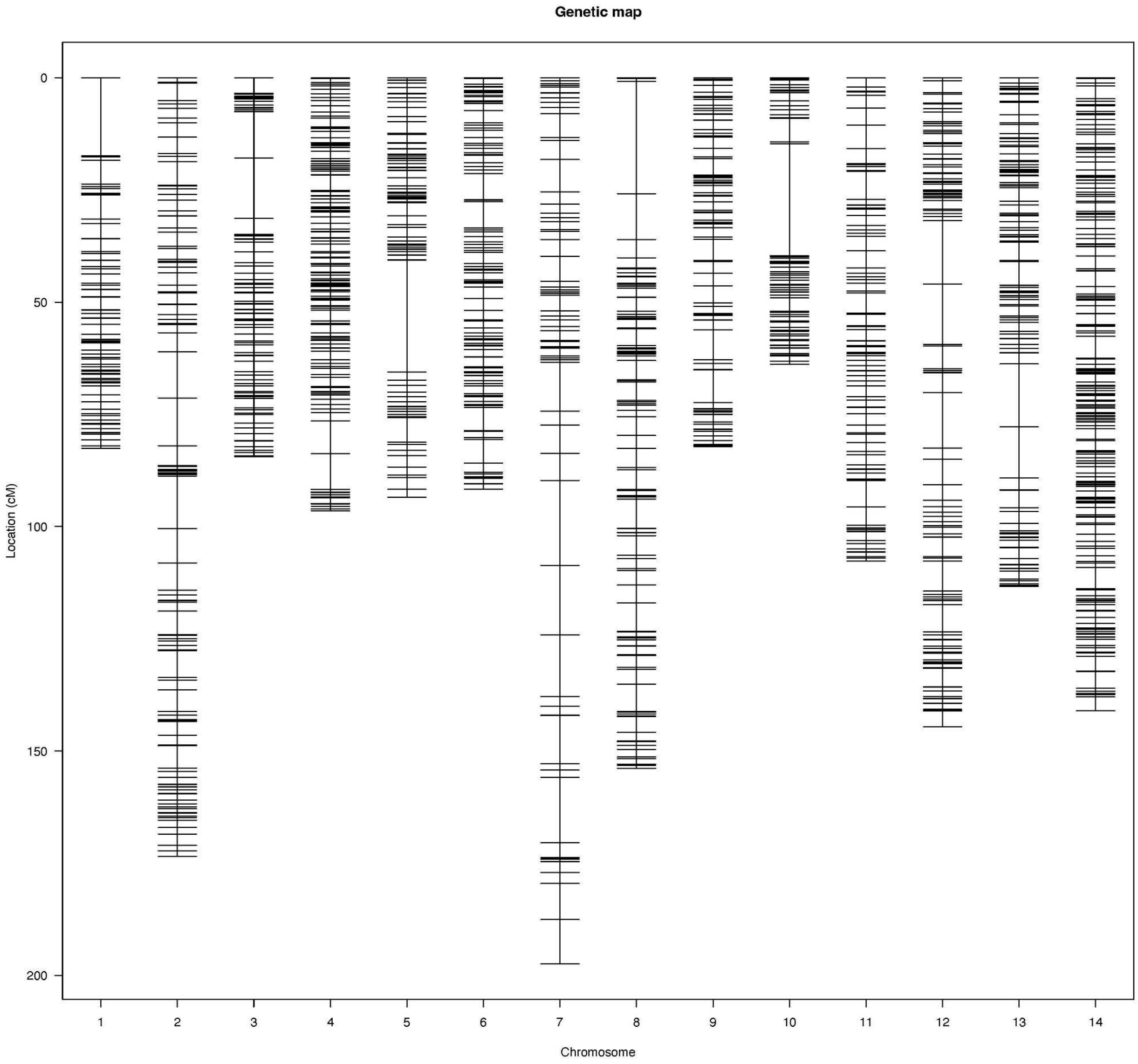

Genetic map
Location (cM)
0
50
100
150
200
Chromosome
1
2
3
4
5
6
7
8
9
10
11
12
13
14


C

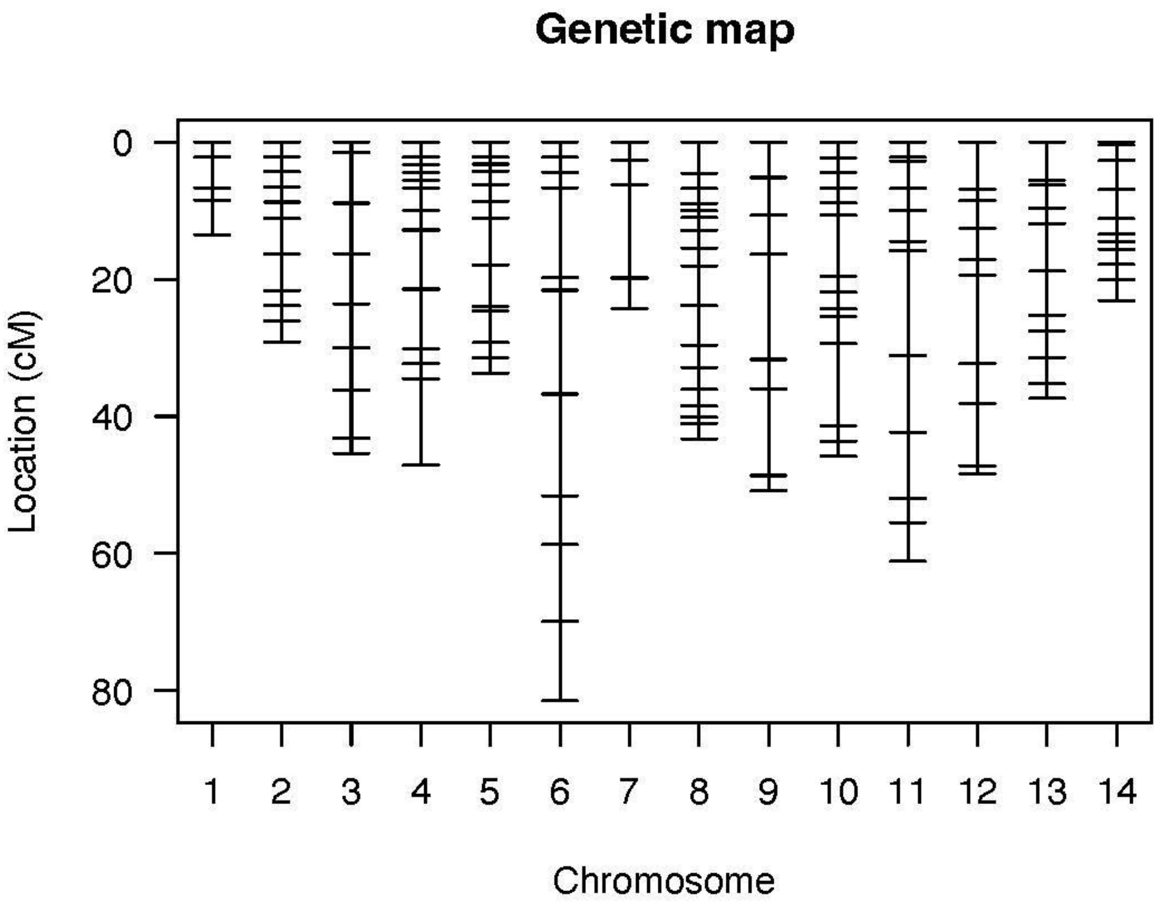
Genetic map
0
20
40
60
80
Location (cM)
1
2
3
4
5
6
7
8
9
10
11
12
13
14
Chromosome

D

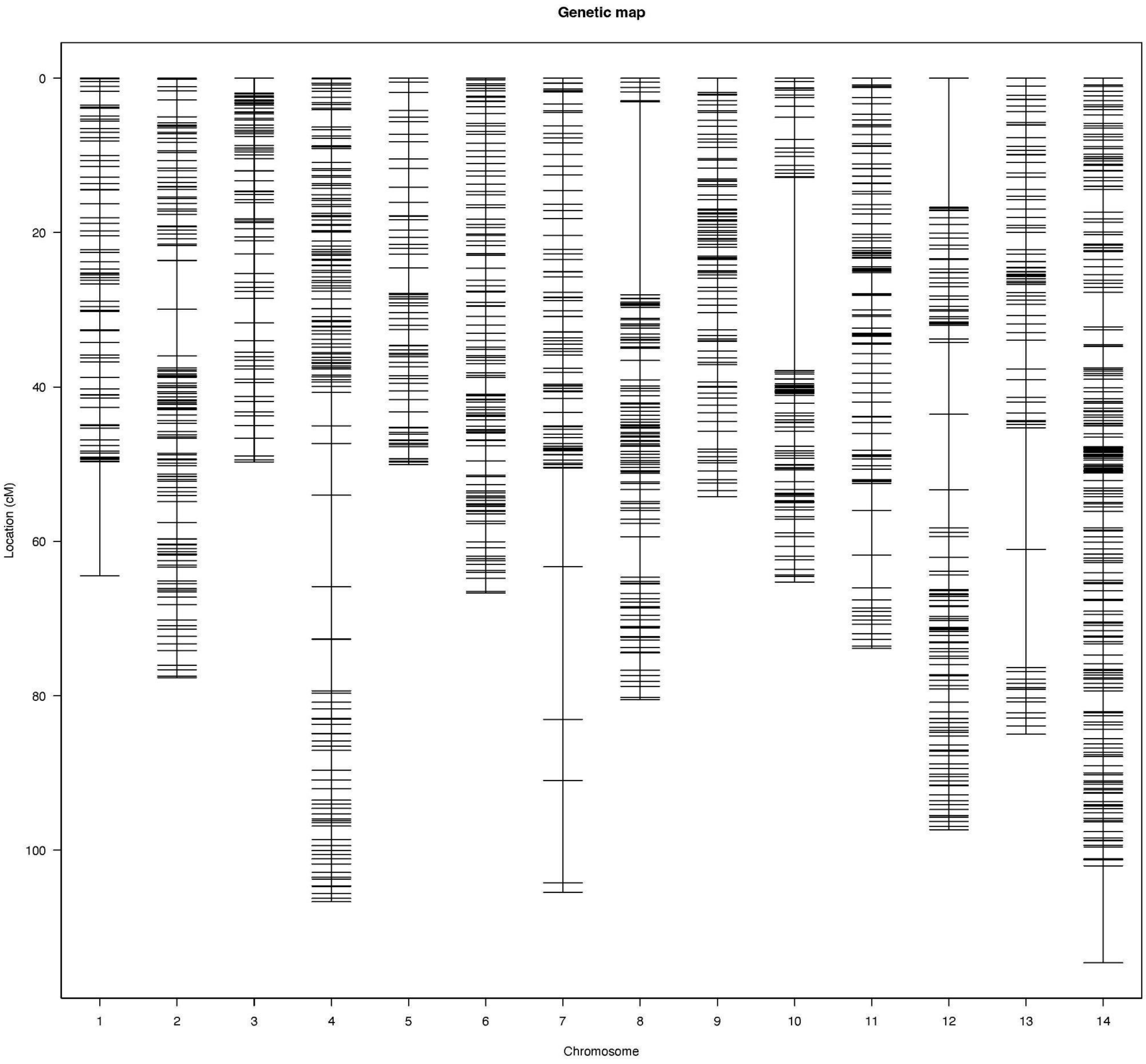
Genetic map
Location (cM)
0
20
40
60
80
100
1
2
3
4
5
6
7
8
9
10
11
12
13
14
Chromosome
